# Brain age prediction from structural and functional connectivity across the lifespan using connectome-based predictive modeling

**DOI:** 10.64898/2026.09.21.753233

**Authors:** Marie C McCusker, Javid Dadashkarimi, Huili Sun, Matthew Rosenblatt, Wei Dai, Dustin Scheinost

## Abstract

Brain maturation and aging are characterized by widespread changes in structural and functional brain organization. The extent to which these modalities capture shared versus distinct signatures of lifespan development and aging remains unclear. In this study, we applied connectome-based predictive modeling (CPM) to connectomes derived from diffusion MRI and resting-state functional MRI in Human Connectome Project Development, Young Adult, Aging, and combined Lifespan cohorts to predict chronological age. Both structural and functional connectomes significantly predicted age across cohorts, demonstrating that age-related information is distributed throughout whole-brain connectivity patterns. Prediction performance varied across lifespan stages, with generally stronger performance in the Development, Aging, and Lifespan cohorts than in Young Adulthood. Structural and functional models produced significantly correlated predicted ages in several cohorts, indicating shared age-related information across modalities. Structural-functional convergence analyses revealed limited and variable correspondence between modality-specific predictive features, suggesting that structural and functional connectomes capture both shared and complementary aspects of brain maturation and aging. Network-level analyses demonstrated that age-predictive information was distributed across canonical brain networks, including cerebellar, frontoparietal, salience, somatomotor, and subcortical networks, with patterns varying across cohorts and modalities. Multimodal CPM improved prediction relative to unimodal models in several cohorts, supporting the complementary contribution of structural and functional connectivity to age prediction. Cross-cohort, cross-sex, and cross-modality analyses further characterized the generalizability and shared information of predictive models across lifespan stages, sexes, and imaging modalities. This work demonstrates the utility of CPM for characterizing distributed structural and functional connectivity patterns associated with chronological age across the lifespan and provides a foundation for future studies of individual variability in brain maturation and aging.

## Introduction

Brain age models estimate an individual’s chronological age from neuroimaging data using machine learning (Cole et al., 2019; Cole and Franke, 2017; Franke et al., 2010) and are used to characterize age-related variation in the brain. Deviations between predicted and chronological age, commonly termed the brain age gap estimate, have been proposed as markers of individual differences in brain development and aging. These brain age gaps correlate with a wide range of phenotypic information and may have clinical relevance for neurological and psychiatric conditions (Anatürk et al., 2021; Bashyam et al., 2020; Cole and Franke, 2017; Franke and Gaser, 2019; Zhang et al., 2025). Consequently, brain age prediction is increasingly used to study individual differences in brain development, aging, and disease.

Many neuroimaging modalities have been used to predict brain age (Baecker et al., 2021; Soumya Kumari and Sundarrajan, 2024). Brain age models based on structural and functional features have been developed over different periods of development and adulthood. Most have focused on morphometric measures derived from structural MRI, including cortical thickness, gray matter volume, and other regional measures (Bashyam et al., 2020; Cole et al., 2017; Franke et al., 2012, 2010; Madan and Kensinger, 2018). White matter features have also been used to characterize age-related variation and predict age across development and adulthood (Kopetzky et al., 2024; Mwangi et al., 2013; Yap et al., 2013). Recent studies have examined functional MRI measures, such as resting-state functional connectivity (Chang et al., 2024; Li et al., 2018).

One approach for predicting age from connectomes is connectome-based predictive modeling (CPM; Shen et al., 2017). CPM is a data-driven framework that uses whole-brain connectivity profiles to predict behavior, cognitive, or demographic variables. It identifies connectivity features associated with a target phenotype and uses them to generate predictions for previously unseen individuals. Compared to other machine-learning approaches, CPM offers direct interpretation of predictive brain networks (Gao et al., 2019; Shen et al., 2017). Nevertheless, its application to brain age has been narrower. For example, prior CPM brain-age studies have focused on predicting age from adult functional connectomes (Kim et al., 2022) and neonatal structural and functional connectomes (Sun et al., 2024).

While prior research has provided valuable insights into age prediction using structural and functional connectomes, gaps persist in the literature. First, many brain age studies focus on relatively narrow age ranges, including developmental or adult samples, limiting our understanding of how predictive performance and age-related connectivity patterns differ across development, young adulthood, and aging (Kim et al., 2022; Sun et al., 2024). Second, few studies have directly compared the predictive performance of structural and functional connectomes within the same participants and across distinct periods of the lifespan. Some studies have examined structural and functional information within the same participants (Guan et al., 2024; Liem et al., 2017), but it remains unclear whether structural and functional connectomes capture common or modality-specific connectivity signatures of age. Some studies have combined structural and functional features to determine whether a combined model performs differently than single-modality models (de Lange et al., 2020; Guan et al., 2024; Liem et al., 2017; Niu et al., 2020; Rokicki et al., 2021), while findings regarding the benefit of multimodal integration have been mixed. Connectome-based approaches have also demonstrated that combining information across connectomes can improve prediction relative to individual connectomes (Gao et al., 2019). Third, sex differences in age prediction models remain understudied in age prediction across the lifespan.

In this study, we used structural and functional connectomes from the Human Connectome Project (HCP) to investigate brain age prediction across the lifespan using connectome-based predictive modeling. The sample included approximately 1,500 healthy individuals spanning childhood through older adulthood.

We evaluated structural, functional, and multimodal models; characterized the network organization and structural-functional convergence of predictive connectivity patterns; and assessed model generalizability across age groups, imaging modalities, and sex. We hypothesized that both structural and functional connectomes would reliably predict chronological age, with higher prediction accuracy in the developmental and older-adult cohorts due to greater individual variability during these periods. We further hypothesized that multimodal integration would improve prediction while revealing complementary modality-specific and shared connectivity patterns.

## Methods

### Participants

This study used diffusion MRI (dMRI), resting-state functional MRI (rsfMRI), and demographic data of healthy individuals from three open-source Human Connectome Project (HCP) datasets: HCP-Young Adults (HCP-YA), HCP-Development (HCP-D), and HCP-Aging (HCP-A). Participants were required to have both dMRI and rsfMRI scans. The three resulting datasets were combined to create the HCP-Lifespan (HCP-LS) cohort. The final sample’s demographics are in **Table 1**. All the analyses were performed on this subset. Additionally, male and female groups did not differ significantly in age in HCP-D, HCP-A, or HCP-LS, but differed in HCP-YA (*t*-test, *p* < 0.05).

**Table 1.** Human Connectome Project Demographics. HCP-Development, HCP-Young Adult, and HCP-Aging datasets after matching dMRI and rsfMRI modalities and thresholding under 0.1 mm framewise displacement.

|  | *N* | Mean Age (SD; range) | Sex (M, F) | Welch t-test<br>*p*-value |
| --- | --- | --- | --- | --- |
| **HCP-Development (D)** | 458 | 15.1 years (3.8; 8.1-21.9) | 210, 248 | 0.62484 |
| **HCP-Young Adult (YA)** | 723 | 28.4 years (3.7; 22-37) | 355, 368 | 8.34E-09 |
| **HCP-Aging (A)** | 290 | 56.0 years (13.3; 36-100) | 119, 171 | 0.202009 |
| **HCP-Lifespan (LS)** | 1471 | 29.7 years (15.8; 8.1-100) | 684, 787 | 0.085591 |
| HCP, Human Connectome Project; M, Males; F, Females |  |  |  |  |

### MRI Data Processing and Quality Control

Details about the HCP datasets and image protocols are provided in the corresponding references for HCP-D (Harms et al., 2018; Somerville et al., 2018), HCP-YA (Van Essen et al., 2013), and HCP-A (Bookheimer et al., 2019; Harms et al., 2018). Participants with excess head motion (> 0.1 mm mean framewise displacement) or poor-quality anatomical data were excluded. Mean framewise displacement was included as a covariate during rsfMRI feature selection. Data processing procedures are explained in the supplemental materials.

### Connectome Construction

Processed dMRI tractography data were parcellated using the Shen268-node atlas (Shen et al., 2013). Structural connectomes were constructed based on the mean quantitative anisotropy (QA) value of fibers connecting two regions (QA End). For rsfMRI, preprocessed rsfMRI BOLD time series were averaged within each Shen268-node atlas region, and functional connectivity was calculated as the Pearson correlation between node timeseries followed by Fisher’s z-transformation. Both modalities yielded 268 × 268 connectivity matrices per participant.

### Connectome-Based Predictive Modeling

Connectome-based predictive modeling (CPM; Shen et al., 2017) was used to predict chronological age from structural and functional connectivity separately within each age cohort. Models were evaluated using 10-fold cross-validation repeated across 100 iterations using predefined, identical fold assignments across modalities and CPM approaches. Within each iteration, participants were partitioned into training and testing sets such that each participant served as an independent test case once per fold. Within each training fold, edge-wise Pearson correlations were computed between connectivity strength and chronological age. When covariates were included, partial correlations were used. Edges significantly correlated with age (*p* < 0.05) were selected as predictive features. A linear regression model was then fit between network strength and age in the training data and applied to held-out test participants to predict age. Ridge regression CPM (Gao et al., 2019) was also performed as a robustness analysis, with detailed methods provided in the supplemental materials.

The same CPM procedure was applied to within- and between-network combinations using 10 predefined networks from the Shen268 atlas (Shen et al., 2013), and separately within male and female groups to examine sex differences. Detailed network and sex-stratified analyses are described in the supplemental materials.

### Model Performance Evaluation

CPM performance was evaluated separately by age cohort and imaging modality using Pearson’s correlation coefficient (*r*), Spearman’s rank correlation (*ρ*), mean squared error (MSE), mean absolute error (MAE), and cross-validation coefficient of determination (*q²*; Poldrack et al., 2020). Performance across the 100 iterations was summarized using the median-performing run, based on Pearson’s *r*.

Statistical significance was assessed using permutation-based null models following standard CPM procedures (Shen et al., 2017). Age labels were randomly shuffled while preserving the original connectomes and cross-validation fold assignments. Null distributions were generated using 10,000 permutations, and observed performance was compared with the null distribution using one-tailed nonparametric permutation tests. For correlation metrics, *p*-values were calculated as the proportion of null iterations with values equal to or greater than the observed value. For error metrics, the comparison direction was reversed. Benjamini-Hochberg false discovery rate (FDR) correction was applied within the relevant analyses, with a significance threshold of *p* < 0.05. Ridge regression CPM was evaluated as a robustness analysis (see supplemental materials).

### Exploratory Analyses of Model Characteristics and Bias

Exploratory analyses characterized shared information and prediction bias across structural and functional models. Within each age cohort, predicted ages were correlated across modalities, and brain age gap (BAG; predicted age − chronological age) was calculated and compared across modalities. Associations between BAG and chronological age, as well as between BAG and predicted age, were assessed to characterize age-dependent bias and error structure. These analyses were exploratory and were not used to assess predictive accuracy. Correlations were corrected for multiple comparisons using FDR, with *p* < 0.05 considered significant.

### Whole-Brain Predictive Networks

To characterize connectivity features contributing to age prediction, CPM edges significantly associated with age (*p* < 0.05) were identified separately for positive and negative associations within each training fold. Consensus feature sets were generated by retaining edges selected across all folds within a run (fold-wise intersection). Signed 268 × 268 predictive networks were constructed by assigning +1 and −1 to positive and negative predictive edges, respectively, and 0 to all other edges. The median-performing run, selected based on Pearson’s *r* between predicted and chronological age, was used as the representative model for each cohort, modality, and sex-stratified condition. Signed networks from these runs were retained for downstream analyses, and absolute-valued versions were generated to quantify edge magnitude independent of direction.

### Structural-Functional Predictive Pattern Convergence

To assess convergence between structural and functional predictive connectomes, median-performing consensus CPM networks were compared separately within each age cohort. Binary adjacency matrices representing positive, negative, and absolute predictive edges were generated using the Shen268 parcellation (Shen et al., 2013). Convergence was assessed at three levels: edge, node, and network.

At the edge level, structural and functional CPM networks were compared using binary edge representations. The upper triangular portion of each 268 × 268 matrices were vectorized, excluding self-connections, and compared using Pearson correlation. To evaluate whether shared predictive edges exceeded chance expectations, a hypergeometric cumulative density test was performed. The number of overlapping predictive edges between structural and functional networks was compared against the expected overlap given the total number of possible edges and the number of predictive edges identified by each modality.

At the node level, node degree was calculated as the number of predictive connections for each node, and structural-functional convergence was assessed using Pearson correlation between node degree vectors.

At the network level, predictive edge densities were calculated within and between predefined Shen networks (Shen et al., 2013). The upper triangles of the resulting network density matrices, excluding within-network densities, were compared using Pearson correlation.

Statistical significance at each level was assessed using 10,000 permutations by randomly shuffling the structural edge, node, or network representations while holding the corresponding functional representation fixed. Permutation-derived p-values were corrected for multiple comparisons using FDR within each level of analysis.

### Sex Differences in Prediction Performance

Sex differences in prediction performance were assessed by comparing the Pearson correlations from the independent male and female samples using Fisher’s *r*-to-*z* transformation with the corresponding male and female sample sizes. Two-sided *p*-values were calculated for each of the cohort-by-modality comparisons and corrected using the Benjamini-Hochberg FDR procedure with FDR-corrected *p* < 0.05 considered statistically significant.

### Model Generalizability Using Cross-Prediction

To assess model generalizability, the median-performing run was selected for each cohort and modality, and the corresponding data were used to refit a final CPM model. Positive and negative predictive edges were reselected using the original significance threshold, and the resulting signed network was used to calculate network strength as the sum of positive edge weights minus the sum of negative edge weights. A linear regression relating network strength to chronological age was fit, and the resulting network and regression coefficients were applied without retraining to independent datasets.

Cross-cohort analyses trained models in one cohort and tested them in the remaining cohorts while holding imaging modality constant, excluding HCP-LS self-comparisons to avoid leakage. Cross-modality analyses trained models using one modality and tested them on the other within the same cohort. Cross-sex analyses trained male-only and female-only models and tested them in the opposite sex within the same cohort and modality. Prediction performance was evaluated using Pearson *r* correlations.

Pearson correlation p-values were corrected using the Benjamini-Hochberg FDR procedure within each cross-prediction analysis.

### Multimodal CPM

To evaluate whether combining structural and functional connectivity improved age prediction, structural and functional connectomes were vectorized and concatenated into a single feature vector containing dMRI QA End edges followed by rsfMRI edges. A multimodal CPM model was trained separately within each age cohort using both modalities simultaneously. Unlike the unimodal rsfMRI CPM, no motion covariate was included because framewise displacement was specific to the rsfMRI acquisition and had no corresponding diffusion MRI measure. Prediction performance was evaluated using Pearson’s *r*, MAE, MSE, and permutation-based significance testing.

### Statistical comparison between multimodal and unimodal CPM performance

To evaluate whether multimodal CPM improved prediction performance relative to unimodal models, we performed paired iteration-wise comparisons across the 100 repetitions of 10-fold cross-validation.

Because multimodal and unimodal models were evaluated using identical train/test partitions, performance metrics were directly paired within each iteration. Differences between multimodal and corresponding unimodal models were calculated for Pearson correlation, MSE, and MAE. For Pearson correlation, positive differences indicated greater multimodal performance, whereas for MSE and MAE differences were calculated such that positive values indicated lower multimodal prediction error. The consistency of multimodal improvements across iterations was assessed using a one-sided exact binomial sign test, evaluating whether the proportion of iterations favoring the multimodal model exceeded chance expectation (50%). P-values were corrected for multiple comparisons using the Benjamini-Hochberg FDR procedure.

### Multimodal-Unimodal CPM similarity

A representative median-performing multimodal CPM model was selected using the same procedure as the unimodal models. Predictive edges were separated by modality to reconstruct structural (dMRI QA End) and functional (rsfMRI) signed consensus networks. Within each age cohort, modality-matched multimodal and unimodal consensus networks were compared for positive, negative, and absolute predictive edges at the edge, node, and network levels using the analyses described for structural-functional convergence.

## Results

### Structural and functional connectomes predict chronological age and cross-cohort prediction

Using structural connectomes, we observed robust age prediction in HCP-D (*r* = 0.57, FDR-corrected *p* < 0.0001; *MSE* = 9.61, *MAE* = 2.49), HCP-YA (*r* = 0.24, FDR-corrected *p* < 0.0001; *MSE* = 13.63, *MAE* = 3.02), HCP-A (*r* = 0.69, FDR-corrected *p* < 0.0001; *MSE* = 92.65, *MAE* = 7.58), and HCP-LS (*r* = 0.72, FDR-corrected *p* < 0.0001; *MSE* = 118.84, *MAE* = 8.60). Functional connectomes also showed strong performance, with HCP-D (*r* = 0.74, FDR-corrected *p* < 0.0001; *MSE* = 6.45, *MAE* = 2.02), HCP-YA (*r* = 0.23, FDR-corrected *p* < 0.0001; *MSE* = 13.24, *MAE* = 2.98,), HCP-A (*r* = 0.48, FDR-corrected *p* < 0.0001; *MSE* = 137.95, *MAE* = 9.47), and HCP-LS (*r* = 0.75, FDR-corrected *p* < 0.0001; *MSE* = 107.29, *MAE* = 7.55). Additional details about the performance results are in **Table S1**. These relationships between chronological age and predicted age from the CPM model are shown in **Figure 1A**. Generalization among the cohorts was weak and variable, with dMRI models producing small-to-moderate positive correlations and rsfMRI models not generalizing (**Figure 1B; Table S2**). All the rCPM results can be found in the supplement **Figures S3-S7** and **Tables S12-S23**.

**Figure 1.**
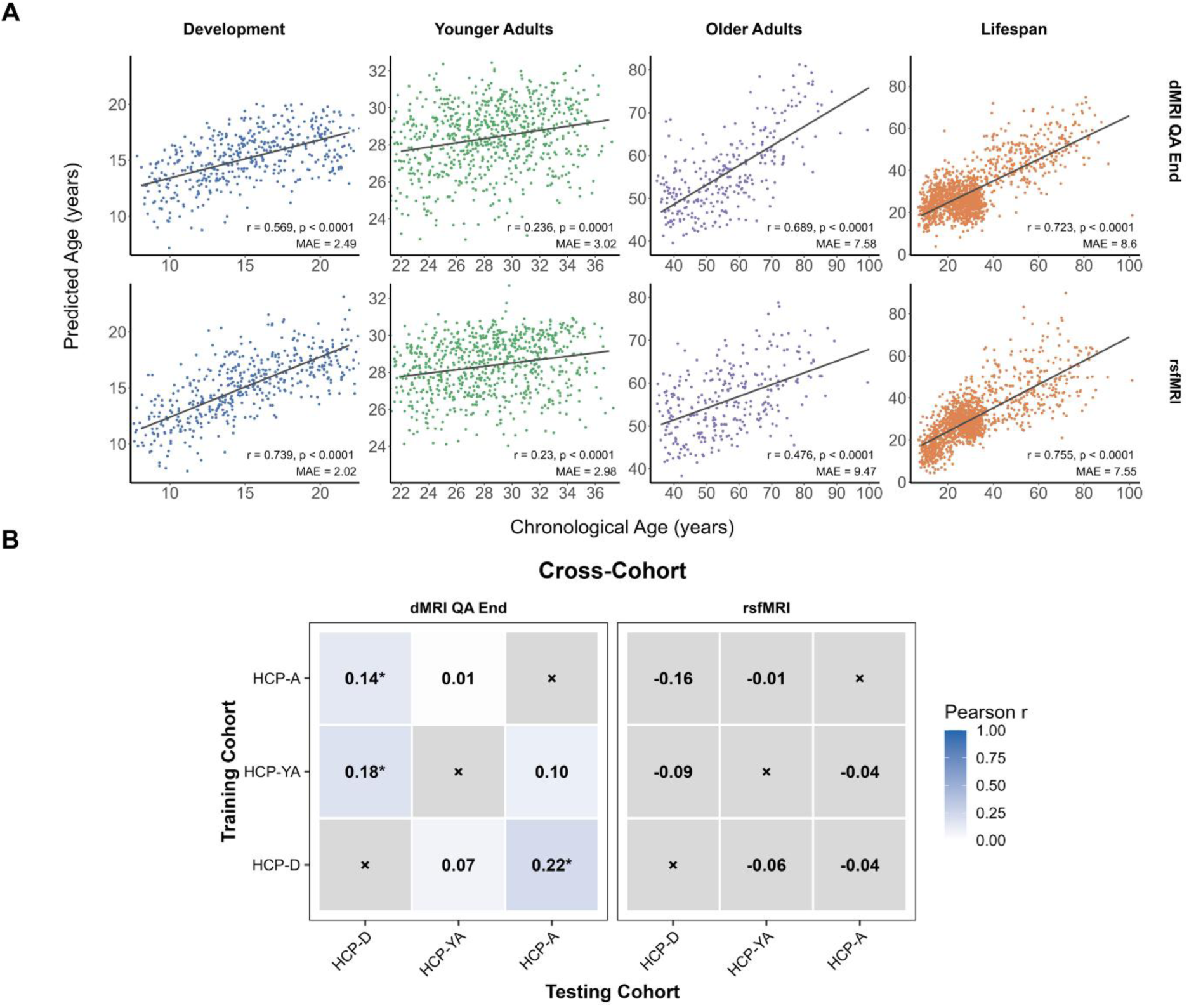
CPM brain age prediction performance across age cohorts. Scatterplots show predicted age versus chronological age for the median-performing CPM model in the developmental (HCP-D), young adult (HCP-YA), older adult (HCP-A), and lifespan (HCP-LS) cohorts. The top row shows structural connectivity models derived from dMRI quantitative anisotropy (QA End), and the bottom row shows resting-state functional connectivity (rsfMRI) models. Each point represents an individual participant. Solid gray lines indicate the linear regression fit between chronological and predicted age. Insets display the Pearson correlation coefficient (r) and mean absolute error (MAE) for the median-performing model; all prediction models remained significant following permutation testing (p < 0.0001). Points are colored by cohort: blue = developmental, green = young adults, purple = older adults, and orange = lifespan. (B) Heatmaps summarize the generalizability of final CPM age prediction models across cohorts evaluated through external validation/cross-sex prediction. Heatmap values represent the Pearson correlation coefficient (r) between predicted and chronological age when models were applied to independent datasets. Darker shading indicates stronger age prediction performance (higher Pearson r). Asterisks indicate predictions that remained significant after false discovery rate (FDR) correction (p < 0.05). Gray cells marked with × indicate comparisons that were not performed (e.g., identical training and testing cohort).

Across modalities, significant positive correlations between predicted age were observed in HCP-D (*r* = 0.5, FDR-corrected *p* < 0.001), HCP-A (*r* = 0.39, FDR-corrected *p* < 0.001), and HCP-LS (*r* = 0.56, FDR-corrected *p* < 0.001), indicating that individuals predicted to be older (or younger) by one modality also tended to be predicted as older (or younger) by the other modality (**Figure 2A**). No significant relationship was observed in HCP-YA. Similarly, across modalities, significant positive correlations between BAGS were observed across all cohorts (all *r*’s ≥ .49, FDR-corrected *p* < 0.001, **Figure 2B**), indicating that individuals with relatively larger positive or negative BAG values in one modality tended to show similar deviations in the other modality.

**Figure 2.**
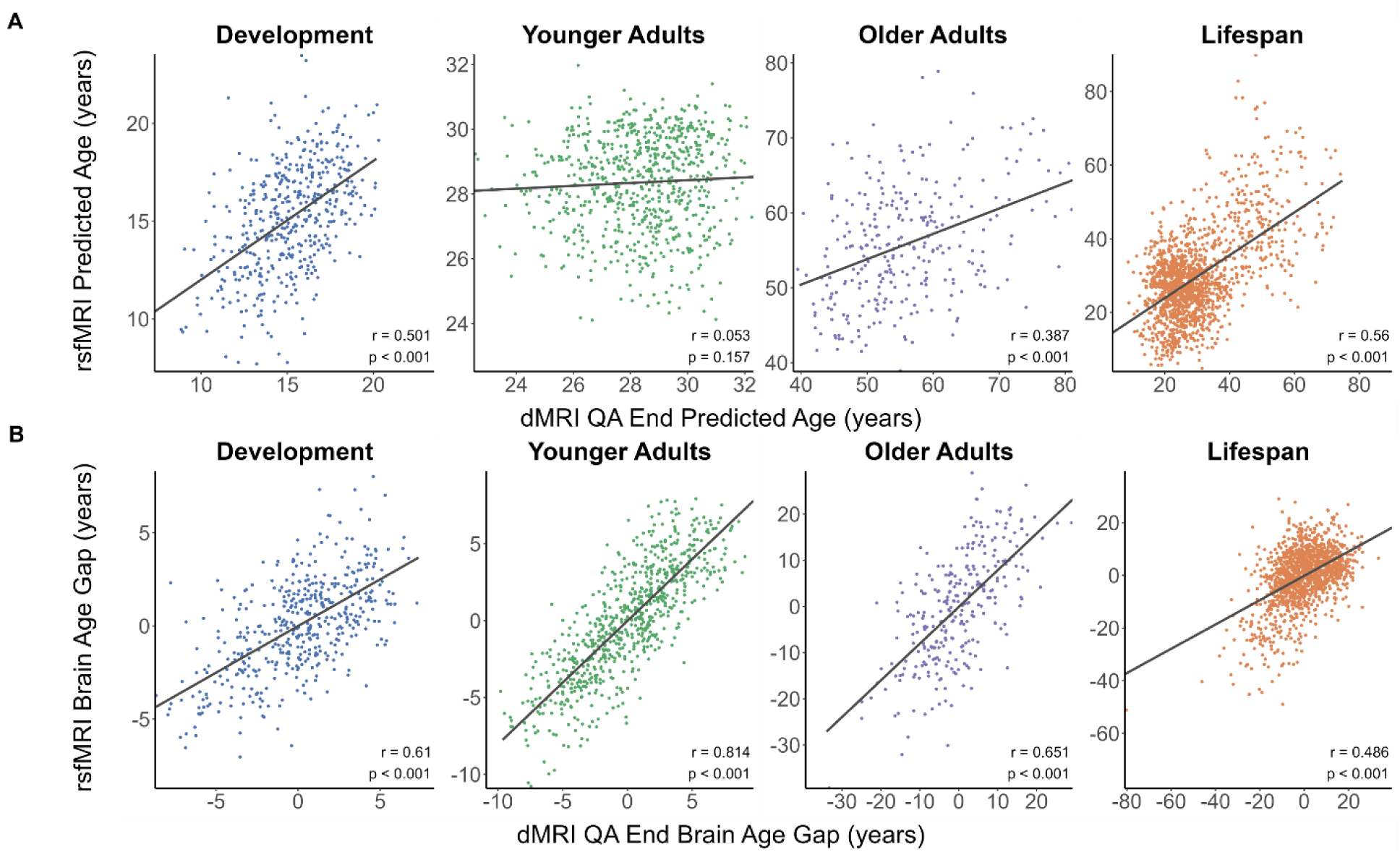
Correlations between predicted edges and brain age gaps. (A) Correlation between structural and functional CPM predicted ages. Scatterplots illustrate the relationship between predicted ages generated by the median-performing structural connectivity model (dMRI QA End; x-axis) and the median-performing rsfMRI model (y-axis) for the developmental (HCP-D), young adult (HCP-YA), older adult (HCP-A), and lifespan (HCP-LS) cohorts. (B) Correlation between structural and functional brain age gap (BAG) estimates. Scatterplots show the relationship between BAG (predicted age − chronological age) estimated by the median-performing structural dMRI QA End CPM model (x-axis) and the median-performing rsfMRI CPM model (y-axis) model for the developmental (HCP-D), young adult (HCP-YA), older adult (HCP-A), and lifespan (HCP-LS) cohorts. Each point represents an individual participant. Solid gray lines indicate the linear regression fit between structural and functional BAG estimates. Insets display the Pearson correlation coefficient (r) and false discovery rate (FDR)-corrected p-value for each cohort. Points are colored according to cohort (HCP-D, blue; HCP-YA, green; HCP-A, purple; HCP-LS, orange).

Across both modalities, significant negative correlations were observed between BAG and chronological age in all cohorts (all *r*’s < −0.6, all FDR-corrected *p*’s <0.001; **Figure 3A**). This pattern indicates that younger participants tended to have positive BAG values, whereas older participants tended to have negative BAG values, reflecting overestimation at younger ages and underestimation at older ages. This phenomenon is commonly observed in brain-age prediction studies and is consistent with age-related prediction bias (Le et al., 2018; Liang et al., 2019; Smith et al., 2019; Treder et al., 2021). Most cohorts and modalities showed non-significant association between predicted age and BAG estimates (**Figure 3B**). A positive correlation was observed in the HCP-YA cohort (dMRI: *r* = 0.236, FDR-corrected *p* < 0.001; rsfMRI: *r* = 0.164, FDR-corrected *p* < 0.001).

**Figure 3.**
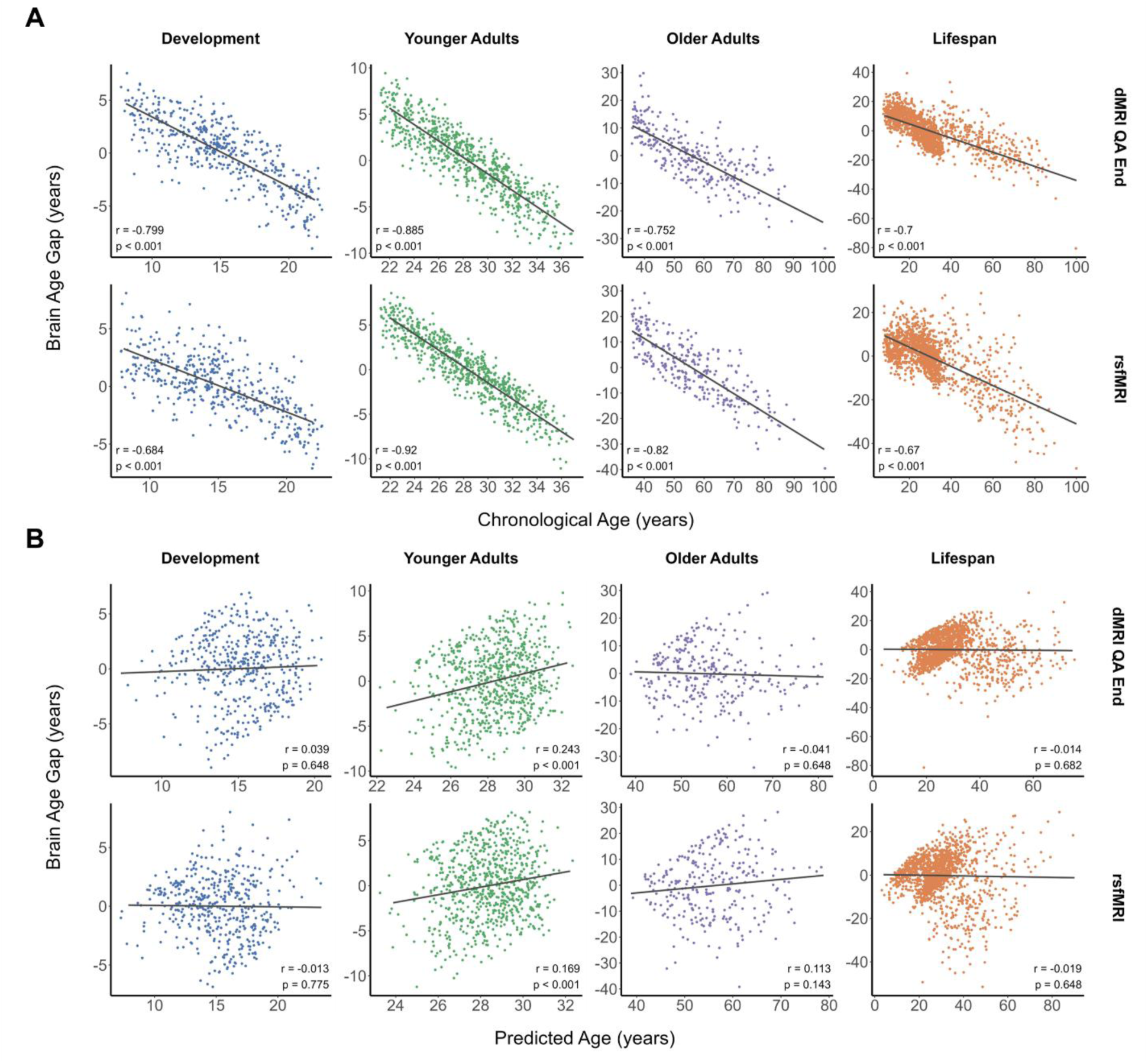
Relationships between brain age gap with chronological and predicted ages. (A) Relationship between brain age gap (BAG) and chronological age. Scatterplots show brain age gap (BAG; predicted age − chronological age) as a function of chronological age for the median-performing CPM models in the developmental (HCP-D), young adult (HCP-YA), older adult (HCP-A), and lifespan (HCP-LS) cohorts. The top row shows structural connectivity models derived from dMRI QA End, and the bottom row shows rsfMRI models. (B) Relationship between brain age gap (BAG) and predicted age. Scatterplots show brain age gap (BAG; predicted age − chronological age; y-axis) as a function of CPM predicted age (x-axis) for the median-performing structural and functional CPM models in the developmental (HCP-D), young adult (HCP-YA), older adult (HCP-A), and lifespan (HCP-LS) cohorts. The top row shows dMRI QA End models, and the bottom row shows rsfMRI models. Each point represents an individual participant. Solid gray lines indicate the linear regression fit between predicted age and BAG. Insets display the Pearson correlation coefficient (r) and false discovery rate (FDR)-corrected p-value for each cohort and modality. Points are colored according to cohort (HCP-D, blue; HCP-YA, green; HCP-A, purple; HCP-LS, orange).

### Connectome features involved in age prediction: Whole brain predictive networks

**Figure 4** shows the brain regions and connections most predictive of chronological age for each age cohort and imaging modality. Predictive information about age appears to vary across age cohorts and imaging modality, with stronger involvement of subcortical and cerebellar networks for dMRI models and somatomotor, salience, subcortical, frontoparietal, and cerebellar involvement for rsfMRI models.

**Figure 4.**
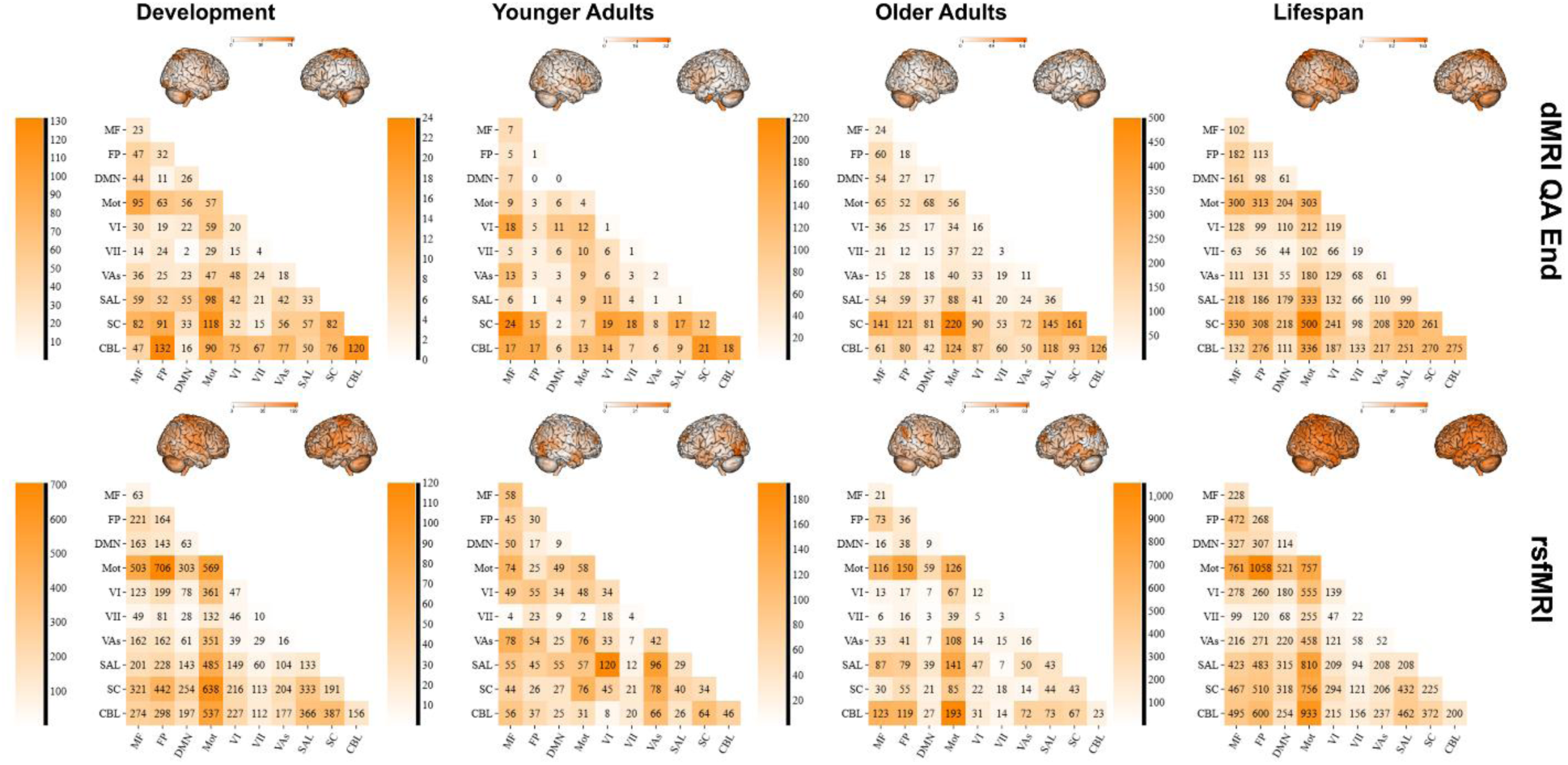
Consensus predictive connectivity patterns contributing to CPM-based age prediction. Visualization of the median-performing CPM consensus networks illustrating the spatial distribution of connectivity features that contributed to age prediction across the developmental (HCP-D), young adult (HCP-YA), older adult (HCP-A), and lifespan (HCP-LS) cohorts. The top row displays networks from dMRI QA End, and the bottom row displays networks from rsfMRI. Brain renderings above each matrix illustrate the anatomical distribution of age-predictive edges mapped onto the Shen 268-node atlas.

Further, the number of edges in the models also differed across imaging modalities and cohorts (**Table 2**). Functional models generally identified substantially more predictive connections than diffusion models in HCP-D, HCP-YA, and HCP-LS. In contrast, HCP-A exhibited comparable edge densities between modalities.

**Table 2.** Total predictive edge densities. Total edge density for each imaging modality in age prediction from the median performing CPM predictive networks.

| Age Cohort | dMRI QA End Total Edges | rsfMRI Total Edges |
| --- | --- | --- |
| HCP-Development | 2,646 | 11,818 |
| HCP-Young Adults | 740 | 2,299 |
| HCP-Aging | 3,136 | 2,645 |
| HCP-Lifespan | 9,585 | 18,235 |

HCP, Human Connectome Project; dMRI, diffusion MRI; QA, quantitative anisotropy; rsfMRI, resting-state functional MRI

Corresponding absolute sum matrices show the magnitude of predictive edge contributions between atlas-defined regions, with higher values indicating regions connected by a greater number of predictive edges regardless of directionality.

### Brain age prediction from individual canonical networks

Individual network CPM analyses demonstrated that age-predictive information was broadly distributed across canonical brain networks, although performance varied across cohorts and modalities. Within-network prediction from both structural and functional connectomes was strongest in HCP-D, HCP-A, and HCP-LS cohorts and weakest in HCP-YA (**Figure 5A** and **Table S5**). Across cohorts, the cerebellar, frontoparietal, salience, and subcortical networks consistently demonstrated the strongest age prediction for both modalities. However, rsfMRI generally produced stronger age prediction than dMRI, with significant age prediction observed for most functional networks across cohorts, whereas structural prediction was more variable and weaker in HCP-YA.

**Figure 5.**
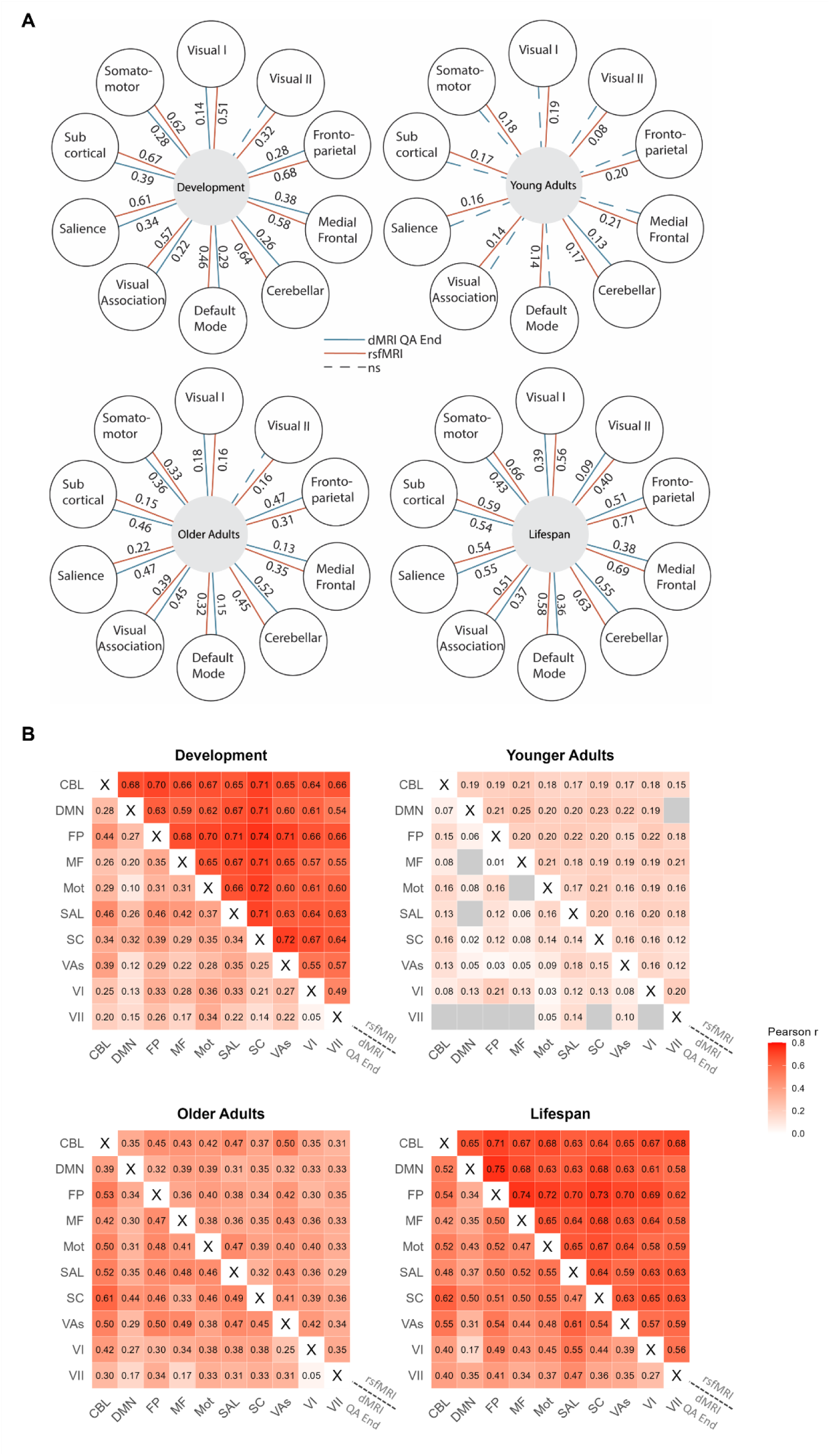
Individual network-level CPM age prediction performance across the lifespan. (A) Individual within-network CPM age prediction performance. Flower plots display the median-performing individual within-network CPM age prediction models for each cohort: developmental (top left), young adult (top right), older adult (bottom left), and lifespan (bottom right). Petals represent the 10 Shen 268-node atlas networks, the center denotes the age cohort, and stems indicate the Pearson correlation coefficient (r) between predicted and chronological age for each network-specific model. dMRI QA End models are shown with blue lines, and rsfMRI models are shown with red lines. Solid lines indicate network-level models that achieved significance following permutation testing against the null distribution, whereas dashed lines indicate non-significant models. The plots highlight modality- and network-specific variation in CPM age prediction performance across age groups. (B) Individual between-network CPM age prediction performance. Heatmaps display Pearson correlation coefficients (r) for significant between-network CPM age prediction models after FDR correction across developmental, young adult, older adult, and lifespan cohorts. The upper triangle represents rsfMRI models, and the lower triangle represents dMRI QA End models. The diagonal is excluded. Non-significant connections are shown in gray, and significant connections are color-coded according to predictive performance (Pearson r). Values within each cell indicate the corresponding Pearson correlation coefficient. Network abbreviations: CBL = cerebellar, DMN = default mode network, FP = frontoparietal, MF = medial frontal, Mot = somatomotor, SAL = salience, SC = subcortical, VAs = visual association, VI = visual I, VII = visual II.

Between-network analyses similarly demonstrated widespread age-related predictive information for both structural and functional connectomes (**Figure 5B** and **Table S5**). For dMRI, between-network prediction was broadly distributed in HCP-D, HCP-A, and HCP-LS but weaker in HCP-YA. rsfMRI showed widespread significant between-network prediction across nearly all network pairs in every cohort despite smaller effect sizes in HCP-YA. Across both modalities, the cerebellar, frontoparietal, salience, somatomotor, and subcortical networks consistently showed strong age prediction, while visual II and some default mode network interactions contributed less consistently. rsfMRI also exhibited stronger predictive performance than dMRI.

### Structural-functional convergence and cross-modality prediction

The structural-functional convergence analyses revealed that at the edge level, significant correlations between modality edge vectors were observed in HCP-D for positive (*r* = 0.012, FDR-corrected *p* = 0.040), negative (*r* = 0.023, FDR-corrected *p* = 0.001), and absolute (*r* = 0.015, FDR-corrected *p* = 0.016) networks, whereas HCP-A demonstrated significant negative correlations for positive (*r* = −0.014, FDR-corrected *p* = 0.016), negative (*r* = −0.020, FDR-corrected *p* = 0.001), and absolute (*r* = −0.020, FDR-corrected *p* = 0.001) networks. However, these had low effect sizes. No edge-level convergence was observed in HCP-YA or HCP-LS. The hypergeometric cumulative density test showed similar patterns.

No node-degree associations survived FDR correction across any cohort. At the network level, significant network density convergence was observed in HCP-D for negative (*r* = 0.411, FDR-corrected *p* = 0.026) and absolute (*r* = 0.515, FDR-corrected *p* = 0.002) networks, whereas HCP-A demonstrated significant divergence for the positive network (*r* = −0.661, FDR-corrected *p* = 0.001). See complete results in **Figure S1A and Table S3.**

Cross-modality transfer between dMRI QA End and rsfMRI was limited and strongly cohort-dependent (**Figure S1** and **Table S4**). In HCP-D, dMRI QA End-trained models showed modest but significant prediction of rsfMRI connectivity (*r* = 0.313, FDR-corrected *p* = 6.71×10⁻12), with weaker reverse transfer from rsfMRI to dMRI QA End (*r* = 0.152, FDR-corrected *p* = 1.14×10⁻3). In HCP-YA, HCP-A, and HCP-LS, cross-modal transfer was generally weak or non-significant in both directions, with rsfMRI models showing particularly limited generalization to diffusion connectomes.

### Sex-stratified age prediction and cross-sex model generalizability

The male-only and female-only models showed significant Pearson *r* correlations (**Figure 6A** and **Table S6**). Across all cohorts, no significant sex differences in prediction performance were observed after FDR correction (**Table S7**). Although the HCP-LS cohort showed a greater prediction accuracy for males than females for the structural connectomes prior to multiple-comparison correction, this difference did not survive FDR correction. Cross-sex prediction was robust and statistically significant across cohorts (**Figure 6B and Table S8**), with the strongest effects observed in the HCP-LS cohort.

**Figure 6.**
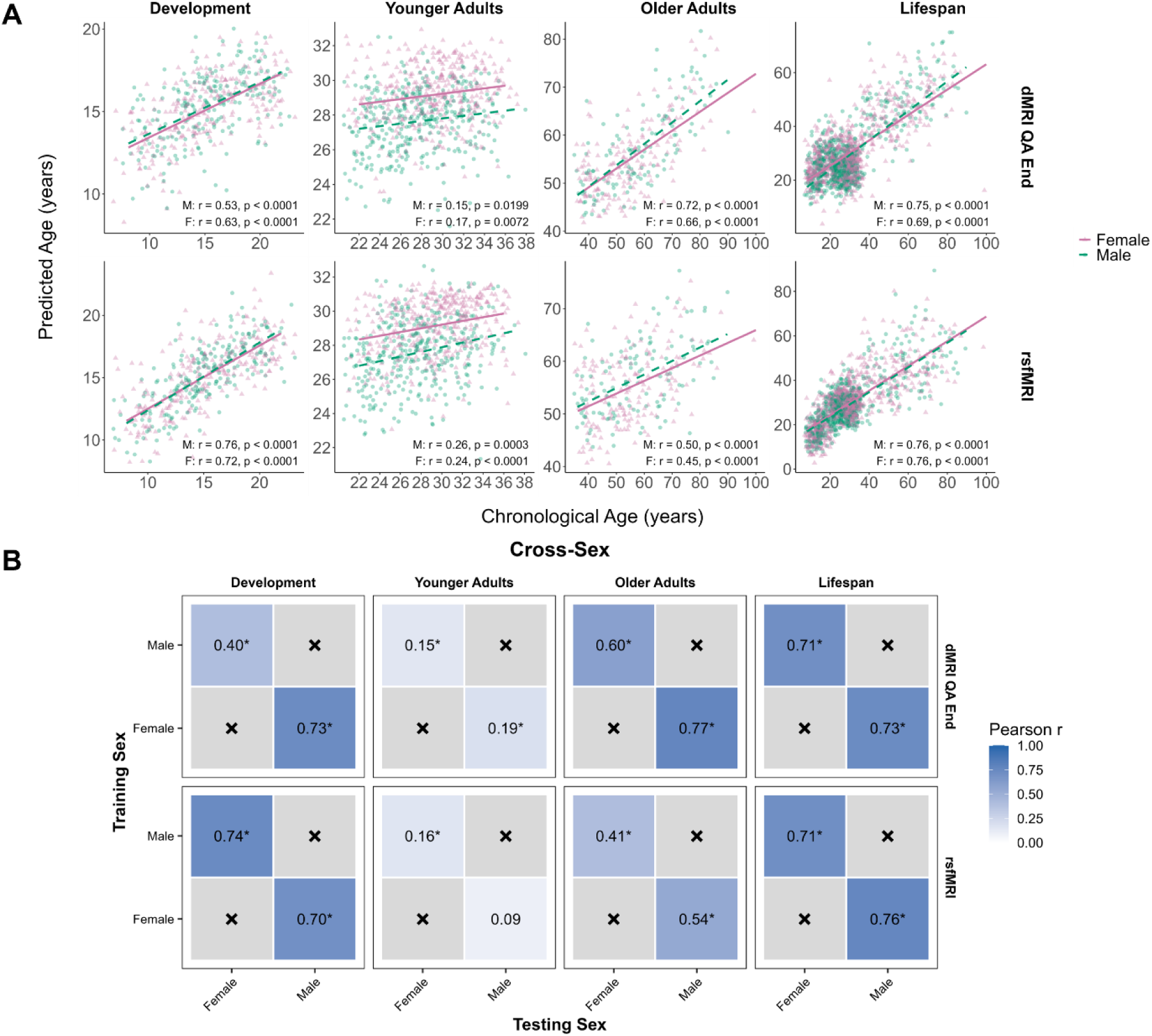
Sex-stratified CPM and cross-sex generalizability. (A) Sex-stratified CPM age prediction performance. Predicted age plotted against chronological age for male-only and female-only CPM models across age cohorts and imaging modalities. Male-only and female-only models were trained separately within each cohort, and sex-specific regression lines were fitted to visualize prediction accuracy, age-related prediction bias, and the relationship between chronological and predicted age. Male models are shown in green and female models are shown in pink. (B) Heatmaps summarize the generalizability of final CPM age prediction models across sex evaluated through external validation/cross-sex prediction.

Heatmap values represent the Pearson correlation coefficient (r) between predicted and chronological age when models were applied to independent datasets. Darker shading indicates stronger age prediction performance (higher Pearson r). Asterisks indicate predictions that remained significant after false discovery rate (FDR) correction (p < 0.05). Gray cells marked with × indicate comparisons that were not performed (e.g., identical training and testing sex).

### Multimodal CPM

We found that the multimodal CPM significantly predicted age for all aging cohorts, following similar cohort and modality trends as observed for unimodal CPM models. Representative consensus networks were reconstructed separately for the dMRI QA End and rsfMRI components of the median-performing multimodal CPM model for each age cohort (**Table S1**). Also, the modality-specific connectivity patterns from the multimodal model were widely distributed across the brain (**Figure S2**). Additionally, consistent with the unimodal models, the developmental, young adult, and lifespan multimodal models contained a greater number of rsfMRI than dMRI QA End predictive edges, whereas the aging cohort exhibited a more balanced distribution of structural and functional predictive edges (**Table S9**).

For the combined dMRI QA End and rsfMRI model, multimodal CPM significantly outperformed the unimodal dMRI QA End model across HCP-D, HCP-YA, and HCP-LS cohorts, with higher Pearson correlations observed in 96–100% of cross-validation iterations. However, multimodal CPM did not improve prediction relative to dMRI QA End in HCP-A, where the dMRI QA End model performed better across all iterations. When compared with rsfMRI-only models, multimodal CPM showed significant improvements in HCP-A, HCP-YA, and HCP-LS, with higher Pearson correlations observed in 99–100% of iterations. In contrast, the multimodal model did not improve prediction over rsfMRI in HCP-D, where rsfMRI demonstrated higher performance across most iterations. Similar patterns were observed for MSE and MAE. Results are shown in **Figure 7** and **Table S10**.

**Figure 7.**
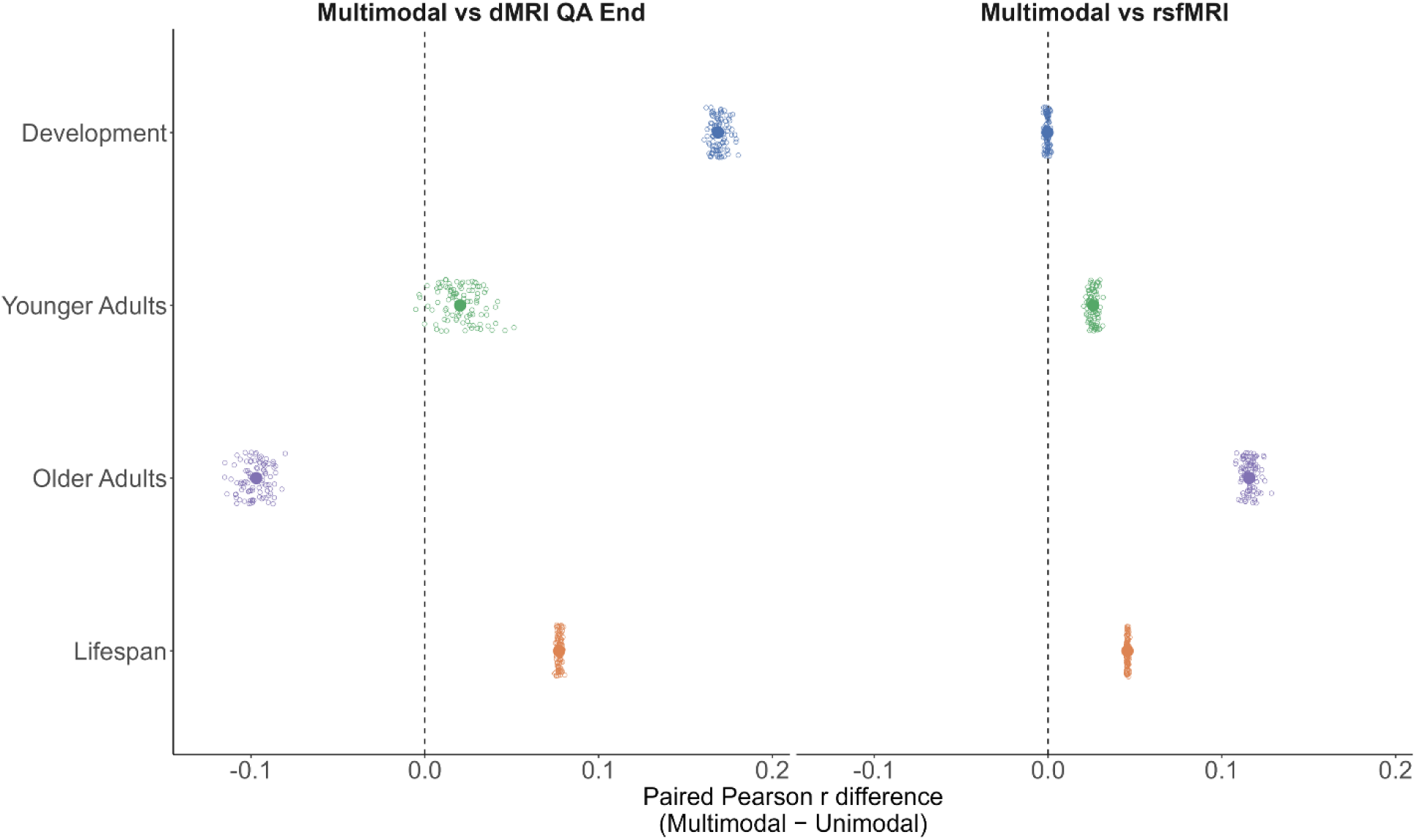
Paired comparison of multimodal and unimodal CPM age prediction performance using dMRI QA End and rsfMRI connectomes. Distribution of paired iteration-wise differences in Pearson correlation coefficients between multimodal CPM combining dMRI QA End and rsfMRI connectomes and corresponding unimodal CPM models across HCP cohorts. Each point represents one of 100 paired cross-validation iterations, and larger points indicate the median difference across iterations. Positive values indicate higher prediction performance for the multimodal model, whereas negative values indicate higher performance for the corresponding unimodal model. Comparisons are shown for multimodal CPM versus dMRI QA End alone (left) and multimodal CPM versus rsfMRI alone (right) across Development (HCP-D), Young Adult (HCP-YA), Aging (HCP-A), and Lifespan (HCP-LS) cohorts.

### Multimodal-unimodal CPM similarity

Multimodal-unimodal similarity was assessed at the edge, node, and network levels for positive, negative, and absolute predictive networks (**Table S11**). Across all cohorts, multimodal-derived predictive information closely resembled their unimodal counterparts. Edge-weight distributions, node-degree profiles, and network edge-density patterns were all highly correlated, and predictive edge overlap was significantly greater than expected by chance.

## Discussion

This study examined lifespan brain age prediction from structural and functional connectomes using CPM across HCP Development, Young Adult, Aging, and Lifespan cohorts. Both structural and functional connectomes significantly predicted chronological age across all cohorts, but prediction performance varied with age range and modality. Prediction was weakest in young adulthood and stronger in development, older adulthood, and the combined lifespan sample. Edges that predicted age were distributed across multiple canonical networks. Structural and functional predicted ages were correlated in most cohorts, yet multimodal CPM improved prediction relative to at least one unimodal model in every cohort while retaining modality-specific predictive architecture.

### Lifespan variability in connectome-based age prediction

The marked variation in prediction performance across cohorts suggests that connectome-based age prediction depends strongly on lifespan stage. Prediction was generally weakest in HCP-YA and stronger in HCP-D, HCP-A, and HCP-LS based on the relationship between chronological age and predicted age. This pattern is consistent with established nonlinear trajectories of white matter and functional network organization across development and aging, with more pronounced changes during development and later life than during early adulthood (Edde et al., 2021; Fjell and Walhovd, 2010; Jockwitz and Caspers, 2021; Yap et al., 2013). Also, while HCP-YA had a weaker Pearson *r* it had a relatively strong MAE, which could be driven by a relatively narrow age range. The magnitude of prediction was broadly comparable to previous connectome-based age-prediction studies, including structural connectome in HCP-YA (Kopetzky et al., 2024) and functional connectome in healthy adult lifespan samples (Kim et al., 2022).

The stronger performance of the HCP-LS models likely reflects, in part, the broader age range and greater chronological age variance available for prediction (Butler et al., 2021; de Lange et al., 2022). Thus, the higher prediction observed in HCP-LS should be interpreted in the context of its broader age distribution. More broadly, these results extend prior CPM studies, which have generally examined narrower developmental or adult age ranges (Kim et al., 2022; Kopetzky et al., 2024), by demonstrating how structural and functional models vary in performance across multiple stages of the lifespan within a common modeling framework.

### Structural and functional connectomes capture complementary aspects of brain maturation and aging

The stronger structural prediction in HCP-A and the functional prediction in HCP-D further suggest that the relative contributions of structural and functional connectivity to age prediction vary across the lifespan. While both modalities predicted chronological age, their relative predictive performance differed across the lifespan. Functional connectomes showed stronger prediction during development, and structural connectomes showed stronger prediction in older adulthood. This pattern is consistent with age-related differences in the trajectories of functional network organization and structural connectivity across the lifespan (Damoiseaux, 2017; Edde et al., 2021; Yap et al., 2013). The stronger structural prediction in HCP-A is also directionally consistent with another study which found substantially stronger age prediction from diffusion-based white-matter measures than resting-state functional connectivity in a healthy adult sample (de Lange et al., 2020). More importantly, structural and functional predicted ages were correlated across cohorts despite limited correspondence between their predictive features, suggesting that the two modalities appear to capture shared age-related information through distinct feature patterns. In all, structural and functional connectomes appear to provide complementary routes to age prediction, with variations of correspondence across the lifespan. This is consistent with evidence that structural-functional coupling changes with age rather than remaining fixed across the lifespan (Damoiseaux, 2017). It also extends this literature by showing that the connectivity features selected for age prediction can diverge even when the resulting age estimates converge.

### Brain age is encoded in distributed network organization

Predictive edges were broadly distributed across the connectome, with contributions from multiple canonical networks and both within- and between-network connections. This finding is consistent with the increasingly network-based view of brain maturation and aging, in which age-related differences reflect coordinated changes across distributed systems rather than isolated regional effects (Damoiseaux, 2017; Jockwitz and Caspers, 2021). Functional models generally contained more edges than structural models, particularly in HCP-D, HCP-YA, and HCP-LS, whereas the number of edges between functional and structural were more comparable in HCP-A.

The specific networks implicated also differed by modality. Functional models consistently involved somatomotor, salience, frontoparietal, subcortical, and cerebellar systems, while structural models showed prominent subcortical and cerebellar involvement. The functional finding is consistent with previous CPM work identifying subcortical-cerebellar connectivity as an important contributor to adult age prediction (Kim et al., 2022). The prominent contribution of subcortical systems in our structural models is also consistent with prior structural brain-age work showing that the subcortex was a more reliable predictor of age than cortical features in healthy adults (Xifra-Porxas et al., 2021). Network-specific analyses further showed that age could be predicted from both within and between network connectivity, with frontoparietal, salience, cerebellar, somatomotor, and subcortical networks contributing across analyses. Thus, age-related predictive information was distributed across multiple levels of network organization rather than confined to a particular canonical network. The cohort- and modality-specific differences further suggest that the systems supporting age prediction vary across stages of the lifespan rather than constituting a fixed brain-aging network.

### Multimodal CPM improves age prediction while preserving modality-specific signatures

Multimodal CPM improved prediction relative to at least one unimodal model in every cohort and outperformed both unimodal models in HCP-YA and HCP-LS. The magnitude and direction of this benefit varied across the lifespan. Adding rsfMRI improved upon structural models in HCP-D, HCP-YA, and HCP-LS, whereas adding structural connectivity improved upon rsfMRI in HCP-YA, HCP-A, and HCP-LS. In HCP-A, dMRI QA End alone outperformed the multimodal model, while rsfMRI alone performed comparably or better than the multimodal model in HCP-D. These findings indicate that the value of multimodal integration is not uniform across the lifespan but instead depends on the relative amount of nonredundant age-related information contributed by each modality. Multimodal integration may provide the greatest benefit when structural and functional connectivity capture complementary sources of age-related variation, whereas limited improvement may occur when a single modality already captures much of the available predictive signal. This pattern is consistent with previous multimodal brain-age studies showing improved prediction from combining structural and functional features (Liem et al., 2017; Niu et al., 2020).

Furthermore, multimodal integration did not substantially alter the predictive patterns identified by the unimodal models. Structural and functional components of the multimodal models closely resembled their respective unimodal networks at the edge, node, and network levels, with rsfMRI contributing more predictive edges in most cohorts and a more balanced structural-functional contribution in HCP-A. Thus, multimodal improvement appears to arise from integration of complementary modality-specific representations rather than from identifying a new common predictive network.

### Sex-related variability and cross-sex generalizability

Despite evidence for sex-related differences in brain structure and functional connectivity across development and aging (Kaczkurkin et al., 2019; Ritchie et al., 2018; Satterthwaite et al., 2015; Scheinost et al., 2015), prediction performance did not differ significantly between males and females for either modality after multiple-comparison correction. Further, models trained in one sex generalized robustly to the other across nearly all cohorts. This is consistent with a study that reported similar brain-age prediction errors in males and females and highly correlated predictions from sex-specific and mixed-sex models (Bashyam et al., 2020). Another study also found minimal sex differences in global brain-age estimates despite regional sex differences in brain-age patterns (Sanford et al., 2022). Comparable whole-brain prediction performance does not necessarily imply identical neurobiological aging trajectories.

Instead, our findings suggest that the connectivity features supporting chronological age prediction are sufficiently shared between sexes to support strong cross-sex generalization, while sex-related differences in brain organization may be expressed at more localized or modality-specific levels.

### Limitations and future directions

Several limitations warrant consideration. First, the HCP cohorts comprise relatively healthy participants and may not capture the demographic and clinical heterogeneity of the broader population. External validation across independent samples, scanners, and acquisition protocols will therefore be necessary to test model generalizability. Second, cohorts differed in age range, sample size, and age distribution, complicating direct comparisons of prediction performance. Third, the analyses were cross-sectional.

Longitudinal validation will be important to determine whether the predictive signatures identified here track individual changes over time. Finally, structural models were not adjusted for global anatomical measures such as intracranial volume or regional morphology, which may contribute to some of the observed age-predictive signal.

Future work should extend these findings by addressing some of the limitations and testing whether deviations from these normative patterns are associated with cognitive and clinical outcomes.

Longitudinal and nonlinear modeling approaches may further clarify how structural-functional contributions to brain age change within individuals across the lifespan.

### Conclusion

Using large-scale HCP datasets spanning childhood through older adulthood, this study demonstrates that structural and functional connectomes provide robust and complementary information for predicting chronological age, with prediction strength varying across lifespan stages and modalities. Predictive network patterns were widespread across the brain, while structural and functional models captured shared age-related information through largely distinct predictive features. Multimodal CPM generally improved prediction by integrating these complementary representations while preserving modality-specific predictive architectures. Overall, these findings demonstrate the utility of CPM for characterizing distributed connectivity patterns associated with chronological age across the lifespan and provide a foundation for future studies investigating individual variability in brain maturation and aging.

## Acknowledgements

M.M. was supported by the National Science Foundation Graduate Research Fellowship under grant DGE2139841.

