## Supplementary_Materials for "Brain age prediction from structural and functional connectivity across the lifespan using connectome-based predictive modeling"

**Supplement Methods**

#### Data Processing

Diffusion MRI data were acquired on 3T Siemens scanners using HCP diffusion protocols. HCP-YA data were acquired on a Siemens Skyra using a multishell acquisition with a maximum b-value of 3000 s/mm², 1.5-mm isotropic resolution, and 1.5-mm slice thickness. HCP-D and HCP-A data were acquired on Siemens Prisma scanners using the corresponding HCP protocols; HCP-D included 185 diffusion directions across b = 1500 and 3000 s/mm² shells and 28 b = 0 s/mm² images. HCP-D and HCP-A underwent the same diffusion MRI preprocessing and tractography pipeline. Susceptibility-related distortions were corrected using reversed phase-encoding b = 0 images with TOPUP from Tiny FSL (http://github.com/frankyeh/TinyFSL), and diffusion data were aligned to the AC-PC line with b-table orientation verified against a population-averaged template (Yeh et al., 2018).

Diffusion data were reconstructed using generalized q-sampling imaging (Yeh et al., 2010) with a diffusion sampling length ratio of 1.25. Whole-brain deterministic tractography was performed in DSI Studio using a modified fiber assignment by continuous tracking (FACT) algorithm, with quantitative anisotropy (QA) used as the tract termination criterion. The pairwise connectivity strength was calculated as the average QA value of each fiber connecting the two end regions. For HCP-YA, diffusion spin density functions were reconstructed in native space and nonlinearly registered to an MNI-space template using SPM. For all datasets, tracking continued until 10 million streamlines were generated per participant, using an angular cutoff of 60°, 1.0-mm step size, and minimum and maximum tract lengths of 30 and 300 mm, respectively.

Resting-state functional MRI data were acquired on 3T MRI scanners using multiband echo-planar imaging and minimally preprocessed data were used (Glasser et al., 2013). Additional preprocessing was performed using BioImage Suite (Joshi et al., 2011; Papademetris et al., 2006) and custom MATLAB scripts. Motion correction was performed using SPM12. Nuisance regression included linear and quadratic drift terms, mean cerebrospinal fluid, white matter, and global gray-matter signals, and a 24-parameter motion model consisting of six rigid-body motion parameters, their temporal derivatives, and squared terms. Temporal smoothing was performed using a Gaussian filter with an approximate cutoff frequency of 0.12 Hz. Mean framewise displacement (FD) was calculated for each participant, and participants with mean FD >0.1 mm were excluded. Mean FD was additionally included as a covariate during feature selection within each training fold, with covariate adjustment performed independently within each training fold to prevent information leakage from held-out test data.

#### Individual network age prediction

To further examine connectome contributions to age prediction, we conducted network-specific age predictions. Participant connectomes were parcellated into 10 predefined network masks derived from a 268-node atlas (Shen et al., 2013). For each within- and between-network combination, the same CPM procedure was applied independently for each age cohort and imaging modality. For each network mask, age prediction was generated using only the connections within that network using a CPM framework with predefined train-test folds. Network-level performance was assessed based on the median performing run for each network model using the same Pearson correlation metrics. Statistical significance was assessed using a nonparametric permutation test with 1,000 iterations, in which age labels were randomly shuffled within cohort while preserving connectome structure, network masks, motion covariates, and predefined train-test splits.

#### Sex-stratified age prediction

To determine whether connectome-based age prediction performance differed between males and females, CPM analyses were performed separately within each sex group for each age cohort and imaging modality. Each sex-specific model uses an independently generated 10-fold CV partition, with approximately 90% of that sex's participants used for training and 10% for testing, while the same fold assignments are reused across imaging modalities within that sex. Male-only and female-only models were trained using identical procedures as the full-sample analyses, including connectome feature extraction, feature selection, model training, and repeated cross-validation. Prediction performance was evaluated across 100 repeated CPM iterations using Pearson correlation between predicted and chronological age, MAE, and MSE. We identified the median-performing run for each age cohort and imaging modality.

**Supplement Ridge Regression rCPM Methods**

**Ridge Regression Connectome-Based Predictive Modeling (rCPM)**

We implemented ridge regression-based connectome-based predictive modeling (rCPM) to predict chronological age from whole-brain structural (diffusion MRI QA end) and functional (resting-state fMRI) connectivity (Gao et al., 2019). Analyses were performed separately within each cohort (HCP-D, HCP-YA, HCP-A, and HCP-LS) using a 268-node Shen atlas. For each participant, the upper triangular connectivity matrix was vectorized to form an edge-wise feature vector.

Models were trained using k-fold cross-validation with predefined and fixed subject assignments to ensure consistency across iterations and comparability across analyses. Each participant served as an independent test case exactly once per fold. This procedure was repeated across multiple iterations (100 for main analyses and up to 10,000 for permutation testing), while preserving identical fold structures across runs.

Within each training fold, edges were screened for their association with age using Pearson correlation or partial correlation when motion covariates were included. Covariate adjustment was applied only during this feature selection step in the training data. Edges meeting a significance threshold of $p < 0.05$ were retained for subsequent multivariate modeling. This ensured that feature selection was performed exclusively within the training set to avoid information leakage.

Age prediction was performed using a ridge-regularized linear regression model implemented through MATLAB's lasso function with an extremely small mixing parameter ($\alpha = 1e{-6}$), yielding an L2-dominated penalty. In this formulation, the model estimates a set of edge-wise weights that jointly predict age while constraining coefficient magnitude to reduce overfitting in the high-dimensional feature space, where the number of edges far exceeds the number of subjects.

The regularization strength ($\lambda$) was selected independently within each training fold using internal 10-fold cross-validation. The final model corresponds to the $\lambda$ value minimizing cross-validated prediction error (or the 1-standard-error solution when applicable). The fitted model takes the form:

$$\hat{y} = X^{(sel)}\beta + \beta_0$$

where $X^{(sel)}$is the matrix of selected edges within the training fold.

The trained model was applied to held-out test participants using only edges selected in the corresponding training fold. This yielded one out-of-sample age prediction per participant per fold. Predictions were concatenated across folds to generate a single predicted age estimate for each subject.

Model performance was evaluated across folds by comparing predicted and observed age using Pearson's correlation coefficient (r), Spearman's rank correlation ($\rho$), mean squared error (MSE), mean absolute error (MAE), and predictive coefficient of determination ($q^2$). These metrics provide a comprehensive assessment of CPM performance by evaluating correlation strength, ranking accuracy, prediction error magnitude, and overall explained variance in out-of-sample age prediction. Performance was represented across iterations using the median performing run to provide a robust estimate of predictive accuracy.

Statistical significance was assessed using permutation testing. For null models, age labels were randomly permuted prior to model training while preserving connectivity structure and fold assignments. The full modeling pipeline was repeated across 10,000 permutations to generate empirical null distributions of performance metrics. Empirical *p*-values were computed based on the proportion of permuted correlations exceeding observed model performance.

**Sex-Stratified Age Prediction Using Ridge Regression**

Same as in CPM.

**Exploratory analyses of model characteristics and bias**

Same as in CPM.

**Median Performing Representative rCPM Model**

For rCPM, predictive information was represented by continuous regression coefficients rather than binary edge selection. We derived a representative weighted predictive network from the repeated cross-validation models. For each rCPM iteration, ridge regression coefficients from each cross-validation fold were projected back onto their corresponding connectome edges, generating fold-specific weighted adjacency matrices. Matrices were symmetrized and averaged across folds to obtain a consensus coefficient network for each repeated iteration.

Model iterations were ranked by cross-validated predictive performance, defined as the Pearson correlation between predicted and observed phenotype values. The iteration with median predictive performance was selected to represent the overall rCPM model. The corresponding averaged coefficient matrix was retained as the median rCPM network. Signed coefficient matrices were used to preserve information regarding the direction of prediction, with positive and negative coefficients indicating edges associated with increases or decreases in predicted phenotype values, respectively. Absolute coefficient matrices were also calculated to assess the magnitude of predictive contribution independent of sign.

**Structural-functional rCPM coefficient convergence**

To evaluate the extent to which structural and functional connectomes identify overlapping predictive architecture, we performed a complementary structural–functional convergence analysis using the median-performing rCPM coefficient networks from each cohort. This analysis quantified the similarity of the connection weights contributing to age prediction across dMRI QA End and rsfMRI models. For each cohort, the median-performing rCPM coefficient matrix was extracted separately for structural and functional models, resulting in weighted 268 × 268 adjacency matrices corresponding to the Shen268 parcellation.

Structural-functional convergence was evaluated at three spatial scales: edge, node, and canonical network levels. At the edge level, corresponding connections between structural and functional coefficient matrices were compared using Pearson correlation across the upper triangle of the adjacency matrix. Analyses were performed using both signed coefficients, which preserve the direction of prediction-related associations, and absolute coefficients, which quantify convergence in coefficient magnitude independent of direction. Edges with zero coefficients in both models were excluded from edge-level comparisons to avoid inflation from shared non-predictive connections.

At the node level, regional contributions to prediction were quantified by summing the absolute values of all incident edge coefficients for each node. These node-level contribution profiles represent the overall predictive importance of each brain region within the structural and functional models. Structural and functional node contribution profiles were compared using Pearson correlation to determine whether regions contributing strongly to age prediction in one modality also contributed strongly in the other modality.

At the network level, coefficient organization was summarized according to the 10 canonical functional systems defined by the Shen268 atlas. For each modality, mean coefficient contribution was calculated for connections within and between each pair of canonical networks, generating a 10 × 10 network contribution matrix. Structural and functional network contribution matrices were compared by correlating the corresponding upper-triangle network pairs. Network-level analyses were performed separately using signed and absolute coefficient representations to evaluate convergence of both coefficient direction and coefficient magnitude.

Statistical significance for each convergence metric was assessed using permutation testing (10,000 iterations). For edge-level analyses, null distributions were generated by randomly permuting structural coefficient assignments while preserving the functional coefficient vector. For node-level analyses, node identities in the structural contribution profile were randomly permuted. For network-level analyses, canonical network labels were permuted while maintaining the overall network contribution structure. Empirical p-values were calculated as the proportion of permutation-derived correlations equal to or exceeding the observed correlation. Resulting p-values were corrected for multiple comparisons across analyses using the Benjamini–Hochberg FDR with a significance threshold of $p < 0.05$.

**Individual network age prediction**

Individual network prediction using rCPM followed the same procedure described before for whole-brain models, except that analyses were performed separately on connectomes masked to individual within- and between-network connections. To better accommodate the reduced feature space within individual network masks, the regularization parameter (λ) was selected using the minimum cross-validation error criterion (IndexMinMSE) rather than the one-standard-error rule (Index1SE). This approach favored less regularized models to retain predictive features within the smaller network-specific connectomes. Age prediction performance was evaluated using the same performance metrics. Statistical significance was assessed using a nonparametric permutation test with 1,000 iterations.

**Final rCPM model construction**

Using the selected median-performing run, a final rCPM model was retrained using all available training subjects. Feature selection was recomputed on the full dataset using correlation (or partial correlation when covariates were included) between each edge and the target phenotype, thresholded by the original significance criterion derived from the training pipeline. The resulting edge subset was used to construct a subject × feature design matrix from vectorized lower-triangle connectomes. Ridge regression (implemented via elastic net with $\alpha = 1e{-6}$, effectively ridge-dominant regularization) was then performed with 10-fold cross-validation to select the optimal regularization parameter (λ), using the minimum cross-validated error criterion (1-SE rule where applicable). The final model was refit on the full training dataset using the selected λ to obtain stable regression coefficients and intercept. These coefficients were reinserted into 268 × 268 space to generate a symmetric weighted coefficient matrix representing the final rCPM model. This model was only used to reconstruct the ridge regression equation parameters required for external validation cross prediction.

**Model generalizability using cross prediction**

Generalizability of the final rCPM models was evaluated using three external validation frameworks without retraining. First, cross-modality within-cohort prediction was performed by applying each cohort-specific model trained on one imaging modality to connectomes derived from the other modality within the same cohort. For each subject, connectomes were vectorized using the identical lower-triangle ordering defined during model training, and only edges retained in the final model were included in prediction. Age estimates were computed as a linear combination of selected edges and learned coefficients (Predicted Age = $X\beta$ + intercept), and BAG was defined as the difference between predicted and chronological age.

Second, cross-cohort within-modality generalization was assessed by training rCPM models within each cohort and applying them to independent cohorts while holding imaging modality constant. This evaluated whether learned connectome–phenotype mappings generalize across life stages and population samples.

Third, cross-sex generalization was performed by training separate rCPM models in male and female subsamples within each cohort and modality, followed by out-of-sex testing. Male-trained models were applied to female connectomes and vice versa. This allowed assessment of sex-specific versus sex-general predictive structure in connectome-derived age signals.

Across all external validation settings, predictive performance was quantified using Pearson correlation, Spearman correlation, MSE, MAE, and Q-score, and statistical significance was evaluated using FDR correction.

**Multimodal rCPM**

To examine multimodal age prediction using a regularized modeling framework, multimodal ridge regression rCPM was applied within each age cohort. For every participant, dMRI QA End and rsfMRI connectivity matrices were vectorized by extracting the lower triangular elements and concatenating into a single multimodal feature vector, with all structural connectivity edges preceding the functional connectivity edges. Within each training fold, edges significantly associated with chronological age ($p < 0.05$) were identified using univariate correlation and retained as predictors. Ridge regression was then fit to the selected multimodal features, with the regularization parameter ($\lambda$) determined by 10-fold cross-validation within the training data using the one-standard-error rule. Because framewise displacement was available only for the rsfMRI data, head motion was not included as a covariate in the multimodal analyses to avoid modality-specific adjustment. The resulting model generated age predictions by jointly leveraging structural and functional connectivity information. Performance was evaluated using the same framework as the unimodal rCPM analyses, including Pearson correlation, mean absolute error (MAE), mean squared error (MSE), and permutation testing for statistical significance.

**Statistical comparison between multimodal and unimodal rCPM performance**

Same as in CPM.

**Multimodal-unimodal rCPM similarity**

To determine whether incorporating functional connectivity altered the structural predictive architecture identified by dMRI QA End, coefficient similarity was assessed between unimodal dMRI QA End rCPM models and the dMRI QA End component of multimodal rCPM models. Analyses were performed separately within each age cohort using the median-performing rCPM model from the repeated cross-validation procedure. Ridge regression coefficient matrices were reconstructed as 268 × 268 weighted connectivity networks based on the Shen atlas for both unimodal and multimodal models. Similarity between unimodal and multimodal dMRI QA End coefficient networks were evaluated at three spatial scales: edge, node, and network levels as described earlier in the structural-functional convergence section.

### Supplement CPM Results

**Structural and functional connectomes predict chronological age and cross-cohort prediction**

| Modality | Cohort | Pearson r | Null CPM FDR-corrected p | Spearman ρ | MAE | MSE | $q^2$ |
|---|---|---|---|---|---|---|---|
| **dMRI QA End** | HCP-D | 0.568982 | 1.00E-04 | 0.576715 | 2.48851 | 9.613579 | 0.322646 |
| | HCP-YA | 0.235917 | 0.00015 | 0.219248 | 3.019141 | 13.62755 | -0.00348 |
| | HCP-A | 0.689025 | 1.00E-04 | 0.650204 | 7.580617 | 92.65064 | 0.473762 |
| | HCP-LS | 0.723272 | 1.00E-04 | 0.486502 | 8.603228 | 118.8374 | 0.523018 |
| **rsfMRI** | HCP-D | 0.73871 | 1.00E-04 | 0.754528 | 2.023578 | 6.449124 | 0.545607 |
| | HCP-YA | 0.230011 | 1.00E-04 | 0.243854 | 2.98046 | 13.24158 | 0.024936 |
| | HCP-A | 0.476254 | 1.00E-04 | 0.448079 | 9.470178 | 137.9459 | 0.216493 |
| | HCP-LS | 0.75476 | 1.00E-04 | 0.719196 | 7.552796 | 107.2911 | 0.569362 |
| **Multimodal dMRI QA End + rsfMRI** | HCP-D | 0.738062 | 1.00E-04 | 0.748614 | 2.036536 | 6.469089 | 0.5442 |
| | HCP-YA | 0.255706 | 1.00E-04 | 0.264641 | 2.968241 | 13.20061 | 0.027953 |
| | HCP-A | 0.592222 | 1.00E-04 | 0.560334 | 8.574933 | 114.4098 | 0.350174 |
| | HCP-LS | 0.800465 | 1.00E-04 | 0.733202 | 7.071322 | 89.68665 | 0.640022 |

*Table S1. CPM performance metrics from the median-performing diffusion MRI (dMRI) QA End, resting-state fMRI (rsfMRI), and multimodal connectome-based predictive models for each age cohort (HCP-D, HCP-YA, HCP-A, and HCP-LS). Model performance was evaluated with Pearson's correlation (r), Spearman's rank correlation (ρ), mean absolute error (MAE), mean squared error (MSE), and the predictive coefficient of determination ($q^2$).*

| Modality | TrainCohort | TestCohort | Pearson r | p-value | FDR-corrected p | Spearman rho | MSE | MAE | $q^2$ |
|---|---|---|---|---|---|---|---|---|---|
| **dMRI QA End** | HCP-D | HCP-YA | 0.070519 | 0.05806 | 0.074649 | 0.070501 | 442.0004 | 20.43447 | -31.5474 |
| **dMRI QA End** | HCP-D | HCP-A | 0.217466 | 0.00019 | 0.000326 | 0.202405 | 2031.649 | 43.1701 | -10.5394 |
| **dMRI QA End** | HCP-D | HCP-LS | 0.435278 | 4.68E-69 | 2.81E-68 | 0.35743 | 637.7614 | 20.76784 | -1.55981 |
| **dMRI QA End** | HCP-YA | HCP-D | 0.177811 | 0.000131 | 0.000235 | 0.178254 | 64.93465 | 7.081542 | -3.57517 |
| **dMRI QA End** | HCP-YA | HCP-A | 0.099737 | 0.090007 | 0.111733 | 0.091634 | 1214.389 | 32.2231 | -5.8975 |
| **dMRI QA End** | HCP-YA | HCP-LS | 0.231671 | 2.25E-19 | 7.35E-19 | 0.376907 | 274.0087 | 10.74038 | -0.0998 |
| **dMRI QA End** | HCP-A | HCP-D | 0.141807 | 0.002351 | 0.003526 | 0.14476 | 272.3424 | 14.40445 | -18.1887 |

| | | | | | | | | | |
|---|---|---|---|---|---|---|---|---|---|
| **dMRI QA End** | HCP-A | HCP-YA | 0.01119 | 0.763903 | 0.808838 | -0.00417 | 285.5398 | 15.01218 | -20.0262 |
| **dMRI QA End** | HCP-A | HCP-LS | 0.45546 | 3.14E-76 | 2.26E-75 | 0.394334 | 307.1592 | 15.35757 | -0.23286 |
| **dMRI QA End** | HCP-LS | HCP-D | 0.414705 | 1.83E-20 | 6.60E-20 | 0.418615 | 1717.983 | 39.62688 | -120.046 |
| **dMRI QA End** | HCP-LS | HCP-YA | 0.103456 | 0.005362 | 0.007721 | 0.094099 | 855.6278 | 26.81907 | -62.0054 |
| **dMRI QA End** | HCP-LS | HCP-A | 0.483765 | 2.04E-18 | 6.11E-18 | 0.488896 | 2459.552 | 46.09521 | -12.9698 |
| **rsfMRI** | HCP-D | HCP-YA | -0.05836 | 0.116898 | 0.135752 | -0.07813 | 94.42301 | 8.469715 | -5.95298 |
| **rsfMRI** | HCP-D | HCP-A | -0.03628 | 0.538333 | 0.599693 | -0.04745 | 1396.389 | 34.49195 | -6.93123 |
| **rsfMRI** | HCP-D | HCP-LS | 0.384234 | 5.98E-53 | 3.07E-52 | 0.393441 | 326.2721 | 11.89328 | -0.30957 |
| **rsfMRI** | HCP-YA | HCP-D | -0.08878 | 0.057631 | 0.074649 | -0.05518 | 82.2812 | 8.063562 | -4.79737 |
| **rsfMRI** | HCP-YA | HCP-A | -0.03527 | 0.549719 | 0.599693 | -0.00342 | 1207.333 | 31.94716 | -5.85743 |
| **rsfMRI** | HCP-YA | HCP-LS | 0.126534 | 1.12E-06 | 2.38E-06 | 0.287 | 276.2328 | 10.81331 | -0.10873 |
| **rsfMRI** | HCP-A | HCP-D | -0.15515 | 0.000864 | 0.001413 | -0.14087 | 611.8152 | 22.9183 | -42.1073 |
| **rsfMRI** | HCP-A | HCP-YA | -0.00839 | 0.821806 | 0.82862 | -0.01006 | 455.9573 | 19.40244 | -32.5751 |
| **rsfMRI** | HCP-A | HCP-LS | 0.502741 | 4.76E-95 | 4.28E-94 | 0.378012 | 453.1572 | 18.97173 | -0.81885 |
| **rsfMRI** | HCP-LS | HCP-D | 0.554238 | 3.08E-38 | 1.38E-37 | 0.56663 | 190.3443 | 11.27187 | -12.4113 |

| | | | | | | | | | |
|---|---|---|---|---|---|---|---|---|---|
| **rsfMRI** | HCP-LS | HCP-YA | 0.059263 | 0.111351 | 0.133622 | 0.029577 | 144.7102 | 9.517515 | -9.65595 |
| **rsfMRI** | HCP-LS | HCP-A | 0.397684 | 1.99E-12 | 5.11E-12 | 0.397707 | 610.2322 | 18.9453 | -2.46601 |

*Table S2. Cross-cohort model generalizability using the final CPM models.*

**Structural-functional convergence and cross-modality model generalizability**

| **Edge-level** | | | | | | | |
|---|---|---|---|---|---|---|---|
| **Cohort** | **Type** | **Edge Correlation r** | **Edge Correlation p** | **Edge Correlation p-FDR** | **Hypergeometric p** | **Hypergeometric p-FDR** | **Jaccard** |
| **HCP-D** | Positive | 0.012291645 | 0.02019798 | 0.04039596 | 0.013428639 | 0.053714558 | 0.015416077 |
| **HCP-D** | Negative | 0.023330154 | 1.00E-04 | 0.00119988 | 9.15E-06 | 0.000109755 | 0.056456932 |
| **HCP-D** | Absolute | 0.01498449 | 0.00539946 | 0.015838416 | 0.002560743 | 0.01536446 | 0.069506063 |
| **HCP-YA** | Positive | 0.0050405 | 0.404359564 | 0.539146085 | 0.228563875 | 0.5485533 | 0.004758128 |
| **HCP-YA** | Negative | -0.003441877 | 0.538546145 | 0.587504886 | 0.773982099 | 0.999983099 | 0.00968661 |
| **HCP-YA** | Absolute | 0.002762723 | 0.596640336 | 0.596640336 | 0.321462068 | 0.642924136 | 0.017068273 |
| **HCP-A** | Positive | -0.014016722 | 0.00659934 | 0.015838416 | 0.998160072 | 0.999983099 | 0.010976088 |
| **HCP-A** | Negative | -0.020022813 | 0.00029997 | 0.00119988 | 0.999983099 | 0.999983099 | 0.012970579 |
| **HCP-A** | Absolute | -0.019581654 | 0.00019998 | 0.00119988 | 0.99994356 | 0.999983099 | 0.032137118 |
| **HCP-LS** | Positive | -0.010007568 | 0.057994201 | 0.09941863 | 0.972223505 | 0.999983099 | 0.103727956 |
| **HCP-LS** | Negative | -0.003965367 | 0.453354665 | 0.544025597 | 0.778898083 | 0.999983099 | 0.081168831 |
| **HCP-LS** | Absolute | 0.005151638 | 0.331466853 | 0.49720028 | 0.167900928 | 0.503702783 | 0.215165546 |
| **Node-level** | | | | | | | |

| Cohort | Type | Node Correlation r | Node Correlation p | Node Correlation p-FDR |
|---|---|---|---|---|
| **HCP-D** | Positive | 0.079526614 | 0.196780322 | 0.386721328 |
| **HCP-D** | Negative | -0.06023046 | 0.322267773 | 0.386721328 |
| **HCP-D** | Absolute | 0.07325841 | 0.221977802 | 0.386721328 |
| **HCP-YA** | Positive | -0.08274415 | 0.176982302 | 0.386721328 |
| **HCP-YA** | Negative | 0.101800263 | 0.091690831 | 0.366763324 |
| **HCP-YA** | Absolute | 0.107662195 | 0.075892411 | 0.366763324 |
| **HCP-A** | Positive | -0.105430397 | 0.086791321 | 0.366763324 |
| **HCP-A** | Negative | 0.014140127 | 0.818218178 | 0.818218178 |
| **HCP-A** | Absolute | -0.064913888 | 0.296970303 | 0.386721328 |
| **HCP-LS** | Positive | -0.064069454 | 0.298570143 | 0.386721328 |
| **HCP-LS** | Negative | -0.054254384 | 0.374962504 | 0.409050004 |
| **HCP-LS** | Absolute | -0.073118594 | 0.233076692 | 0.386721328 |
| | | | | **Network-level** |
| **Cohort** | **Type** | **Upper Triangle r** | **Upper Triangle p** | **Upper Triangle p-FDR** |
| **HCP-D** | Positive | 0.166367609 | 0.273972603 | 0.547945205 |
| **HCP-D** | Negative | 0.411342411 | 0.00639936 | 0.02559744 |
| **HCP-D** | Absolute | 0.514743885 | 0.00039996 | 0.00239976 |
| **HCP-YA** | Positive | -0.113181141 | 0.462253775 | 0.792435042 |
| **HCP-YA** | Negative | -0.009762065 | 0.95080492 | 0.95080492 |

| | | | | |
|---|---|---|---|---|
| **HCP-YA** | Absolute | -0.060453107 | 0.699330067 | 0.821154248 |
| **HCP-A** | Positive | -0.660613905 | 1.00E-04 | 0.00119988 |
| **HCP-A** | Negative | -0.177679545 | 0.239776022 | 0.547945205 |
| **HCP-A** | Absolute | -0.333495393 | 0.02319768 | 0.069593041 |
| **HCP-LS** | Positive | -0.067961758 | 0.644935506 | 0.821154248 |
| **HCP-LS** | Negative | -0.048730923 | 0.752724728 | 0.821154248 |
| **HCP-LS** | Absolute | 0.064626428 | 0.671632837 | 0.821154248 |

*Table S3. CPM structural-functional convergence analysis results at the edge, node, and network levels.*

| **Cohort** | **TrainModality** | **TestModality** | **Pearson r** | **p-value** | **FDR-corrected p** | **Spearman rho** | **MSE** | **MAE** | **q** |
|---|---|---|---|---|---|---|---|---|---|
| **HCP-D** | dMRI QA End | rsfMRI | 0.313458 | 6.71E-12 | 1.61E-11 | 0.30371 | 18.30923 | 3.299658 | -0.29003 |
| **HCP-D** | rsfMRI | dMRI QA End | 0.151514 | 0.001144 | 0.001961 | 0.163976 | 14.9013 | 3.164521 | -0.04992 |
| **HCP-YA** | dMRI QA End | rsfMRI | 0.020214 | 0.58738 | 0.640778 | 0.030483 | 28.23559 | 4.354614 | -1.07917 |
| **HCP-YA** | rsfMRI | dMRI QA End | -0.00864 | 0.816556 | 0.852058 | -0.00457 | 18.89073 | 3.579453 | -0.39105 |
| **HCP-A** | dMRI QA End | rsfMRI | -0.23005 | 7.69E-05 | 0.000142 | -0.20621 | 905.828 | 23.87413 | -4.14493 |
| **HCP-A** | rsfMRI | dMRI QA End | -0.39588 | 2.55E-12 | 6.79E-12 | -0.36216 | 213.2639 | 12.37301 | -0.2113 |
| **HCP-LS** | dMRI QA End | rsfMRI | -0.03084 | 0.23709 | 0.299482 | -0.13698 | 1211.166 | 30.39562 | -3.8613 |

| | | | | | | | | | |
|---|---|---|---|---|---|---|---|---|---|
| **HCP-LS** | rsfMRI | dMRI QA End | -0.12086 | 3.34E-06 | 6.69E-06 | -0.08159 | 264.5668 | 11.13931 | -0.0619 |

*Table S4. Cross-modality CPM model generalizability using the final CPM models.*

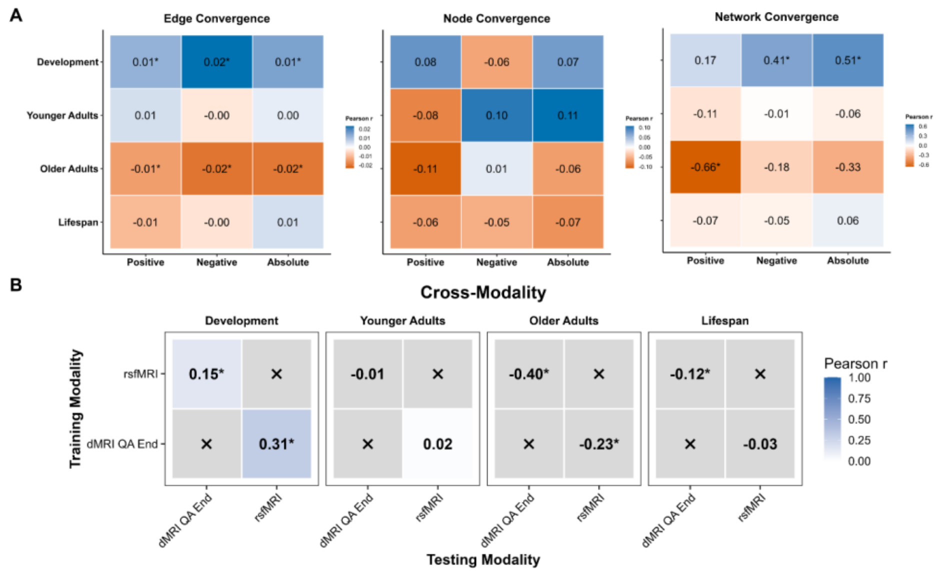

*Figure S1. CPM Structural-functional convergence analyses at the edge, node, and network levels. (A) Heatmaps show the correspondence between structural (dMRI QA End) and functional (rsfMRI) predictive networks across the Development, Younger Adult, Older Adult, and Lifespan cohorts. Edge-level convergence was quantified as the Pearson correlation between structural and functional predictive edge weights (left). Node-level convergence was quantified as the Pearson correlation between node predictive degrees (middle). Network-level convergence was quantified as the Pearson correlation between canonical network interaction matrices derived from the Shen atlas (right). Corresponding legends to the right of each. Separate analyses were performed for positive, negative, and absolute predictive networks. Cell values indicate Pearson correlation coefficients, and asterisks denote comparisons that remained significant after FDR correction (FDR < 0.05). (B) Heatmaps summarize the generalizability of final CPM age prediction models across modality evaluated through external validation/cross-sex prediction. Heatmap values represent the Pearson correlation coefficient (r) between predicted and chronological age when models were applied to independent datasets. Darker shading indicates stronger age prediction performance (higher Pearson r). Asterisks indicate predictions that remained significant after FDR correction (p < 0.05). Gray cells marked with × indicate comparisons that were not performed (e.g., identical training and testing modality).*

**Brain age prediction from individual canonical networks**

**Commented [MM1]:** Remove these tables for cpm and rcpm

| Between-Network | | | | | | | | |
|---|---|---|---|---|---|---|---|---|
| **Cohort** | **Modality** | **Network** | **Pearson r** | **pFDR** | **Spearman rho** | **MSE** | **MAE** | **q2** |

| | | | | | | | | |
|---|---|---|---|---|---|---|---|---|
| **HCP-D** | dMRI QA End | CBL-DMN | 0.278367 | 0.001257545 | 0.292211 | 13.21536 | 2.962237 | 0.068871 |
| **HCP-D** | dMRI QA End | CBL-FP | 0.435247 | 0.001257545 | 0.434931 | 11.58135 | 2.754535 | 0.184001 |
| **HCP-D** | dMRI QA End | CBL-MF | 0.255027 | 0.001257545 | 0.260263 | 13.54595 | 3.055303 | 0.045578 |
| **HCP-D** | dMRI QA End | CBL-Mot | 0.290837 | 0.001257545 | 0.295708 | 13.1706 | 2.970116 | 0.072025 |
| **HCP-D** | dMRI QA End | CBL-SAL | 0.464293 | 0.001257545 | 0.465993 | 11.21719 | 2.719318 | 0.209658 |
| **HCP-D** | dMRI QA End | CBL-SC | 0.344913 | 0.001257545 | 0.364762 | 12.77466 | 2.868875 | 0.099922 |
| **HCP-D** | dMRI QA End | CBL-VAs | 0.387931 | 0.001257545 | 0.38667 | 12.10873 | 2.889625 | 0.146842 |
| **HCP-D** | dMRI QA End | CBL-VII | 0.200347 | 0.001257545 | 0.188379 | 13.93894 | 3.103502 | 0.017889 |
| **HCP-D** | dMRI QA End | CBL-VI | 0.247101 | 0.002406932 | 0.246137 | 13.62247 | 3.028136 | 0.040187 |
| **HCP-D** | dMRI QA End | DMN-FP | 0.269107 | 0.001257545 | 0.286123 | 13.3191 | 2.993703 | 0.061562 |
| **HCP-D** | dMRI QA End | DMN-MF | 0.197668 | 0.001257545 | 0.199822 | 13.88523 | 3.114824 | 0.021673 |
| **HCP-D** | dMRI QA End | DMN-Mot | 0.096168 | 0.050063573 | 0.125179 | 14.63894 | 3.183081 | -0.03143 |
| **HCP-D** | dMRI QA End | DMN-SAL | 0.263544 | 0.001257545 | 0.262856 | 13.39697 | 3.015113 | 0.056075 |
| **HCP-D** | dMRI QA End | DMN-SC | 0.317062 | 0.001257545 | 0.329734 | 12.9454 | 2.944593 | 0.087892 |
| **HCP-D** | dMRI QA End | DMN-VAs | 0.124456 | 0.028345385 | 0.141267 | 14.08067 | 3.11497 | 0.007903 |
| **HCP-D** | dMRI QA End | DMN-VII | 0.149924 | 0.0078125 | 0.121738 | 14.01809 | 3.15402 | 0.012312 |
| **HCP-D** | dMRI QA End | DMN-VI | 0.133765 | 0.013000144 | 0.138661 | 14.0481 | 3.119227 | 0.010198 |
| **HCP-D** | dMRI QA End | FP-MF | 0.350189 | 0.001257545 | 0.335732 | 12.59401 | 2.975177 | 0.11265 |
| **HCP-D** | dMRI QA End | FP-Mot | 0.305557 | 0.001257545 | 0.312953 | 13.07657 | 2.985315 | 0.07865 |
| **HCP-D** | dMRI QA End | FP-SAL | 0.45505 | 0.001257545 | 0.463978 | 11.32379 | 2.773636 | 0.202147 |
| **HCP-D** | dMRI QA End | FP-SC | 0.389431 | 0.001257545 | 0.391331 | 12.17617 | 2.838744 | 0.142091 |
| **HCP-D** | dMRI QA End | FP-VAs | 0.286494 | 0.001257545 | 0.289682 | 13.16401 | 3.038139 | 0.07249 |
| **HCP-D** | dMRI QA End | FP-VII | 0.259613 | 0.001257545 | 0.247371 | 13.38481 | 3.058598 | 0.056932 |

| | | | | | | | | |
|---|---|---|---|---|---|---|---|---|
| **HCP-D** | dMRI QA End | FP-VI | 0.331796 | 0.001257545 | 0.342481 | 12.75163 | 2.921089 | 0.101545 |
| **HCP-D** | dMRI QA End | MF-Mot | 0.30953 | 0.001257545 | 0.317235 | 13.0796 | 2.972362 | 0.078437 |
| **HCP-D** | dMRI QA End | MF-SAL | 0.419055 | 0.001257545 | 0.422758 | 11.80075 | 2.773603 | 0.168542 |
| **HCP-D** | dMRI QA End | MF-SC | 0.289219 | 0.001257545 | 0.301543 | 13.22444 | 2.972732 | 0.068231 |
| **HCP-D** | dMRI QA End | MF-VAs | 0.222045 | 0.001257545 | 0.235812 | 13.76662 | 3.0394 | 0.030031 |
| **HCP-D** | dMRI QA End | MF-VII | 0.17392 | 0.004624634 | 0.179553 | 13.97357 | 3.106981 | 0.01545 |
| **HCP-D** | dMRI QA End | MF-VI | 0.284516 | 0.001257545 | 0.287046 | 13.19078 | 3.004204 | 0.070603 |
| **HCP-D** | dMRI QA End | Mot-SAL | 0.37232 | 0.001257545 | 0.380414 | 12.33705 | 2.875295 | 0.130755 |
| **HCP-D** | dMRI QA End | Mot-SC | 0.352369 | 0.001257545 | 0.360324 | 12.57396 | 2.898872 | 0.114063 |
| **HCP-D** | dMRI QA End | Mot-VAs | 0.280317 | 0.001257545 | 0.297383 | 13.14863 | 3.009453 | 0.073573 |
| **HCP-D** | dMRI QA End | Mot-VII | 0.342395 | 0.001257545 | 0.353577 | 12.59152 | 2.923984 | 0.112826 |
| **HCP-D** | dMRI QA End | Mot-VI | 0.364337 | 0.001257545 | 0.346714 | 12.48237 | 2.892787 | 0.120516 |
| **HCP-D** | dMRI QA End | SAL-SC | 0.343447 | 0.001257545 | 0.340669 | 12.79549 | 2.971255 | 0.098455 |
| **HCP-D** | dMRI QA End | SAL-VAs | 0.34903 | 0.001257545 | 0.334864 | 12.58954 | 2.924541 | 0.112966 |
| **HCP-D** | dMRI QA End | SAL-VII | 0.221674 | 0.001257545 | 0.200778 | 13.71296 | 3.077185 | 0.033812 |
| **HCP-D** | dMRI QA End | SAL-VI | 0.33405 | 0.001257545 | 0.31129 | 12.71769 | 2.933005 | 0.103936 |
| **HCP-D** | dMRI QA End | SC-VAs | 0.247281 | 0.001257545 | 0.257739 | 13.53592 | 3.050239 | 0.046285 |
| **HCP-D** | dMRI QA End | SC-VII | 0.137438 | 0.008798084 | 0.146987 | 14.3478 | 3.104183 | -0.01092 |
| **HCP-D** | dMRI QA End | SC-VI | 0.205172 | 0.001257545 | 0.208339 | 14.06379 | 3.124477 | 0.009093 |
| **HCP-D** | dMRI QA End | VAs-VII | 0.222048 | 0.002406932 | 0.234175 | 13.58556 | 3.080851 | 0.042788 |
| **HCP-D** | dMRI QA End | VAs-VI | 0.270152 | 0.001257545 | 0.271237 | 13.28309 | 3.028315 | 0.064099 |
| **HCP-D** | dMRI QA End | VI-VII | 0.049069 | 0.178571429 | 0.045279 | 14.71086 | 3.162528 | -0.0365 |
| **HCP-YA** | dMRI QA End | CBL-DMN | 0.065466 | 0.145716353 | 0.06977 | 14.06774 | 3.13288 | -0.0359 |

| | | | | | | | | |
|---|---|---|---|---|---|---|---|---|
| **HCP-YA** | dMRI QA End | CBL-FP | 0.145728 | 0.012260467 | 0.137875 | 13.78202 | 3.088395 | -0.01486 |
| **HCP-YA** | dMRI QA End | CBL-MF | 0.082188 | 0.091783217 | 0.066728 | 13.9665 | 3.131816 | -0.02844 |
| **HCP-YA** | dMRI QA End | CBL-Mot | 0.161946 | 0.01125 | 0.151492 | 13.66714 | 3.056279 | -0.0064 |
| **HCP-YA** | dMRI QA End | CBL-SAL | 0.134455 | 0.012260467 | 0.13065 | 13.83186 | 3.102196 | -0.01853 |
| **HCP-YA** | dMRI QA End | CBL-SC | 0.163145 | 0.012260467 | 0.161591 | 13.69955 | 3.042406 | -0.00879 |
| **HCP-YA** | dMRI QA End | CBL-VAs | 0.127968 | 0.014985015 | 0.119391 | 13.69308 | 3.082991 | -0.00831 |
| **HCP-YA** | dMRI QA End | CBL-VII | -0.07778 | 0.785785786 | -0.06655 | 14.22474 | 3.158449 | -0.04746 |
| **HCP-YA** | dMRI QA End | CBL-VI | 0.083554 | 0.100699301 | 0.076492 | 14.01025 | 3.135922 | -0.03167 |
| **HCP-YA** | dMRI QA End | DMN-FP | 0.059681 | 0.171391109 | 0.063388 | 13.952 | 3.154286 | -0.02738 |
| **HCP-YA** | dMRI QA End | DMN-MF | -0.03007 | 0.583440583 | -0.05926 | 14.09378 | 3.135964 | -0.03782 |
| **HCP-YA** | dMRI QA End | DMN-Mot | 0.079009 | 0.091783217 | 0.069436 | 14.04734 | 3.128395 | -0.0344 |
| **HCP-YA** | dMRI QA End | DMN-SAL | -0.00689 | 0.539310238 | -0.00042 | 14.29712 | 3.147547 | -0.05279 |
| **HCP-YA** | dMRI QA End | DMN-SC | 0.022835 | 0.334665335 | 0.004507 | 14.09886 | 3.143406 | -0.03819 |
| **HCP-YA** | dMRI QA End | DMN-VAs | 0.05351 | 0.171391109 | 0.052091 | 13.94036 | 3.141742 | -0.00693 |
| **HCP-YA** | dMRI QA End | DMN-VII | 0.08104 | 0.091783217 | 0.08662 | 13.53401 | 3.088968 | 0.003403 |
| **HCP-YA** | dMRI QA End | DMN-VI | 0.133393 | 0.012844299 | 0.137287 | 13.57451 | 3.082355 | 0.000421 |
| **HCP-YA** | dMRI QA End | FP-MF | 0.007006 | 0.487215487 | -0.00921 | 14.73002 | 3.205287 | -0.08467 |
| **HCP-YA** | dMRI QA End | FP-Mot | 0.161045 | 0.012260467 | 0.159396 | 13.49287 | 3.060054 | 0.006432 |
| **HCP-YA** | dMRI QA End | FP-SAL | 0.12157 | 0.027472527 | 0.098877 | 14.03132 | 3.095442 | -0.03322 |
| **HCP-YA** | dMRI QA End | FP-SC | 0.12021 | 0.012260467 | 0.1261 | 13.96751 | 3.087288 | -0.02852 |
| **HCP-YA** | dMRI QA End | FP-VAs | 0.030218 | 0.310718693 | 0.026351 | 14.22353 | 3.126318 | -0.04737 |
| **HCP-YA** | dMRI QA End | FP-VII | -0.03788 | 0.640609391 | -0.02882 | 14.09271 | 3.155407 | -0.03774 |
| **HCP-YA** | dMRI QA End | FP-VI | 0.20707 | 0.01125 | 0.189242 | 13.24869 | 3.020707 | 0.024413 |

| | | | | | | | | |
|---|---|---|---|---|---|---|---|---|
| **HCP-YA** | dMRI QA End | MF-Mot | -0.00856 | 0.550699301 | -0.00857 | 14.42008 | 3.179154 | -0.06184 |
| **HCP-YA** | dMRI QA End | MF-SAL | 0.064857 | 0.145716353 | 0.065384 | 14.05629 | 3.143199 | -0.03506 |
| **HCP-YA** | dMRI QA End | MF-SC | 0.080594 | 0.103230103 | 0.077996 | 14.01646 | 3.107245 | -0.03212 |
| **HCP-YA** | dMRI QA End | MF-VAs | 0.052329 | 0.171391109 | 0.05865 | 13.72158 | 3.093405 | -0.01041 |
| **HCP-YA** | dMRI QA End | MF-VII | -0.01798 | 0.539310238 | -0.00721 | 13.87766 | 3.11876 | -0.0219 |
| **HCP-YA** | dMRI QA End | MF-VI | 0.128248 | 0.016858142 | 0.124053 | 13.65603 | 3.087813 | -0.00558 |
| **HCP-YA** | dMRI QA End | Mot-SAL | 0.164356 | 0.012260467 | 0.165346 | 13.70488 | 3.042775 | -0.00918 |
| **HCP-YA** | dMRI QA End | Mot-SC | 0.142049 | 0.012844299 | 0.134394 | 13.73954 | 3.067582 | -0.01173 |
| **HCP-YA** | dMRI QA End | Mot-VAs | 0.08508 | 0.091783217 | 0.092884 | 13.88812 | 3.092353 | -0.02267 |
| **HCP-YA** | dMRI QA End | Mot-VII | 0.052361 | 0.192080647 | 0.056609 | 14.0406 | 3.093424 | -0.0339 |
| **HCP-YA** | dMRI QA End | Mot-VI | 0.02983 | 0.321107464 | 0.024096 | 14.10551 | 3.146136 | -0.03868 |
| **HCP-YA** | dMRI QA End | SAL-SC | 0.141974 | 0.018510901 | 0.152732 | 13.67944 | 3.071653 | -0.00731 |
| **HCP-YA** | dMRI QA End | SAL-VAs | 0.175164 | 0.012260467 | 0.157071 | 13.38562 | 3.035748 | 0.01433 |
| **HCP-YA** | dMRI QA End | SAL-VII | 0.144263 | 0.01125 | 0.139926 | 13.35188 | 3.048915 | 0.016814 |
| **HCP-YA** | dMRI QA End | SAL-VI | 0.120513 | 0.02839266 | 0.111128 | 13.85991 | 3.098681 | -0.02059 |
| **HCP-YA** | dMRI QA End | SC-VAs | 0.152488 | 0.01125 | 0.143795 | 13.61913 | 3.073501 | -0.00287 |
| **HCP-YA** | dMRI QA End | SC-VII | -0.03848 | 0.600697674 | -0.03214 | 13.93769 | 3.129361 | -0.02632 |
| **HCP-YA** | dMRI QA End | SC-VI | 0.134424 | 0.012844299 | 0.131108 | 13.59646 | 3.094765 | -0.0012 |
| **HCP-YA** | dMRI QA End | VAs-VII | 0.09548 | 0.051853707 | 0.085322 | 13.51034 | 3.08923 | 0.005146 |
| **HCP-YA** | dMRI QA End | VAs-VI | 0.075472 | 0.103230103 | 0.082507 | 13.72773 | 3.084847 | -0.01086 |
| **HCP-YA** | dMRI QA End | VI-VII | -0.03414 | 0.567054712 | -0.05442 | 13.8995 | 3.126872 | -0.02351 |
| **HCP-A** | dMRI QA End | CBL-DMN | 0.386363 | 0.001278119 | 0.34324 | 150.7744 | 9.915539 | 0.143629 |
| **HCP-A** | dMRI QA End | CBL-FP | 0.525344 | 0.001278119 | 0.49333 | 127.7309 | 9.170274 | 0.274512 |

| | | | | | | | | |
|---|---|---|---|---|---|---|---|---|
| **HCP-A** | dMRI QA End | CBL-MF | 0.421317 | 0.001278119 | 0.40945 | 146.0236 | 9.86676 | 0.170613 |
| **HCP-A** | dMRI QA End | CBL-Mot | 0.49763 | 0.001278119 | 0.444772 | 133.0101 | 9.370207 | 0.244527 |
| **HCP-A** | dMRI QA End | CBL-SAL | 0.516106 | 0.001278119 | 0.492338 | 129.6977 | 9.138943 | 0.263341 |
| **HCP-A** | dMRI QA End | CBL-SC | 0.606305 | 0.001278119 | 0.577151 | 111.4671 | 8.402173 | 0.366888 |
| **HCP-A** | dMRI QA End | CBL-VAs | 0.501837 | 0.001278119 | 0.488212 | 132.2591 | 9.293078 | 0.248793 |
| **HCP-A** | dMRI QA End | CBL-VII | 0.295897 | 0.001278119 | 0.283579 | 161.9232 | 10.35941 | 0.080306 |
| **HCP-A** | dMRI QA End | CBL-VI | 0.417172 | 0.001278119 | 0.377381 | 146.7912 | 9.84745 | 0.166253 |
| **HCP-A** | dMRI QA End | DMN-FP | 0.344445 | 0.001278119 | 0.33763 | 156.5854 | 10.15276 | 0.110624 |
| **HCP-A** | dMRI QA End | DMN-MF | 0.296889 | 0.002261307 | 0.291777 | 162.5475 | 10.47358 | 0.07676 |
| **HCP-A** | dMRI QA End | DMN-Mot | 0.307717 | 0.002261307 | 0.293813 | 160.9121 | 10.20199 | 0.086049 |
| **HCP-A** | dMRI QA End | DMN-SAL | 0.345274 | 0.001278119 | 0.285029 | 155.6025 | 10.13938 | 0.116207 |
| **HCP-A** | dMRI QA End | DMN-SC | 0.442991 | 0.002261307 | 0.408496 | 141.7902 | 9.753187 | 0.194658 |
| **HCP-A** | dMRI QA End | DMN-VAs | 0.286647 | 0.002315529 | 0.275842 | 162.3111 | 10.25623 | 0.078103 |
| **HCP-A** | dMRI QA End | DMN-VII | 0.166029 | 0.009333437 | 0.191845 | 172.0552 | 10.82824 | 0.022758 |
| **HCP-A** | dMRI QA End | DMN-VI | 0.271882 | 0.004436557 | 0.266241 | 163.7075 | 10.36187 | 0.070172 |
| **HCP-A** | dMRI QA End | FP-MF | 0.465519 | 0.001278119 | 0.458369 | 138.4962 | 9.639252 | 0.213367 |
| **HCP-A** | dMRI QA End | FP-Mot | 0.48124 | 0.001278119 | 0.465815 | 136.0061 | 9.600396 | 0.22751 |
| **HCP-A** | dMRI QA End | FP-SAL | 0.460048 | 0.001278119 | 0.439209 | 139.4053 | 9.528026 | 0.208204 |
| **HCP-A** | dMRI QA End | FP-SC | 0.464445 | 0.001278119 | 0.451668 | 138.5075 | 9.545084 | 0.213303 |
| **HCP-A** | dMRI QA End | FP-VAs | 0.503969 | 0.001278119 | 0.461985 | 131.5267 | 9.334251 | 0.252953 |
| **HCP-A** | dMRI QA End | FP-VII | 0.339572 | 0.001278119 | 0.281212 | 156.7335 | 10.27574 | 0.109783 |
| **HCP-A** | dMRI QA End | FP-VI | 0.29991 | 0.001278119 | 0.292104 | 163.3962 | 10.43411 | 0.07194 |
| **HCP-A** | dMRI QA End | MF-Mot | 0.407735 | 0.001278119 | 0.404977 | 147.694 | 10.0972 | 0.161126 |

| | | | | | | | | |
|---|---|---|---|---|---|---|---|---|
| **HCP-A** | dMRI QA End | MF-SAL | 0.481813 | 0.001278119 | 0.434867 | 135.9733 | 9.553986 | 0.227697 |
| **HCP-A** | dMRI QA End | MF-SC | 0.327407 | 0.001278119 | 0.29276 | 160.2688 | 10.48078 | 0.089703 |
| **HCP-A** | dMRI QA End | MF-VAs | 0.485307 | 0.001278119 | 0.45719 | 134.8759 | 9.500509 | 0.23393 |
| **HCP-A** | dMRI QA End | MF-VII | 0.167982 | 0.01575042 | 0.141 | 173.1136 | 10.83058 | 0.016747 |
| **HCP-A** | dMRI QA End | MF-VI | 0.337187 | 0.001278119 | 0.299621 | 158.3092 | 10.39168 | 0.100833 |
| **HCP-A** | dMRI QA End | Mot-SAL | 0.4613 | 0.001278119 | 0.421079 | 139.3314 | 9.73571 | 0.208623 |
| **HCP-A** | dMRI QA End | Mot-SC | 0.460005 | 0.001278119 | 0.430742 | 139.2253 | 9.534685 | 0.209226 |
| **HCP-A** | dMRI QA End | Mot-VAs | 0.376743 | 0.001278119 | 0.33001 | 151.7014 | 10.00133 | 0.138364 |
| **HCP-A** | dMRI QA End | Mot-VII | 0.333509 | 0.001278119 | 0.307427 | 158.6383 | 10.33986 | 0.098964 |
| **HCP-A** | dMRI QA End | Mot-VI | 0.377354 | 0.001278119 | 0.353758 | 152.2373 | 10.10548 | 0.135321 |
| **HCP-A** | dMRI QA End | SAL-SC | 0.485312 | 0.001278119 | 0.451503 | 135.0984 | 9.511343 | 0.232666 |
| **HCP-A** | dMRI QA End | SAL-VAs | 0.474998 | 0.001278119 | 0.459518 | 137.0054 | 9.376115 | 0.221835 |
| **HCP-A** | dMRI QA End | SAL-VII | 0.307421 | 0.002261307 | 0.297325 | 161.0393 | 10.35287 | 0.085327 |
| **HCP-A** | dMRI QA End | SAL-VI | 0.37502 | 0.001278119 | 0.353003 | 152.8328 | 9.98015 | 0.131938 |
| **HCP-A** | dMRI QA End | SC-VAs | 0.448929 | 0.001278119 | 0.425225 | 141.2339 | 9.587007 | 0.197818 |
| **HCP-A** | dMRI QA End | SC-VII | 0.332897 | 0.001278119 | 0.352181 | 158.0449 | 10.14321 | 0.102334 |
| **HCP-A** | dMRI QA End | SC-VI | 0.377314 | 0.001278119 | 0.357414 | 153.1142 | 10.05933 | 0.13034 |
| **HCP-A** | dMRI QA End | VAs-VII | 0.310257 | 0.001278119 | 0.273333 | 160.51 | 10.32068 | 0.088333 |
| **HCP-A** | dMRI QA End | VAs-VI | 0.251146 | 0.001278119 | 0.23015 | 168.3972 | 10.68553 | 0.043535 |
| **HCP-A** | dMRI QA End | VI-VII | 0.045 | 0.182939363 | 0.014052 | 183.8903 | 11.0298 | -0.04446 |
| **HCP-LS** | dMRI QA End | CBL-DMN | 0.515684 | 0.001075552 | 0.350049 | 182.9001 | 10.18587 | 0.265888 |
| **HCP-LS** | dMRI QA End | CBL-FP | 0.539128 | 0.001075552 | 0.355172 | 176.7561 | 10.27455 | 0.290548 |
| **HCP-LS** | dMRI QA End | CBL-MF | 0.417532 | 0.001075552 | 0.194301 | 205.7789 | 10.86294 | 0.174058 |

| | | | | | | | | |
|---|---|---|---|---|---|---|---|---|
| **HCP-LS** | dMRI QA End | CBL-Mot | 0.518685 | 0.001075552 | 0.275007 | 182.1435 | 10.46181 | 0.268924 |
| **HCP-LS** | dMRI QA End | CBL-SAL | 0.47676 | 0.001075552 | 0.298953 | 192.6215 | 10.53795 | 0.226868 |
| **HCP-LS** | dMRI QA End | CBL-SC | 0.623749 | 0.001075552 | 0.380042 | 152.2282 | 9.7783 | 0.388996 |
| **HCP-LS** | dMRI QA End | CBL-VAs | 0.54527 | 0.001075552 | 0.298633 | 175.0801 | 10.37333 | 0.297275 |
| **HCP-LS** | dMRI QA End | CBL-VII | 0.395121 | 0.001075552 | 0.217663 | 210.5966 | 10.93605 | 0.154721 |
| **HCP-LS** | dMRI QA End | CBL-VI | 0.396672 | 0.001075552 | 0.260174 | 210.2014 | 10.91027 | 0.156307 |
| **HCP-LS** | dMRI QA End | DMN-FP | 0.338535 | 0.001075552 | 0.095281 | 220.6132 | 11.09542 | 0.114517 |
| **HCP-LS** | dMRI QA End | DMN-MF | 0.352324 | 0.001075552 | 0.12641 | 218.3314 | 11.06042 | 0.123676 |
| **HCP-LS** | dMRI QA End | DMN-Mot | 0.428033 | 0.001075552 | 0.332948 | 203.5824 | 10.54342 | 0.182874 |
| **HCP-LS** | dMRI QA End | DMN-SAL | 0.373041 | 0.001075552 | 0.208134 | 214.7073 | 10.82071 | 0.138222 |
| **HCP-LS** | dMRI QA End | DMN-SC | 0.499271 | 0.001075552 | 0.367818 | 187.0879 | 10.21626 | 0.249079 |
| **HCP-LS** | dMRI QA End | DMN-VAs | 0.312519 | 0.00111166 | 0.119279 | 224.8895 | 11.10729 | 0.097353 |
| **HCP-LS** | dMRI QA End | DMN-VII | 0.34954 | 0.001182033 | 0.28195 | 218.8413 | 10.8427 | 0.121629 |
| **HCP-LS** | dMRI QA End | DMN-VI | 0.168812 | 0.001075552 | 0.031224 | 243.3462 | 11.41368 | 0.023273 |
| **HCP-LS** | dMRI QA End | FP-MF | 0.504817 | 0.001075552 | 0.247627 | 185.6797 | 10.59642 | 0.254731 |
| **HCP-LS** | dMRI QA End | FP-Mot | 0.515948 | 0.001075552 | 0.378633 | 182.8598 | 10.2969 | 0.266049 |
| **HCP-LS** | dMRI QA End | FP-SAL | 0.498226 | 0.001075552 | 0.328705 | 187.328 | 10.42354 | 0.248115 |
| **HCP-LS** | dMRI QA End | FP-SC | 0.512123 | 0.001075552 | 0.272767 | 183.8502 | 10.52352 | 0.262074 |
| **HCP-LS** | dMRI QA End | FP-VAs | 0.54326 | 0.001075552 | 0.38268 | 175.6554 | 10.18454 | 0.294966 |
| **HCP-LS** | dMRI QA End | FP-VII | 0.412004 | 0.001075552 | 0.252221 | 206.9623 | 10.87431 | 0.169308 |
| **HCP-LS** | dMRI QA End | FP-VI | 0.489691 | 0.001075552 | 0.306896 | 189.4529 | 10.49316 | 0.239586 |
| **HCP-LS** | dMRI QA End | MF-Mot | 0.468995 | 0.001075552 | 0.24725 | 194.3864 | 10.64093 | 0.219784 |
| **HCP-LS** | dMRI QA End | MF-SAL | 0.517022 | 0.001075552 | 0.268428 | 182.5612 | 10.35792 | 0.267248 |

| | | | | | | | | |
|---|---|---|---|---|---|---|---|---|
| **HCP-LS** | dMRI QA End | MF-SC | 0.501779 | 0.001075552 | 0.361931 | 186.4397 | 10.37027 | 0.251681 |
| **HCP-LS** | dMRI QA End | MF-VAs | 0.440197 | 0.001075552 | 0.228488 | 200.9156 | 10.74715 | 0.193578 |
| **HCP-LS** | dMRI QA End | MF-VII | 0.344789 | 0.001075552 | 0.120336 | 219.615 | 11.11013 | 0.118524 |
| **HCP-LS** | dMRI QA End | MF-VI | 0.427216 | 0.001075552 | 0.208462 | 203.7277 | 10.85365 | 0.182291 |
| **HCP-LS** | dMRI QA End | Mot-SAL | 0.553645 | 0.001075552 | 0.376849 | 172.7987 | 10.20662 | 0.306432 |
| **HCP-LS** | dMRI QA End | Mot-SC | 0.550373 | 0.001075552 | 0.335007 | 173.685 | 10.20901 | 0.302874 |
| **HCP-LS** | dMRI QA End | Mot-VAs | 0.475169 | 0.001075552 | 0.339666 | 192.9616 | 10.47241 | 0.225503 |
| **HCP-LS** | dMRI QA End | Mot-VII | 0.374259 | 0.001075552 | 0.201944 | 214.425 | 10.91665 | 0.139355 |
| **HCP-LS** | dMRI QA End | Mot-VI | 0.44639 | 0.001075552 | 0.283876 | 199.6351 | 10.69352 | 0.198718 |
| **HCP-LS** | dMRI QA End | SAL-SC | 0.468022 | 0.001075552 | 0.251799 | 194.6184 | 10.70314 | 0.218853 |
| **HCP-LS** | dMRI QA End | SAL-VAs | 0.607732 | 0.001075552 | 0.421028 | 157.1464 | 9.637813 | 0.369256 |
| **HCP-LS** | dMRI QA End | SAL-VII | 0.472062 | 0.001075552 | 0.369102 | 193.6866 | 10.41003 | 0.222593 |
| **HCP-LS** | dMRI QA End | SAL-VI | 0.549518 | 0.001075552 | 0.3785 | 173.95 | 10.06985 | 0.301811 |
| **HCP-LS** | dMRI QA End | SC-VAs | 0.54351 | 0.001075552 | 0.302245 | 175.5735 | 10.31549 | 0.295294 |
| **HCP-LS** | dMRI QA End | SC-VII | 0.362486 | 0.001075552 | 0.179281 | 216.4449 | 10.99957 | 0.131248 |
| **HCP-LS** | dMRI QA End | SC-VI | 0.440625 | 0.001075552 | 0.303425 | 200.9353 | 10.69687 | 0.193499 |
| **HCP-LS** | dMRI QA End | VAs-VII | 0.350515 | 0.001075552 | 0.188602 | 218.5673 | 10.97235 | 0.122729 |
| **HCP-LS** | dMRI QA End | VAs-VI | 0.388545 | 0.001075552 | 0.258843 | 211.8818 | 10.79214 | 0.149563 |
| **HCP-LS** | dMRI QA End | VI-VII | 0.274387 | 0.001075552 | 0.027098 | 230.5727 | 11.35412 | 0.074542 |
| **HCP-D** | rsfMRI | CBL-DMN | 0.676604 | 0.000999001 | 0.70688 | 7.696011 | 2.237619 | 0.457754 |
| **HCP-D** | rsfMRI | CBL-FP | 0.701384 | 0.000999001 | 0.71915 | 7.210974 | 2.124448 | 0.491929 |
| **HCP-D** | rsfMRI | CBL-MF | 0.661141 | 0.000999001 | 0.679806 | 7.989474 | 2.232407 | 0.437077 |
| **HCP-D** | rsfMRI | CBL-Mot | 0.66883 | 0.000999001 | 0.685334 | 7.844481 | 2.242604 | 0.447293 |
| **HCP-D** | rsfMRI | CBL-SAL | 0.646303 | 0.000999001 | 0.674487 | 8.266055 | 2.275056 | 0.41759 |
| **HCP-D** | rsfMRI | CBL-SC | 0.713493 | 0.000999001 | 0.735553 | 6.9679 | 2.088746 | 0.509055 |
| **HCP-D** | rsfMRI | CBL-VAs | 0.654895 | 0.000999001 | 0.66722 | 8.106046 | 2.262923 | 0.428864 |

| | | | | | | | | |
|---|---|---|---|---|---|---|---|---|
| **HCP-D** | rsfMRI | CBL-VII | 0.655972 | 0.000999001 | 0.667932 | 8.085916 | 2.2598 | 0.430282 |
| **HCP-D** | rsfMRI | CBL-VI | 0.635968 | 0.000999001 | 0.650129 | 8.454949 | 2.334941 | 0.404281 |
| **HCP-D** | rsfMRI | DMN-FP | 0.629151 | 0.000999001 | 0.634742 | 8.580508 | 2.362077 | 0.395434 |
| **HCP-D** | rsfMRI | DMN-MF | 0.591296 | 0.000999001 | 0.604145 | 9.248485 | 2.440005 | 0.34837 |
| **HCP-D** | rsfMRI | DMN-Mot | 0.621634 | 0.000999001 | 0.632952 | 8.713635 | 2.421053 | 0.386054 |
| **HCP-D** | rsfMRI | DMN-SAL | 0.670955 | 0.000999001 | 0.704977 | 7.804299 | 2.224055 | 0.450124 |
| **HCP-D** | rsfMRI | DMN-SC | 0.709811 | 0.000999001 | 0.724285 | 7.042257 | 2.153299 | 0.503816 |
| **HCP-D** | rsfMRI | DMN-VAs | 0.595854 | 0.000999001 | 0.612167 | 9.160139 | 2.451761 | 0.354594 |
| **HCP-D** | rsfMRI | DMN-VII | 0.537819 | 0.000999001 | 0.559071 | 10.12025 | 2.571962 | 0.286947 |
| **HCP-D** | rsfMRI | DMN-VI | 0.612261 | 0.000999001 | 0.627264 | 8.892328 | 2.439659 | 0.373464 |
| **HCP-D** | rsfMRI | FP-MF | 0.677728 | 0.000999001 | 0.688232 | 7.67666 | 2.2148 | 0.459117 |
| **HCP-D** | rsfMRI | FP-Mot | 0.700508 | 0.000999001 | 0.701697 | 7.228296 | 2.171709 | 0.490708 |
| **HCP-D** | rsfMRI | FP-SAL | 0.708497 | 0.000999001 | 0.734635 | 7.06875 | 2.109463 | 0.50195 |
| **HCP-D** | rsfMRI | FP-SC | 0.742764 | 0.000999001 | 0.753238 | 6.365584 | 2.033997 | 0.551493 |
| **HCP-D** | rsfMRI | FP-VAs | 0.712966 | 0.000999001 | 0.725684 | 6.978833 | 2.135708 | 0.508285 |
| **HCP-D** | rsfMRI | FP-VII | 0.657327 | 0.000999001 | 0.659886 | 8.063099 | 2.269084 | 0.43189 |
| **HCP-D** | rsfMRI | FP-VI | 0.662797 | 0.000999001 | 0.67426 | 7.959772 | 2.27189 | 0.43917 |
| **HCP-D** | rsfMRI | MF-Mot | 0.64689 | 0.000999001 | 0.655112 | 8.257674 | 2.294362 | 0.41818 |
| **HCP-D** | rsfMRI | MF-SAL | 0.670099 | 0.000999001 | 0.692032 | 7.820635 | 2.171182 | 0.448973 |
| **HCP-D** | rsfMRI | MF-SC | 0.705622 | 0.000999001 | 0.717347 | 7.12623 | 2.108949 | 0.4979 |
| **HCP-D** | rsfMRI | MF-VAs | 0.64575 | 0.000999001 | 0.655798 | 8.276261 | 2.271679 | 0.416871 |
| **HCP-D** | rsfMRI | MF-VII | 0.547651 | 0.000999001 | 0.561693 | 9.973166 | 2.543486 | 0.29731 |
| **HCP-D** | rsfMRI | MF-VI | 0.57385 | 0.000999001 | 0.581295 | 9.538329 | 2.44107 | 0.327948 |
| **HCP-D** | rsfMRI | Mot-SAL | 0.663759 | 0.000999001 | 0.688139 | 7.940963 | 2.239215 | 0.440495 |
| **HCP-D** | rsfMRI | Mot-SC | 0.720018 | 0.000999001 | 0.725582 | 6.835241 | 2.105862 | 0.518402 |
| **HCP-D** | rsfMRI | Mot-VAs | 0.604879 | 0.000999001 | 0.606968 | 9.008858 | 2.449723 | 0.365253 |
| **HCP-D** | rsfMRI | Mot-VII | 0.600761 | 0.000999001 | 0.609953 | 9.077895 | 2.455624 | 0.360389 |
| **HCP-D** | rsfMRI | Mot-VI | 0.611341 | 0.000999001 | 0.622488 | 8.900208 | 2.459436 | 0.372909 |
| **HCP-D** | rsfMRI | SAL-SC | 0.706027 | 0.000999001 | 0.722756 | 7.118374 | 2.104374 | 0.498453 |
| **HCP-D** | rsfMRI | SAL-VAs | 0.62835 | 0.000999001 | 0.653911 | 8.592262 | 2.337913 | 0.394606 |
| **HCP-D** | rsfMRI | SAL-VII | 0.625999 | 0.000999001 | 0.638981 | 8.636272 | 2.359885 | 0.391505 |
| **HCP-D** | rsfMRI | SAL-VI | 0.64024 | 0.000999001 | 0.647135 | 8.376914 | 2.279474 | 0.409779 |
| **HCP-D** | rsfMRI | SC-VAs | 0.71574 | 0.000999001 | 0.722198 | 6.922797 | 2.10577 | 0.512233 |
| **HCP-D** | rsfMRI | SC-VII | 0.63853 | 0.000999001 | 0.638519 | 8.408345 | 2.329696 | 0.407564 |
| **HCP-D** | rsfMRI | SC-VI | 0.671481 | 0.000999001 | 0.678884 | 7.793776 | 2.219576 | 0.450866 |
| **HCP-D** | rsfMRI | VAs-VII | 0.569931 | 0.000999001 | 0.568518 | 9.590459 | 2.493549 | 0.324275 |
| **HCP-D** | rsfMRI | VAs-VI | 0.554424 | 0.000999001 | 0.563531 | 9.850892 | 2.54495 | 0.305925 |
| **HCP-D** | rsfMRI | VI-VII | 0.490259 | 0.000999001 | 0.509135 | 10.82807 | 2.70084 | 0.237075 |
| **HCP-YA** | rsfMRI | CBL-DMN | 0.190071 | 0.001096465 | 0.204019 | 13.45516 | 3.051733 | 0.009209 |

| | | | | | | | | |
|---|---|---|---|---|---|---|---|---|
| **HCP-YA** | rsfMRI | CBL-FP | 0.194925 | 0.001096465 | 0.198783 | 13.43481 | 3.014327 | 0.010707 |
| **HCP-YA** | rsfMRI | CBL-MF | 0.214879 | 0.001096465 | 0.211454 | 13.12789 | 2.981722 | 0.033308 |
| **HCP-YA** | rsfMRI | CBL-Mot | 0.17807 | 0.001096465 | 0.190196 | 13.30931 | 3.005416 | 0.019949 |
| **HCP-YA** | rsfMRI | CBL-SAL | 0.166014 | 0.001096465 | 0.175901 | 13.46426 | 3.030233 | 0.008539 |
| **HCP-YA** | rsfMRI | CBL-SC | 0.194662 | 0.001096465 | 0.198875 | 13.48194 | 3.012873 | 0.007237 |
| **HCP-YA** | rsfMRI | CBL-VAs | 0.170812 | 0.001096465 | 0.182101 | 13.30367 | 3.008344 | 0.020365 |
| **HCP-YA** | rsfMRI | CBL-VII | 0.152297 | 0.001096465 | 0.159323 | 13.49126 | 3.039997 | 0.006551 |
| **HCP-YA** | rsfMRI | CBL-VI | 0.177048 | 0.001096465 | 0.190174 | 13.41638 | 3.023555 | 0.012065 |
| **HCP-YA** | rsfMRI | DMN-FP | 0.21459 | 0.001096465 | 0.209353 | 13.41256 | 3.02304 | 0.012346 |
| **HCP-YA** | rsfMRI | DMN-MF | 0.250837 | 0.001096465 | 0.251976 | 12.97584 | 2.988753 | 0.044505 |
| **HCP-YA** | rsfMRI | DMN-Mot | 0.196323 | 0.001096465 | 0.19987 | 13.45216 | 3.013829 | 0.00943 |
| **HCP-YA** | rsfMRI | DMN-SAL | 0.198864 | 0.001096465 | 0.19466 | 13.5241 | 3.047587 | 0.004133 |
| **HCP-YA** | rsfMRI | DMN-SC | 0.227112 | 0.001096465 | 0.232474 | 13.2916 | 2.99043 | 0.021253 |
| **HCP-YA** | rsfMRI | DMN-VAs | 0.216132 | 0.001096465 | 0.233971 | 13.18012 | 3.007099 | 0.029462 |
| **HCP-YA** | rsfMRI | DMN-VII | 0.06419 | 0.115884116 | 0.080299 | 14.1834 | 3.15486 | -0.04442 |
| **HCP-YA** | rsfMRI | DMN-VI | 0.192951 | 0.001096465 | 0.211597 | 13.45129 | 3.052612 | 0.009494 |
| **HCP-YA** | rsfMRI | FP-MF | 0.19727 | 0.001096465 | 0.199864 | 13.66625 | 3.051183 | -0.00633 |
| **HCP-YA** | rsfMRI | FP-Mot | 0.204286 | 0.001096465 | 0.203456 | 13.4878 | 2.999952 | 0.006806 |
| **HCP-YA** | rsfMRI | FP-SAL | 0.218031 | 0.001096465 | 0.222171 | 13.38744 | 2.990323 | 0.014196 |
| **HCP-YA** | rsfMRI | FP-SC | 0.197764 | 0.001096465 | 0.210715 | 13.68805 | 3.053066 | -0.00794 |
| **HCP-YA** | rsfMRI | FP-VAs | 0.153949 | 0.001096465 | 0.163068 | 13.87919 | 3.063746 | -0.02202 |
| **HCP-YA** | rsfMRI | FP-VII | 0.181 | 0.001096465 | 0.177096 | 13.37749 | 3.053363 | 0.014929 |
| **HCP-YA** | rsfMRI | FP-VI | 0.221626 | 0.001096465 | 0.236797 | 13.31661 | 3.001001 | 0.019411 |

| | | | | | | | | |
|---|---|---|---|---|---|---|---|---|
| **HCP-YA** | rsfMRI | MF-Mot | 0.210717 | 0.001096465 | 0.205843 | 13.25932 | 3.003691 | 0.02363 |
| **HCP-YA** | rsfMRI | MF-SAL | 0.180441 | 0.001096465 | 0.186131 | 13.5258 | 3.03034 | 0.004008 |
| **HCP-YA** | rsfMRI | MF-SC | 0.19223 | 0.001096465 | 0.198992 | 13.5406 | 3.030634 | 0.002918 |
| **HCP-YA** | rsfMRI | MF-VAs | 0.193176 | 0.001096465 | 0.200588 | 13.30633 | 3.00514 | 0.020168 |
| **HCP-YA** | rsfMRI | MF-VII | 0.206802 | 0.001096465 | 0.205594 | 13.29449 | 3.04315 | 0.02104 |
| **HCP-YA** | rsfMRI | MF-VI | 0.192311 | 0.001096465 | 0.200604 | 13.38884 | 3.01712 | 0.014093 |
| **HCP-YA** | rsfMRI | Mot-SAL | 0.166557 | 0.001096465 | 0.183831 | 13.50457 | 3.017906 | 0.005571 |
| **HCP-YA** | rsfMRI | Mot-SC | 0.20983 | 0.001096465 | 0.212086 | 13.28893 | 2.986501 | 0.02145 |
| **HCP-YA** | rsfMRI | Mot-VAs | 0.16186 | 0.001096465 | 0.17074 | 13.37142 | 3.013048 | 0.015375 |
| **HCP-YA** | rsfMRI | Mot-VII | 0.164124 | 0.001096465 | 0.177724 | 13.58775 | 3.05401 | -0.00055 |
| **HCP-YA** | rsfMRI | Mot-VI | 0.18723 | 0.001096465 | 0.197614 | 13.41763 | 3.022673 | 0.011972 |
| **HCP-YA** | rsfMRI | SAL-SC | 0.198178 | 0.001096465 | 0.208611 | 13.55488 | 3.001959 | 0.001866 |
| **HCP-YA** | rsfMRI | SAL-VAs | 0.15542 | 0.001096465 | 0.159338 | 13.61867 | 3.049729 | -0.00283 |
| **HCP-YA** | rsfMRI | SAL-VII | 0.176378 | 0.001096465 | 0.178328 | 13.50006 | 3.048799 | 0.005903 |
| **HCP-YA** | rsfMRI | SAL-VI | 0.196134 | 0.001096465 | 0.19639 | 13.49057 | 3.029998 | 0.006602 |
| **HCP-YA** | rsfMRI | SC-VAs | 0.164956 | 0.001096465 | 0.177483 | 13.62186 | 3.044834 | -0.00307 |
| **HCP-YA** | rsfMRI | SC-VII | 0.116206 | 0.014303878 | 0.127685 | 14.02656 | 3.113722 | -0.03287 |
| **HCP-YA** | rsfMRI | SC-VI | 0.157609 | 0.002140716 | 0.171243 | 13.65159 | 3.048075 | -0.00526 |
| **HCP-YA** | rsfMRI | VAs-VII | 0.121601 | 0.004181865 | 0.125699 | 13.69484 | 3.060146 | -0.00844 |
| **HCP-YA** | rsfMRI | VAs-VI | 0.164592 | 0.001096465 | 0.185834 | 13.43649 | 3.02507 | 0.010584 |
| **HCP-YA** | rsfMRI | VI-VII | 0.204747 | 0.001096465 | 0.213545 | 13.12881 | 3.016833 | 0.03324 |
| **HCP-A** | rsfMRI | CBL-DMN | 0.348384 | 0.000999001 | 0.343649 | 159.0687 | 10.1694 | 0.096519 |
| **HCP-A** | rsfMRI | CBL-FP | 0.448809 | 0.000999001 | 0.432844 | 142.8877 | 9.754378 | 0.188424 |
| **HCP-A** | rsfMRI | CBL-MF | 0.43484 | 0.000999001 | 0.423064 | 145.6011 | 9.525129 | 0.173013 |
| **HCP-A** | rsfMRI | CBL-Mot | 0.419232 | 0.000999001 | 0.399297 | 148.4176 | 9.898631 | 0.157016 |

| | | | | | | | | |
|---|---|---|---|---|---|---|---|---|
| **HCP-A** | rsfMRI | CBL-SAL | 0.470412 | 0.000999001 | 0.422493 | 137.9998 | 9.618492 | 0.216187 |
| **HCP-A** | rsfMRI | CBL-SC | 0.373482 | 0.000999001 | 0.385434 | 156.0501 | 9.9724 | 0.113664 |
| **HCP-A** | rsfMRI | CBL-VAs | 0.504506 | 0.000999001 | 0.476954 | 132.006 | 9.295328 | 0.25023 |
| **HCP-A** | rsfMRI | CBL-VII | 0.309275 | 0.000999001 | 0.296476 | 166.7838 | 10.51303 | 0.052699 |
| **HCP-A** | rsfMRI | CBL-VI | 0.350736 | 0.000999001 | 0.318665 | 160.755 | 10.3099 | 0.086942 |
| **HCP-A** | rsfMRI | DMN-FP | 0.319675 | 0.000999001 | 0.321328 | 165.2317 | 10.17098 | 0.061515 |
| **HCP-A** | rsfMRI | DMN-MF | 0.391969 | 0.000999001 | 0.393716 | 151.8452 | 9.879427 | 0.137547 |
| **HCP-A** | rsfMRI | DMN-Mot | 0.3945 | 0.000999001 | 0.393544 | 151.7352 | 9.836372 | 0.138172 |
| **HCP-A** | rsfMRI | DMN-SAL | 0.311419 | 0.000999001 | 0.308988 | 165.3702 | 10.26287 | 0.060728 |
| **HCP-A** | rsfMRI | DMN-SC | 0.347396 | 0.000999001 | 0.345271 | 159.533 | 10.12524 | 0.093882 |
| **HCP-A** | rsfMRI | DMN-VAs | 0.319392 | 0.000999001 | 0.294817 | 163.4027 | 10.218 | 0.071903 |
| **HCP-A** | rsfMRI | DMN-VII | 0.329602 | 0.000999001 | 0.336248 | 161.0125 | 10.29375 | 0.085479 |
| **HCP-A** | rsfMRI | DMN-VI | 0.332221 | 0.000999001 | 0.336833 | 160.6514 | 10.21842 | 0.08753 |
| **HCP-A** | rsfMRI | FP-MF | 0.362666 | 0.000999001 | 0.34107 | 157.7599 | 10.06184 | 0.103953 |
| **HCP-A** | rsfMRI | FP-Mot | 0.402178 | 0.000999001 | 0.380773 | 151.4675 | 10.00206 | 0.139693 |
| **HCP-A** | rsfMRI | FP-SAL | 0.379435 | 0.000999001 | 0.381563 | 156.4315 | 10.12855 | 0.111498 |
| **HCP-A** | rsfMRI | FP-SC | 0.336949 | 0.000999001 | 0.337269 | 163.0109 | 10.23108 | 0.074128 |
| **HCP-A** | rsfMRI | FP-VAs | 0.423087 | 0.000999001 | 0.413941 | 147.6517 | 9.818061 | 0.161366 |
| **HCP-A** | rsfMRI | FP-VII | 0.345986 | 0.000999001 | 0.351581 | 159.4635 | 10.17371 | 0.094277 |
| **HCP-A** | rsfMRI | FP-VI | 0.301306 | 0.000999001 | 0.271229 | 168.1105 | 10.37542 | 0.045164 |
| **HCP-A** | rsfMRI | MF-Mot | 0.37906 | 0.000999001 | 0.35512 | 154.3388 | 10.14511 | 0.123384 |
| **HCP-A** | rsfMRI | MF-SAL | 0.357699 | 0.000999001 | 0.353869 | 159.6412 | 10.03526 | 0.093267 |
| **HCP-A** | rsfMRI | MF-SC | 0.350785 | 0.000999001 | 0.347937 | 158.7121 | 10.28205 | 0.098545 |
| **HCP-A** | rsfMRI | MF-VAs | 0.426441 | 0.000999001 | 0.423367 | 146.085 | 9.793345 | 0.170265 |
| **HCP-A** | rsfMRI | MF-VII | 0.330688 | 0.000999001 | 0.323769 | 162.1297 | 10.19526 | 0.079134 |
| **HCP-A** | rsfMRI | MF-VI | 0.364953 | 0.000999001 | 0.373503 | 157.2126 | 10.09769 | 0.107061 |
| **HCP-A** | rsfMRI | Mot-SAL | 0.471626 | 0.000999001 | 0.44079 | 138.0399 | 9.494111 | 0.215959 |
| **HCP-A** | rsfMRI | Mot-SC | 0.388162 | 0.000999001 | 0.368343 | 153.9967 | 10.08323 | 0.125328 |
| **HCP-A** | rsfMRI | Mot-VAs | 0.396533 | 0.000999001 | 0.370903 | 151.1686 | 9.812671 | 0.14139 |
| **HCP-A** | rsfMRI | Mot-VII | 0.333984 | 0.000999001 | 0.289261 | 161.5757 | 10.24686 | 0.08228 |
| **HCP-A** | rsfMRI | Mot-VI | 0.395145 | 0.000999001 | 0.361159 | 152.5772 | 9.833461 | 0.13339 |
| **HCP-A** | rsfMRI | SAL-SC | 0.320467 | 0.000999001 | 0.318009 | 164.5897 | 10.29223 | 0.065161 |
| **HCP-A** | rsfMRI | SAL-VAs | 0.427101 | 0.000999001 | 0.396199 | 147.7308 | 9.856284 | 0.160916 |
| **HCP-A** | rsfMRI | SAL-VII | 0.288355 | 0.000999001 | 0.288005 | 167.91 | 10.60952 | 0.046302 |
| **HCP-A** | rsfMRI | SAL-VI | 0.358053 | 0.000999001 | 0.351012 | 158.1632 | 10.17495 | 0.101662 |
| **HCP-A** | rsfMRI | SC-VAs | 0.411136 | 0.000999001 | 0.397545 | 149.0686 | 9.760559 | 0.153318 |
| **HCP-A** | rsfMRI | SC-VII | 0.358337 | 0.000999001 | 0.360851 | 158.3969 | 9.930956 | 0.100335 |
| **HCP-A** | rsfMRI | SC-VI | 0.392731 | 0.000999001 | 0.38967 | 152.4253 | 10.01598 | 0.134253 |
| **HCP-A** | rsfMRI | VAs-VII | 0.33847 | 0.000999001 | 0.326204 | 159.7949 | 10.18728 | 0.092395 |
| **HCP-A** | rsfMRI | VAs-VI | 0.42113 | 0.000999001 | 0.403961 | 146.4266 | 9.811642 | 0.168324 |

| | | | | | | | | |
|---|---|---|---|---|---|---|---|---|
| **HCP-A** | rsfMRI | VI-VII | 0.351785 | 0.000999001 | 0.367438 | 156.8213 | 9.94504 | 0.109284 |
| **HCP-LS** | rsfMRI | CBL-DMN | 0.649689 | 0.000999001 | 0.633686 | 144.0239 | 8.579045 | 0.421926 |
| **HCP-LS** | rsfMRI | CBL-FP | 0.708365 | 0.000999001 | 0.734603 | 124.1548 | 7.852728 | 0.501676 |
| **HCP-LS** | rsfMRI | CBL-MF | 0.668249 | 0.000999001 | 0.664484 | 137.9305 | 8.297021 | 0.446383 |
| **HCP-LS** | rsfMRI | CBL-Mot | 0.68001 | 0.000999001 | 0.669142 | 133.9933 | 8.310623 | 0.462187 |
| **HCP-LS** | rsfMRI | CBL-SAL | 0.630496 | 0.000999001 | 0.641039 | 150.1882 | 8.783039 | 0.397184 |
| **HCP-LS** | rsfMRI | CBL-SC | 0.639319 | 0.000999001 | 0.690763 | 147.4178 | 8.462647 | 0.408304 |
| **HCP-LS** | rsfMRI | CBL-VAs | 0.649138 | 0.000999001 | 0.613738 | 144.2711 | 8.864446 | 0.420934 |
| **HCP-LS** | rsfMRI | CBL-VII | 0.68401 | 0.000999001 | 0.663187 | 132.6019 | 8.319114 | 0.467771 |
| **HCP-LS** | rsfMRI | CBL-VI | 0.668826 | 0.000999001 | 0.624753 | 137.7178 | 8.502992 | 0.447237 |
| **HCP-LS** | rsfMRI | DMN-FP | 0.74711 | 0.000999001 | 0.730676 | 110.1372 | 7.618757 | 0.557938 |
| **HCP-LS** | rsfMRI | DMN-MF | 0.67733 | 0.000999001 | 0.620279 | 134.8871 | 8.630067 | 0.458599 |
| **HCP-LS** | rsfMRI | DMN-Mot | 0.632193 | 0.000999001 | 0.60513 | 149.6395 | 8.9941 | 0.399386 |
| **HCP-LS** | rsfMRI | DMN-SAL | 0.62689 | 0.000999001 | 0.563033 | 151.2768 | 9.088729 | 0.392815 |
| **HCP-LS** | rsfMRI | DMN-SC | 0.680495 | 0.000999001 | 0.6584 | 133.8033 | 8.345747 | 0.462949 |
| **HCP-LS** | rsfMRI | DMN-VAs | 0.631678 | 0.000999001 | 0.610931 | 149.7701 | 9.025889 | 0.398862 |
| **HCP-LS** | rsfMRI | DMN-VII | 0.579072 | 0.000999001 | 0.582267 | 165.6632 | 9.466146 | 0.335072 |
| **HCP-LS** | rsfMRI | DMN-VI | 0.610731 | 0.000999001 | 0.507537 | 156.279 | 9.220784 | 0.372737 |
| **HCP-LS** | rsfMRI | FP-MF | 0.741228 | 0.000999001 | 0.714262 | 112.3072 | 7.549887 | 0.549229 |
| **HCP-LS** | rsfMRI | FP-Mot | 0.718279 | 0.000999001 | 0.698598 | 120.6595 | 7.911435 | 0.515705 |
| **HCP-LS** | rsfMRI | FP-SAL | 0.696268 | 0.000999001 | 0.667918 | 128.3842 | 8.248533 | 0.4847 |
| **HCP-LS** | rsfMRI | FP-SC | 0.731633 | 0.000999001 | 0.745243 | 115.8224 | 7.51979 | 0.53512 |
| **HCP-LS** | rsfMRI | FP-VAs | 0.697036 | 0.000999001 | 0.678089 | 128.1262 | 8.217066 | 0.485735 |
| **HCP-LS** | rsfMRI | FP-VII | 0.61993 | 0.000999001 | 0.613035 | 153.467 | 9.066355 | 0.384024 |

| | | | | | | | | |
|---|---|---|---|---|---|---|---|---|
| **HCP-LS** | rsfMRI | FP-VI | 0.690145 | 0.000999001 | 0.640178 | 130.5126 | 8.378493 | 0.476157 |
| **HCP-LS** | rsfMRI | MF-Mot | 0.651862 | 0.000999001 | 0.630759 | 143.3393 | 8.757682 | 0.424674 |
| **HCP-LS** | rsfMRI | MF-SAL | 0.638696 | 0.000999001 | 0.568927 | 147.5444 | 8.979694 | 0.407796 |
| **HCP-LS** | rsfMRI | MF-SC | 0.684709 | 0.000999001 | 0.673534 | 132.3728 | 8.163322 | 0.468691 |
| **HCP-LS** | rsfMRI | MF-VAs | 0.632679 | 0.000999001 | 0.617077 | 149.4828 | 8.904662 | 0.400015 |
| **HCP-LS** | rsfMRI | MF-VII | 0.576362 | 0.000999001 | 0.532428 | 166.5308 | 9.445739 | 0.331589 |
| **HCP-LS** | rsfMRI | MF-VI | 0.635264 | 0.000999001 | 0.484211 | 148.6227 | 9.158314 | 0.403468 |
| **HCP-LS** | rsfMRI | Mot-SAL | 0.648573 | 0.000999001 | 0.599037 | 144.3792 | 9.043391 | 0.4205 |
| **HCP-LS** | rsfMRI | Mot-SC | 0.67426 | 0.000999001 | 0.605914 | 135.9279 | 8.576934 | 0.454421 |
| **HCP-LS** | rsfMRI | Mot-VAs | 0.640252 | 0.000999001 | 0.501062 | 147.0579 | 9.101692 | 0.409749 |
| **HCP-LS** | rsfMRI | Mot-VII | 0.588077 | 0.000999001 | 0.397384 | 163.0271 | 9.691765 | 0.345652 |
| **HCP-LS** | rsfMRI | Mot-VI | 0.580305 | 0.000999001 | 0.397446 | 165.3425 | 9.787065 | 0.336359 |
| **HCP-LS** | rsfMRI | SAL-SC | 0.642831 | 0.000999001 | 0.638988 | 146.2298 | 8.663606 | 0.413072 |
| **HCP-LS** | rsfMRI | SAL-VAs | 0.58674 | 0.000999001 | 0.571694 | 163.4802 | 9.453854 | 0.343834 |
| **HCP-LS** | rsfMRI | SAL-VII | 0.626903 | 0.000999001 | 0.564122 | 151.2727 | 9.147383 | 0.392832 |
| **HCP-LS** | rsfMRI | SAL-VI | 0.632284 | 0.000999001 | 0.478188 | 149.5571 | 9.347311 | 0.399717 |
| **HCP-LS** | rsfMRI | SC-VAs | 0.625303 | 0.000999001 | 0.556373 | 151.7989 | 8.987075 | 0.390719 |
| **HCP-LS** | rsfMRI | SC-VII | 0.634513 | 0.000999001 | 0.580646 | 148.9171 | 8.865287 | 0.402286 |
| **HCP-LS** | rsfMRI | SC-VI | 0.646189 | 0.000999001 | 0.510402 | 145.1497 | 9.091986 | 0.417408 |
| **HCP-LS** | rsfMRI | VAs-VII | 0.590791 | 0.000999001 | 0.544098 | 162.3213 | 9.410326 | 0.348485 |
| **HCP-LS** | rsfMRI | VAs-VI | 0.574857 | 0.000999001 | 0.509092 | 166.9496 | 9.435548 | 0.329909 |
| **HCP-LS** | rsfMRI | VI-VII | 0.560398 | 0.000999001 | 0.553794 | 171.072 | 9.514761 | 0.313362 |
| | | | | **Within-Network** | | | | |
| **Cohort** | **Modality** | **Network** | **Pearson r** | **pFDR** | **Spearman rho** | **MSE** | **MAE** | **q2** |

| | | | | | | | | |
|---|---|---|---|---|---|---|---|---|
| **HCP-D** | dMRI QA End | CBL | 0.25851 | 0.001297391 | 0.266478 | 13.51904 | 3.037916 | 0.047474 |
| **HCP-D** | dMRI QA End | DMN | 0.288015 | 0.001297391 | 0.295794 | 13.10072 | 2.942279 | 0.076949 |
| **HCP-D** | dMRI QA End | FP | 0.277555 | 0.001297391 | 0.281073 | 13.28045 | 2.99321 | 0.064285 |
| **HCP-D** | dMRI QA End | MF | 0.377628 | 0.001297391 | 0.380691 | 12.22788 | 2.910166 | 0.138447 |
| **HCP-D** | dMRI QA End | Mot | 0.28109 | 0.001297391 | 0.278267 | 13.12255 | 2.992034 | 0.07541 |
| **HCP-D** | dMRI QA End | SAL | 0.341398 | 0.001297391 | 0.360372 | 12.63617 | 2.901088 | 0.10968 |
| **HCP-D** | dMRI QA End | SC | 0.390864 | 0.001297391 | 0.401782 | 12.15894 | 2.836964 | 0.143304 |
| **HCP-D** | dMRI QA End | VAs | 0.224386 | 0.002360965 | 0.233064 | 13.52777 | 3.072252 | 0.04686 |
| **HCP-D** | dMRI QA End | VII | NaN | NA | NaN | NaN | NaN | NaN |
| **HCP-D** | dMRI QA End | VI | 0.141869 | 0.007042254 | 0.143395 | 14.1027 | 3.117579 | 0.006351 |
| **HCP-YA** | dMRI QA End | CBL | 0.12874 | 0.04995005 | 0.129347 | 13.67999 | 3.08468 | -0.00735 |
| **HCP-YA** | dMRI QA End | DMN | 0.010222 | 0.402097902 | 0.017886 | 13.78471 | 3.110105 | -0.01506 |
| **HCP-YA** | dMRI QA End | FP | 0.06617 | 0.176490176 | 0.053208 | 14.08488 | 3.129753 | -0.03716 |
| **HCP-YA** | dMRI QA End | MF | 0.072514 | 0.127372627 | 0.072157 | 13.76122 | 3.118967 | -0.01333 |
| **HCP-YA** | dMRI QA End | Mot | 0.093425 | 0.11988012 | 0.108335 | 14.03607 | 3.125689 | -0.03357 |
| **HCP-YA** | dMRI QA End | SAL | 0.071874 | 0.173826174 | 0.066546 | 14.03448 | 3.148078 | -0.03345 |
| **HCP-YA** | dMRI QA End | SC | 0.086657 | 0.11988012 | 0.093305 | 13.72189 | 3.075537 | -0.01043 |
| **HCP-YA** | dMRI QA End | VAs | -0.07659 | 0.700763359 | -0.0678 | 14.0131 | 3.137388 | -0.02683 |
| **HCP-YA** | dMRI QA End | VII | -0.10688 | 0.700763359 | -0.12258 | 15.4885 | 3.326253 | -0.0253 |
| **HCP-YA** | dMRI QA End | VI | 0.009958 | 0.402097902 | 0.01948 | 14.05105 | 3.112165 | -0.03467 |
| **HCP-A** | dMRI QA End | CBL | 0.520139 | 0.0017507 | 0.458769 | 128.7838 | 9.243365 | 0.268532 |
| **HCP-A** | dMRI QA End | DMN | 0.151343 | 0.022982635 | 0.144291 | 177.2138 | 10.93755 | -0.00654 |
| **HCP-A** | dMRI QA End | FP | 0.467891 | 0.0017507 | 0.439014 | 138.2433 | 9.587778 | 0.214804 |

| | | | | | | | | |
|---|---|---|---|---|---|---|---|---|
| **HCP-A** | dMRI QA End | MF | 0.131645 | 0.038005813 | 0.172682 | 179.7364 | 11.00758 | -0.02087 |
| **HCP-A** | dMRI QA End | Mot | 0.358746 | 0.0017507 | 0.328595 | 153.9305 | 10.17417 | 0.125703 |
| **HCP-A** | dMRI QA End | SAL | 0.465505 | 0.0017507 | 0.445399 | 138.2045 | 9.479928 | 0.215024 |
| **HCP-A** | dMRI QA End | SC | 0.45808 | 0.0017507 | 0.455031 | 139.8697 | 9.726407 | 0.205566 |
| **HCP-A** | dMRI QA End | VAs | 0.448672 | 0.0017507 | 0.399996 | 140.7724 | 9.636457 | 0.200439 |
| **HCP-A** | dMRI QA End | VII | 0.124076 | 0.0675 | 0.102237 | 173.9467 | 10.98973 | 0.012015 |
| **HCP-A** | dMRI QA End | VI | 0.180227 | 0.022982635 | 0.149579 | 172.9109 | 10.97817 | 0.017898 |
| **HCP-LS** | dMRI QA End | CBL | 0.549456 | 0.001172058 | 0.36595 | 173.951 | 10.10838 | 0.301807 |
| **HCP-LS** | dMRI QA End | DMN | 0.355566 | 0.001172058 | 0.106826 | 217.6631 | 11.06387 | 0.126358 |
| **HCP-LS** | dMRI QA End | FP | 0.513543 | 0.001172058 | 0.324152 | 183.4662 | 10.39427 | 0.263615 |
| **HCP-LS** | dMRI QA End | MF | 0.381545 | 0.001172058 | 0.150236 | 212.9116 | 11.12656 | 0.145429 |
| **HCP-LS** | dMRI QA End | Mot | 0.434937 | 0.001172058 | 0.252474 | 202.0888 | 10.73091 | 0.188869 |
| **HCP-LS** | dMRI QA End | SAL | 0.553174 | 0.001172058 | 0.370382 | 172.9301 | 10.10008 | 0.305905 |
| **HCP-LS** | dMRI QA End | SC | 0.538668 | 0.001172058 | 0.36109 | 176.8577 | 10.2601 | 0.29014 |
| **HCP-LS** | dMRI QA End | VAs | 0.371095 | 0.001172058 | 0.360214 | 215.116 | 10.71246 | 0.136581 |
| **HCP-LS** | dMRI QA End | VII | 0.088934 | 0.021220159 | -0.05004 | 247.2473 | 11.33261 | 0.007615 |
| **HCP-LS** | dMRI QA End | VI | 0.386413 | 0.001172058 | 0.22448 | 212.0412 | 10.83479 | 0.148923 |
| **HCP-D** | rsfMRI | CBL | 0.641498 | 0.001012146 | 0.658845 | 8.352764 | 2.303679 | 0.41148 |
| **HCP-D** | rsfMRI | DMN | 0.458213 | 0.001012146 | 0.468135 | 11.26163 | 2.740295 | 0.206527 |
| **HCP-D** | rsfMRI | FP | 0.683651 | 0.001012146 | 0.680712 | 7.559528 | 2.206001 | 0.46737 |
| **HCP-D** | rsfMRI | MF | 0.582809 | 0.001012146 | 0.583575 | 9.385214 | 2.420036 | 0.338736 |
| **HCP-D** | rsfMRI | Mot | 0.624288 | 0.001012146 | 0.628713 | 8.6689 | 2.425944 | 0.389206 |
| **HCP-D** | rsfMRI | SAL | 0.605918 | 0.001012146 | 0.648901 | 8.989383 | 2.393535 | 0.366626 |
| **HCP-D** | rsfMRI | SC | 0.670349 | 0.001012146 | 0.679622 | 7.815999 | 2.22892 | 0.4493 |
| **HCP-D** | rsfMRI | VAs | 0.573869 | 0.001012146 | 0.579268 | 9.527446 | 2.481873 | 0.328715 |
| **HCP-D** | rsfMRI | VII | 0.32078 | 0.001012146 | 0.318295 | 12.8457 | 2.937565 | 0.094917 |
| **HCP-D** | rsfMRI | VI | 0.514349 | 0.001012146 | 0.521677 | 10.48343 | 2.655342 | 0.261358 |
| **HCP-YA** | rsfMRI | CBL | 0.16912 | 0.001427144 | 0.176134 | 13.31992 | 3.012979 | 0.019168 |

| | | | | | | | | |
|---|---|---|---|---|---|---|---|---|
| **HCP-YA** | rsfMRI | DMN | 0.144045 | 0.001427144 | 0.14275 | 13.52533 | 3.073616 | 0.004042 |
| **HCP-YA** | rsfMRI | FP | 0.197103 | 0.001427144 | 0.191739 | 13.54532 | 3.05807 | 0.00257 |
| **HCP-YA** | rsfMRI | MF | 0.211125 | 0.001427144 | 0.216222 | 13.25345 | 3.014611 | 0.024062 |
| **HCP-YA** | rsfMRI | Mot | 0.179108 | 0.001427144 | 0.187577 | 13.34396 | 3.018267 | 0.017398 |
| **HCP-YA** | rsfMRI | SAL | 0.161798 | 0.001427144 | 0.166051 | 13.55377 | 3.060282 | 0.001948 |
| **HCP-YA** | rsfMRI | SC | 0.165772 | 0.002497502 | 0.171516 | 13.5887 | 3.036297 | -0.00062 |
| **HCP-YA** | rsfMRI | VAs | 0.138381 | 0.003330003 | 0.156801 | 13.43124 | 3.033582 | 0.010971 |
| **HCP-YA** | rsfMRI | VII | 0.088535 | 0.043478261 | 0.097315 | 13.60059 | 3.086018 | -0.0015 |
| **HCP-YA** | rsfMRI | VI | 0.194053 | 0.001427144 | 0.201921 | 13.22201 | 3.026366 | 0.026377 |
| **HCP-A** | rsfMRI | CBL | 0.452717 | 0.001665002 | 0.426208 | 141.182 | 9.663032 | 0.198113 |
| **HCP-A** | rsfMRI | DMN | 0.315735 | 0.001665002 | 0.306156 | 161.6343 | 10.22272 | 0.081947 |
| **HCP-A** | rsfMRI | FP | 0.30662 | 0.001665002 | 0.320318 | 167.1828 | 10.38963 | 0.050433 |
| **HCP-A** | rsfMRI | MF | 0.3525 | 0.001665002 | 0.34378 | 158.2984 | 10.20798 | 0.100894 |
| **HCP-A** | rsfMRI | Mot | 0.32819 | 0.001665002 | 0.296745 | 163.0494 | 10.21587 | 0.07391 |
| **HCP-A** | rsfMRI | SAL | 0.224596 | 0.004281433 | 0.195968 | 178.9326 | 10.74862 | -0.0163 |
| **HCP-A** | rsfMRI | SC | 0.154398 | 0.040959041 | 0.149611 | 187.6954 | 11.17347 | -0.06607 |
| **HCP-A** | rsfMRI | VAs | 0.391965 | 0.001665002 | 0.381523 | 152.3002 | 9.897172 | 0.134963 |
| **HCP-A** | rsfMRI | VII | 0.155332 | 0.023474178 | 0.17808 | 177.9427 | 10.81036 | -0.01068 |
| **HCP-A** | rsfMRI | VI | 0.159305 | 0.023474178 | 0.162871 | 184.392 | 11.04493 | -0.04731 |
| **HCP-LS** | rsfMRI | CBL | 0.634566 | 0.001010101 | 0.691282 | 148.9586 | 8.633962 | 0.40212 |
| **HCP-LS** | rsfMRI | DMN | 0.57688 | 0.001010101 | 0.530279 | 166.3086 | 9.491483 | 0.332481 |
| **HCP-LS** | rsfMRI | FP | 0.707596 | 0.001010101 | 0.705716 | 124.4259 | 7.970628 | 0.500587 |
| **HCP-LS** | rsfMRI | MF | 0.686038 | 0.001010101 | 0.623388 | 131.8994 | 8.426534 | 0.470591 |
| **HCP-LS** | rsfMRI | Mot | 0.657743 | 0.001010101 | 0.434792 | 141.404 | 9.107414 | 0.432442 |
| **HCP-LS** | rsfMRI | SAL | 0.536595 | 0.001010101 | 0.514029 | 177.5008 | 9.8515 | 0.287559 |
| **HCP-LS** | rsfMRI | SC | 0.590949 | 0.001010101 | 0.594713 | 162.3728 | 9.197683 | 0.348279 |
| **HCP-LS** | rsfMRI | VAs | 0.506633 | 0.001010101 | 0.466388 | 185.7679 | 9.973344 | 0.254377 |
| **HCP-LS** | rsfMRI | VII | 0.39586 | 0.001010101 | 0.372392 | 210.3214 | 10.53418 | 0.155826 |

| | | | | | | | | |
|---|---|---|---|---|---|---|---|---|
| **HCP-LS** | rsfMRI | VI | 0.561599 | 0.001010101 | 0.436786 | 170.6314 | 9.827795 | 0.315131 |

*Table S5.CPM performance metrics from the median performing diffusion MRI (dMRI) QA End and resting-state fMRI (rsfMRI) between-network and within-network models for each age cohort (HCP-D, HCP-YA, HCP-A, and HCP-LS). Networks based on the 10 Shen networks. Model performance was evaluated with Pearson's correlation (r), Spearman's rank correlation (ρ), mean absolute error (MAE), mean squared error (MSE), and the predictive coefficient of determination ($q^2$). Network abbreviations: CBL = cerebellar, DMN = default mode network, FP = frontoparietal, MF = medial frontal, Mot = somatomotor, SAL = salience, SC = subcortical, VAs = visual association, VI = visual I, VII = visual II.*

**Sex-stratified CPM age prediction and cross-sex model generalizability**

| Modality | Cohort | Sex | Pearson r | Null CPM FDR-corrected p | Spearman rho | MAE | MSE | q² |
|---|---|---|---|---|---|---|---|---|
| **dMRI QA End** | HCP-D | Males | 0.534466 | 1.00E-04 | 0.541083 | 2.421953 | 9.331308 | 0.282661 |
| | | Females | 0.627691 | 1.00E-04 | 0.644451 | 2.460358 | 9.385997 | 0.381782 |
| | HCP-YA | Males | 0.150632 | 0.019948 | 0.169417 | 3.080099 | 14.28818 | -0.09711 |
| | | Females | 0.165892 | 0.007199 | 0.161895 | 3.027183 | 13.70702 | -0.06126 |
| | HCP-A | Males | 0.722296 | 1.00E-04 | 0.707868 | 7.800144 | 92.10506 | 0.510389 |
| | | Females | 0.658386 | 1.00E-04 | 0.605994 | 7.579138 | 94.60989 | 0.429874 |
| | HCP-LS | Males | 0.754154 | 1.00E-04 | 0.44059 | 7.976077 | 104.2784 | 0.567791 |
| | | Females | 0.691975 | 1.00E-04 | 0.516368 | 9.101914 | 132.9666 | 0.478679 |
| **rsfMRI** | HCP-D | Males | 0.764677 | 1.00E-04 | 0.778508 | 1.799371 | 5.439398 | 0.58185 |
| | | Females | 0.721817 | 1.00E-04 | 0.737684 | 2.175247 | 7.277752 | 0.520644 |
| | HCP-YA | Males | 0.255165 | 0.0003 | 0.265104 | 2.9758 | 13.18602 | -0.01248 |
| | | Females | 0.244104 | 1.00E-04 | 0.25203 | 2.888076 | 12.67014 | 0.019017 |
| | HCP-A | Males | 0.501133 | 1.00E-04 | 0.47796 | 9.710795 | 141.0995 | 0.249944 |
| | | Females | 0.446157 | 1.00E-04 | 0.401498 | 9.1216 | 134.375 | 0.190247 |
| | HCP-LS | Males | 0.762982 | 1.00E-04 | 0.689356 | 7.257383 | 101.2782 | 0.580226 |
| | | Females | 0.757785 | 1.00E-04 | 0.751199 | 7.614267 | 108.9075 | 0.573008 |

*Table S6. CPM performance metrics from the median performing dMRI QA End and rsfMRI male-only and female-only models for each age cohort (HCP-D, HCP-YA, HCP-A, and HCP-LS). Model performance was evaluated with Pearson's correlation (r), Spearman's rank correlation (ρ), mean absolute error (MAE), mean squared error (MSE), and the predictive coefficient of determination ($q^2$).*

#### Comparison of male and female CPM performance metrics

| Cohort | Modality | Male r | Female r | Male N | Female N | Fisher Z | P | FDR |
|---|---|---|---|---|---|---|---|---|
| **HCP_A** | dMRI QA End | 0.722296 | 0.658386 | 119 | 171 | 1.0145 | 0.310344 | 0.720881 |
| **HCP_A** | rsfMRI | 0.501133 | 0.446157 | 119 | 171 | 0.587531 | 0.556847 | 0.751124 |
| **HCP_D** | dMRI QA End | 0.534466 | 0.627691 | 210 | 248 | -1.49588 | 0.134684 | 0.538736 |
| **HCP_D** | rsfMRI | 0.764677 | 0.721817 | 210 | 248 | 1.016381 | 0.309448 | 0.720881 |
| **HCP_LS** | dMRI QA End | 0.754154 | 0.691975 | 684 | 787 | 2.496679 | 0.012536 | 0.150435 |
| **HCP_LS** | rsfMRI | 0.762982 | 0.757785 | 684 | 787 | 0.235221 | 0.814037 | 0.874528 |
| **HCP_YA** | dMRI QA End | 0.150632 | 0.165892 | 355 | 368 | -0.20953 | 0.834033 | 0.874528 |
| **HCP_YA** | rsfMRI | 0.255165 | 0.244104 | 355 | 368 | 0.157909 | 0.874528 | 0.874528 |

*Table S7. Comparison of Pearson r correlations from the median performing Male-only and female-only CPM models using Fisher r-to-z transformation.*

| Cohort | Modality | Train Sex | Test Sex | Pearson r | p-value | FDR-corrected p | Spearman rho | MSE | MAE | $q^2$ |
|---|---|---|---|---|---|---|---|---|---|---|
| **HCP-D** | dMRI QA End | F | M | 0.731634 | 1.83E-36 | 4.89E-36 | 0.737013 | 10.87421 | 2.650462 | 0.164052 |
| **HCP-D** | dMRI QA End | M | F | 0.403659 | 3.89E-11 | 5.84E-11 | 0.405752 | 120.3493 | 10.1214 | -6.92692 |
| **HCP-D** | rsfMRI | F | M | 0.704177 | 9.07E-33 | 2.18E-32 | 0.714814 | 13.29395 | 2.908887 | -0.02196 |
| **HCP-D** | rsfMRI | M | F | 0.74135 | 1.69E-44 | 5.78E-44 | 0.762822 | 12.2562 | 2.666601 | 0.192734 |
| **HCP-YA** | dMRI QA End | F | M | 0.191716 | 0.0002799 | 0.00031988 | 0.192459 | 56.33887 | 6.390409 | -3.32596 |
| **HCP-YA** | dMRI QA End | M | F | 0.154329 | 0.0029947 | 0.00312489 | 0.139587 | 113.2933 | 9.617725 | -7.77172 |
| **HCP-YA** | rsfMRI | F | M | 0.094027 | 0.0768466 | 0.07684663 | 0.098974 | 21.26099 | 3.649104 | -0.63252 |
| **HCP-YA** | rsfMRI | M | F | 0.160889 | 0.001961 | 0.00213928 | 0.178538 | 128.7757 | 10.32547 | -8.97044 |
| **HCP-A** | dMRI QA End | F | M | 0.769592 | 1.53E-24 | 2.83E-24 | 0.757712 | 254.0154 | 13.48417 | -0.35029 |

| | | | | | | | | | | |
|---|---|---|---|---|---|---|---|---|---|---|
| **HCP-A** | dMRI QA End | M | F | 0.596608 | 7.27E-18 | 1.16E-17 | 0.541055 | 279.7627 | 14.00529 | -0.68587 |
| **HCP-A** | rsfMRI | F | M | 0.540724 | 2.18E-10 | 3.08E-10 | 0.552018 | 202.7965 | 11.85807 | -0.07802 |
| **HCP-A** | rsfMRI | M | F | 0.406674 | 3.40E-08 | 4.54E-08 | 0.394094 | 200.9307 | 11.45749 | -0.21082 |
| **HCP-LS** | dMRI QA End | F | M | 0.734636 | 5.23E-117 | 2.09E-116 | 0.41781 | 1474.789 | 35.53221 | -5.11265 |
| **HCP-LS** | dMRI QA End | M | F | 0.709105 | 3.03E-121 | 1.45E-120 | 0.53956 | 1275.641 | 32.44 | -4.00139 |
| **HCP-LS** | rsfMRI | F | M | 0.764291 | 4.59E-132 | 3.67E-131 | 0.736569 | 233.4071 | 11.41796 | 0.032582 |
| **HCP-LS** | rsfMRI | M | F | 0.713537 | 2.01E-123 | 1.21E-122 | 0.691266 | 224.0612 | 11.76612 | 0.121526 |

*Table S8. Cross-sex CPM model generalizability using the final CPM models.*

**Multimodal CPM**

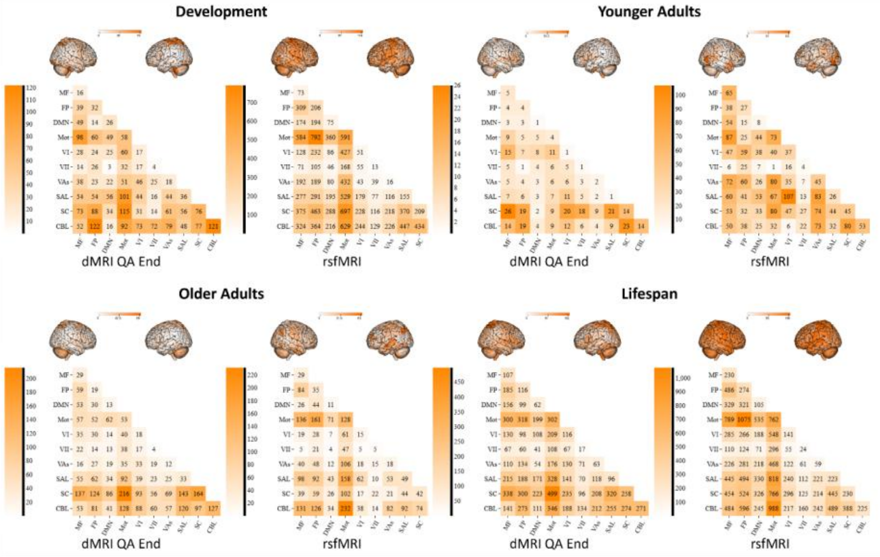

*Figure S2. Modality-specific consensus connectivity patterns contributing to multimodal CPM-based age prediction. Visualization of representative consensus networks extracted from the median-performing multimodal CPM model, which integrated dMRI QA End and rsfMRI features for age prediction. Modality-specific predictive networks were reconstructed separately for the dMRI QA End and rsfMRI*

*components of the multimodal model across developmental (HCP-D; top left), young adult (HCP-YA; top right), older adult (HCP-A; bottom left), and lifespan (HCP-LS; bottom right) cohorts. Brain renderings above each matrix illustrate the anatomical distribution of modality-specific age-predictive edges mapped onto the Shen 268-node atlas. Corresponding absolute sum matrices display the magnitude of predictive edge contributions between atlas-defined regions, with higher values indicating regions connected by a greater number of predictive edges regardless of directionality. These modality-specific consensus networks represent the structural and functional connectivity features contributing to multimodal CPM-based age prediction across the lifespan.*

| Multimodal CPM | | |
|---|---|---|
| **Age Cohort** | **dMRI QA End Total Edges** | **rsfMRI Total Edges** |
| **HCP-Development** | 2,636 | 13,743 |
| **HCP-Young Adults** | 717 | 2,373 |
| **HCP-Aging** | 3,152 | 3004 |
| **HCP-Lifespan** | 9,612 | 18,745 |
| **CPM, Connectome-based Predictive Modeling; HCP, Human Connectome Project; dMRI, diffusion MRI; QA, quantitative anisotropy; rsfMRI, resting-state functional MRI** | | |

*Table S9. Total edge density from dMRI QA End and rsfMRI components from the median performing multimodal consensus predictive networks.*

#### Multimodal vs unimodal CPM performance

| Cohort | Comparison | Metric | Iterations Favoring Multimodal | Iterations Favoring Unimodal | % Iterations Multimodal Better | p-value | FDR-corrected p |
|---|---|---|---|---|---|---|---|
| **HCP_A** | multimodal vs dMRI QA End | mae | 0 | 100 | 0 | 1 | 1 |
| **HCP_A** | multimodal vs dMRI QA End | mse | 0 | 100 | 0 | 1 | 1 |
| **HCP_A** | multimodal vs dMRI QA End | pearson_r | 0 | 100 | 0 | 1 | 1 |
| **HCP_D** | multimodal vs dMRI QA End | mae | 100 | 0 | 100 | 7.89E-31 | 1.05E-30 |
| **HCP_D** | multimodal vs dMRI QA End | mse | 100 | 0 | 100 | 7.89E-31 | 1.05E-30 |
| **HCP_D** | multimodal vs dMRI QA End | pearson_r | 100 | 0 | 100 | 7.89E-31 | 1.05E-30 |
| **HCP_LS** | multimodal vs dMRI QA End | mae | 100 | 0 | 100 | 7.89E-31 | 1.05E-30 |
| **HCP_LS** | multimodal vs dMRI QA End | mse | 100 | 0 | 100 | 7.89E-31 | 1.05E-30 |
| **HCP_LS** | multimodal vs dMRI QA End | pearson_r | 100 | 0 | 100 | 7.89E-31 | 1.05E-30 |

| | | | | | | | |
|---|---|---|---|---|---|---|---|
| **HCP_YA** | multimodal vs dMRI QA End | mae | 100 | 0 | 100 | 7.89E-31 | 1.05E-30 |
| **HCP_YA** | multimodal vs dMRI QA End | mse | 100 | 0 | 100 | 7.89E-31 | 1.05E-30 |
| **HCP_YA** | multimodal vs dMRI QA End | pearson_r | 96 | 4 | 96 | 3.22E-24 | 3.97E-24 |
| **HCP_A** | multimodal vs rsfMRI | mae | 100 | 0 | 100 | 7.89E-31 | 1.05E-30 |
| **HCP_A** | multimodal vs rsfMRI | mse | 100 | 0 | 100 | 7.89E-31 | 1.05E-30 |
| **HCP_A** | multimodal vs rsfMRI | pearson_r | 100 | 0 | 100 | 7.89E-31 | 1.05E-30 |
| **HCP_D** | multimodal vs rsfMRI | mae | 0 | 100 | 0 | 1 | 1 |
| **HCP_D** | multimodal vs rsfMRI | mse | 25 | 75 | 25 | 1 | 1 |
| **HCP_D** | multimodal vs rsfMRI | pearson_r | 33 | 67 | 33 | 0.999796 | 1 |
| **HCP_LS** | multimodal vs rsfMRI | mae | 100 | 0 | 100 | 7.89E-31 | 1.05E-30 |
| **HCP_LS** | multimodal vs rsfMRI | mse | 100 | 0 | 100 | 7.89E-31 | 1.05E-30 |
| **HCP_LS** | multimodal vs rsfMRI | pearson_r | 100 | 0 | 100 | 7.89E-31 | 1.05E-30 |
| **HCP_YA** | multimodal vs rsfMRI | mae | 99 | 1 | 99 | 7.97E-29 | 1.03E-28 |
| **HCP_YA** | multimodal vs rsfMRI | mse | 96 | 4 | 96 | 3.22E-24 | 3.97E-24 |
| **HCP_YA** | multimodal vs rsfMRI | pearson_r | 100 | 0 | 100 | 7.89E-31 | 1.05E-30 |

*Table S10. Paired iteration-wise comparisons of multimodal and unimodal CPM performance across cohorts.*

**Multimodal-unimodal CPM similarity**

| **Edge-level** |
|---|

| Comparison | Cohort | Type | Edge Correlation r | Edge Correlation p | Edge Correlation p-FDR | Hypergeometric p | Hypergeometric p-FDR | Jaccard |
|---|---|---|---|---|---|---|---|---|
| **DTI_MultiQAEND** | HCP-D | Positive | 0.862992169 | 1.00E-04 | 1.00E-04 | 3.22E-229 | 3.22E-229 | 0.761726 |
| **DTI_MultiQAEND** | HCP-D | Negative | 0.892386351 | 1.00E-04 | 1.00E-04 | 3.22E-229 | 3.22E-229 | 0.816395 |
| **DTI_MultiQAEND** | HCP-D | Absolute | 0.884305198 | 1.00E-04 | 1.00E-04 | 3.22E-229 | 3.22E-229 | 0.80643 |
| **DTI_MultiQAEND** | HCP-YA | Positive | 0.7481287 | 1.00E-04 | 1.00E-04 | 3.22E-229 | 3.22E-229 | 0.598765 |
| **DTI_MultiQAEND** | HCP-YA | Negative | 0.799818129 | 1.00E-04 | 1.00E-04 | 3.22E-229 | 3.22E-229 | 0.670851 |
| **DTI_MultiQAEND** | HCP-YA | Absolute | 0.789220754 | 1.00E-04 | 1.00E-04 | 3.22E-229 | 3.22E-229 | 0.657565 |
| **DTI_MultiQAEND** | HCP-A | Positive | 0.899368981 | 1.00E-04 | 1.00E-04 | 3.22E-229 | 3.22E-229 | 0.824169 |
| **DTI_MultiQAEND** | HCP-A | Negative | 0.908005024 | 1.00E-04 | 1.00E-04 | 3.22E-229 | 3.22E-229 | 0.838583 |
| **DTI_MultiQAEND** | HCP-A | Absolute | 0.899226648 | 1.00E-04 | 1.00E-04 | 3.22E-229 | 3.22E-229 | 0.831634 |
| **DTI_MultiQAEND** | HCP-LS | Positive | 0.957028075 | 1.00E-04 | 1.00E-04 | 3.22E-229 | 3.22E-229 | 0.930157 |
| **DTI_MultiQAEND** | HCP-LS | Negative | 0.945055923 | 1.00E-04 | 1.00E-04 | 3.22E-229 | 3.22E-229 | 0.906763 |
| **DTI_MultiQAEND** | HCP-LS | Absolute | 0.943405427 | 1.00E-04 | 1.00E-04 | 3.22E-229 | 3.22E-229 | 0.920468 |
| **REST_MultiQAEND** | HCP-D | Positive | 0.829275683 | 1.00E-04 | 1.00E-04 | 3.22E-229 | 3.22E-229 | 0.749563 |
| **REST_MultiQAEND** | HCP-D | Negative | 0.827072509 | 1.00E-04 | 1.00E-04 | 3.22E-229 | 3.22E-229 | 0.746385 |
| **REST_MultiQAEND** | HCP-D | Absolute | 0.781772705 | 1.00E-04 | 1.00E-04 | 3.22E-229 | 3.22E-229 | 0.748 |
| **REST_MultiQAEND** | HCP-YA | Positive | 0.810472383 | 1.00E-04 | 1.00E-04 | 3.22E-229 | 3.22E-229 | 0.689911 |

| | | | | | | | | |
|---|---|---|---|---|---|---|---|---|
| **REST_MultiQAEND** | HCP-YA | Negative | 0.830997128 | 1.00E-04 | 1.00E-04 | 3.22E-229 | 3.22E-229 | 0.718593 |
| **REST_MultiQAEND** | HCP-YA | Absolute | 0.814636408 | 1.00E-04 | 1.00E-04 | 3.22E-229 | 3.22E-229 | 0.704487 |
| **REST_MultiQAEND** | HCP-A | Positive | 0.820604746 | 1.00E-04 | 1.00E-04 | 3.22E-229 | 3.22E-229 | 0.701481 |
| **REST_MultiQAEND** | HCP-A | Negative | 0.825918144 | 1.00E-04 | 1.00E-04 | 3.22E-229 | 3.22E-229 | 0.712826 |
| **REST_MultiQAEND** | HCP-A | Absolute | 0.816540282 | 1.00E-04 | 1.00E-04 | 3.22E-229 | 3.22E-229 | 0.708195 |
| **REST_MultiQAEND** | HCP-LS | Positive | 0.921001084 | 1.00E-04 | 1.00E-04 | 3.22E-229 | 3.22E-229 | 0.889296 |
| **REST_MultiQAEND** | HCP-LS | Negative | 0.91655628 | 1.00E-04 | 1.00E-04 | 3.22E-229 | 3.22E-229 | 0.883008 |
| **REST_MultiQAEND** | HCP-LS | Absolute | 0.875471518 | 1.00E-04 | 1.00E-04 | 3.22E-229 | 3.22E-229 | 0.886157 |
| **Node-level** | | | | | | | | |
| **Comparison** | **Cohort** | **Type** | **Node Correlation r** | **Node Correlation p** | **Node Correlation p-FDR** | | | |
| **DTI_MultiQAEND** | HCP-D | Positive | 0.969045174 | 1.00E-04 | 1.00E-04 | | | |
| **DTI_MultiQAEND** | HCP-D | Negative | 0.990525503 | 1.00E-04 | 1.00E-04 | | | |
| **DTI_MultiQAEND** | HCP-D | Absolute | 0.987410132 | 1.00E-04 | 1.00E-04 | | | |
| **DTI_MultiQAEND** | HCP-YA | Positive | 0.923106693 | 1.00E-04 | 1.00E-04 | | | |
| **DTI_MultiQAEND** | HCP-YA | Negative | 0.956704259 | 1.00E-04 | 1.00E-04 | | | |
| **DTI_MultiQAEND** | HCP-YA | Absolute | 0.946590201 | 1.00E-04 | 1.00E-04 | | | |
| **DTI_MultiQAEND** | HCP-A | Positive | 0.992575381 | 1.00E-04 | 1.00E-04 | | | |
| **DTI_MultiQAEND** | HCP-A | Negative | 0.991788786 | 1.00E-04 | 1.00E-04 | | | |

| | | | | | |
|---|---|---|---|---|---|
| **DTI_MultiQAEND** | HCP-A | Absolute | 0.99167028 | 1.00E-04 | 1.00E-04 |
| **DTI_MultiQAEND** | HCP-LS | Positive | 0.997826932 | 1.00E-04 | 1.00E-04 |
| **DTI_MultiQAEND** | HCP-LS | Negative | 0.997022169 | 1.00E-04 | 1.00E-04 |
| **DTI_MultiQAEND** | HCP-LS | Absolute | 0.996843751 | 1.00E-04 | 1.00E-04 |
| **REST_MultiQAEND** | HCP-D | Positive | 0.947698487 | 1.00E-04 | 1.00E-04 |
| **REST_MultiQAEND** | HCP-D | Negative | 0.945588044 | 1.00E-04 | 1.00E-04 |
| **REST_MultiQAEND** | HCP-D | Absolute | 0.951456097 | 1.00E-04 | 1.00E-04 |
| **REST_MultiQAEND** | HCP-YA | Positive | 0.95793879 | 1.00E-04 | 1.00E-04 |
| **REST_MultiQAEND** | HCP-YA | Negative | 0.943646629 | 1.00E-04 | 1.00E-04 |
| **REST_MultiQAEND** | HCP-YA | Absolute | 0.964592591 | 1.00E-04 | 1.00E-04 |
| **REST_MultiQAEND** | HCP-A | Positive | 0.94851752 | 1.00E-04 | 1.00E-04 |
| **REST_MultiQAEND** | HCP-A | Negative | 0.942837923 | 1.00E-04 | 1.00E-04 |
| **REST_MultiQAEND** | HCP-A | Absolute | 0.961243225 | 1.00E-04 | 1.00E-04 |
| **REST_MultiQAEND** | HCP-LS | Positive | 0.98139974 | 1.00E-04 | 1.00E-04 |
| **REST_MultiQAEND** | HCP-LS | Negative | 0.965244196 | 1.00E-04 | 1.00E-04 |
| **REST_MultiQAEND** | HCP-LS | Absolute | 0.976101834 | 1.00E-04 | 1.00E-04 |
| **Network-level** | | | | | |

| Comparison | Cohort | Type | Upper Triangle r | Upper Triangle p | Upper Triangle p-FDR |
|---|---|---|---|---|---|
| **DTI_MultiQAEND** | HCP-D | Positive | 0.879468964 | 1.00E-04 | 1.00E-04 |
| **DTI_MultiQAEND** | HCP-D | Negative | 0.981550433 | 1.00E-04 | 1.00E-04 |
| **DTI_MultiQAEND** | HCP-D | Absolute | 0.975377323 | 1.00E-04 | 1.00E-04 |
| **DTI_MultiQAEND** | HCP-YA | Positive | 0.88269041 | 1.00E-04 | 1.00E-04 |
| **DTI_MultiQAEND** | HCP-YA | Negative | 0.910230388 | 1.00E-04 | 1.00E-04 |
| **DTI_MultiQAEND** | HCP-YA | Absolute | 0.914890634 | 1.00E-04 | 1.00E-04 |
| **DTI_MultiQAEND** | HCP-A | Positive | 0.99200131 | 1.00E-04 | 1.00E-04 |
| **DTI_MultiQAEND** | HCP-A | Negative | 0.982080712 | 1.00E-04 | 1.00E-04 |
| **DTI_MultiQAEND** | HCP-A | Absolute | 0.989854384 | 1.00E-04 | 1.00E-04 |
| **DTI_MultiQAEND** | HCP-LS | Positive | 0.995143842 | 1.00E-04 | 1.00E-04 |
| **DTI_MultiQAEND** | HCP-LS | Negative | 0.991809962 | 1.00E-04 | 1.00E-04 |
| **DTI_MultiQAEND** | HCP-LS | Absolute | 0.995647921 | 1.00E-04 | 1.00E-04 |
| **REST_MultiQAEND** | HCP-D | Positive | 0.944209745 | 1.00E-04 | 1.00E-04 |
| **REST_MultiQAEND** | HCP-D | Negative | 0.918848938 | 1.00E-04 | 1.00E-04 |
| **REST_MultiQAEND** | HCP-D | Absolute | 0.959067742 | 1.00E-04 | 1.00E-04 |
| **REST_MultiQAEND** | HCP-YA | Positive | 0.88113119 | 1.00E-04 | 1.00E-04 |

| | | | | | |
|---|---|---|---|---|---|
| **REST_MultiQAEND** | HCP-YA | Negative | 0.904943811 | 1.00E-04 | 1.00E-04 |
| **REST_MultiQAEND** | HCP-YA | Absolute | 0.875773456 | 1.00E-04 | 1.00E-04 |
| **REST_MultiQAEND** | HCP-A | Positive | 0.897196442 | 1.00E-04 | 1.00E-04 |
| **REST_MultiQAEND** | HCP-A | Negative | 0.933711984 | 1.00E-04 | 1.00E-04 |
| **REST_MultiQAEND** | HCP-A | Absolute | 0.929086222 | 1.00E-04 | 1.00E-04 |
| **REST_MultiQAEND** | HCP-LS | Positive | 0.977095729 | 1.00E-04 | 1.00E-04 |
| **REST_MultiQAEND** | HCP-LS | Negative | 0.957711069 | 1.00E-04 | 1.00E-04 |
| **REST_MultiQAEND** | HCP-LS | Absolute | 0.942159072 | 1.00E-04 | 1.00E-04 |

*Table S11. CPM multimodal-unimodal similarity analysis results at the edge, node, and network levels.*

**Supplementary Ridge Regression rCPM Results**

**Structural and functional connectomes predict age**

| **Modality** | **Cohort** | **Pearson r** | **Null CPM FDR-corrected p** | **Spearman ρ** | **MAE** | **MSE** | **$q^2$** |
|---|---|---|---|---|---|---|---|
| **dMRI QA End** | HCP-D | 0.649903 | 1.00E-04 | 0.659688 | 2.644274 | 10.17821 | 0.282863 |
| | HCP-YA | 0.25328 | 1.00E-04 | 0.241732 | 3.009334 | 12.83213 | 0.055087 |
| | HCP-A | 0.682911 | 1.00E-04 | 0.639399 | 9.707289 | 138.4804 | 0.213457 |
| | HCP-LS | 0.734147 | 1.00E-04 | 0.525604 | 9.949285 | 186.9745 | 0.249534 |
| **rsfMRI** | HCP-D | 0.813507 | 1.00E-04 | 0.827971 | 1.912161 | 5.587146 | 0.606341 |
| | HCP-YA | 0.316071 | 1.00E-04 | 0.328531 | 2.905649 | 12.27967 | 0.095768 |
| | HCP-A | 0.520273 | 1.00E-04 | 0.499031 | 10.06468 | 150.2893 | 0.146385 |
| | HCP-LS | 0.828302 | 1.00E-04 | 0.80081 | 8.447299 | 140.2829 | 0.436942 |
| **Multimodal dMRI QA End + rsfMRI** | HCP-D | 0.831699 | 1.00E-04 | 0.845434 | 1.855473 | 5.270029 | 0.628684 |
| | HCP-YA | 0.378124 | 1.00E-04 | 0.382751 | 2.845036 | 11.77438 | 0.132976 |
| | HCP-A | 0.725368 | 1.00E-04 | 0.701751 | 9.138547 | 124.079 | 0.295254 |
| | HCP-LS | 0.871868 | 1.00E-04 | 0.807607 | 7.86656 | 118.1583 | 0.525744 |

*Table S12. rCPM performance metrics from the median-performing diffusion MRI (dMRI) QA End, resting-state fMRI (rsfMRI), and multimodal connectome-based predictive models for each age cohort (HCP-D, HCP-YA, HCP-A, and HCP-LS). Model performance was evaluated with Pearson's correlation*

**Commented [MM2]:** Finishing null

*(r), Spearman's rank correlation (ρ), mean absolute error (MAE), mean squared error (MSE), and the predictive coefficient of determination ($q^2$).*

| Train Cohort | Test Cohort | Modality | Pearson r | p-value | FDR-corrected p | Spearman rho | MSE | MAE | $q^2$ |
|---|---|---|---|---|---|---|---|---|---|
| HCP-D | HCP-YA | dMRI QA End | 0.081950245 | 0.027564691 | 0.035676633 | 0.082118284 | 188.5875644 | 13.22516364 | -12.88692769 |
| HCP-D | HCP-A | dMRI QA End | 0.204477283 | 0.000458016 | 0.000785171 | 0.187408658 | 1727.479939 | 39.43071046 | -8.811765083 |
| HCP-D | HCP-LS | dMRI QA End | 0.520266327 | 8.24E-103 | 7.42E-102 | 0.429525458 | 435.6197379 | 14.98524556 | -0.748462216 |
| HCP-YA | HCP-D | dMRI QA End | 0.199808401 | 1.65E-05 | 3.49E-05 | 0.198550923 | 169.2372289 | 12.47194309 | -10.92412738 |
| HCP-YA | HCP-A | dMRI QA End | 0.129252486 | 0.027748492 | 0.035676633 | 0.118610119 | 949.4815845 | 27.83438792 | -4.392878982 |
| HCP-YA | HCP-LS | dMRI QA End | 0.313596737 | 6.27E-35 | 2.26E-34 | 0.536964879 | 244.5641653 | 10.64428565 | 0.018384235 |
| HCP-A | HCP-D | dMRI QA End | 0.138676017 | 0.002938794 | 0.004599851 | 0.144935654 | 1575.375051 | 39.50764856 | -109.9978751 |
| HCP-A | HCP-YA | dMRI QA End | -0.000515796 | 0.988953591 | 0.988953591 | -0.009309503 | 833.1382843 | 28.60558716 | -60.34938505 |
| HCP-A | HCP-LS | dMRI QA End | 0.386213949 | 1.59E-53 | 9.57E-53 | 0.347454524 | 923.4190127 | 28.1386105 | -2.706359269 |
| HCP-LS | HCP-D | dMRI QA End | 0.41211773 | 3.32E-20 | 1.09E-19 | 0.418286299 | 180.5419007 | 12.98880747 | -11.72063266 |
| HCP-LS | HCP-YA | dMRI QA End | 0.105658764 | 0.004454108 | 0.006413915 | 0.09952318 | 15.06294842 | 3.19410033 | -0.109182761 |
| HCP-LS | HCP-A | dMRI QA End | 0.50292216 | 5.37E-20 | 1.49E-19 | 0.494632433 | 590.8359236 | 21.16511003 | -2.355838266 |
| HCP-D | HCP-YA | rsfMRI | -0.034540518 | 0.35370956 | 0.410759489 | -0.05195615 | 141.1915076 | 11.23402498 | -9.396848074 |
| HCP-D | HCP-A | rsfMRI | -0.045336226 | 0.441828462 | 0.49705702 | -0.040756137 | 1639.984793 | 38.19893891 | -8.314808906 |

| | | | | | | | | | |
|---|---|---|---|---|---|---|---|---|---|
| HCP-D | HCP-LS | rsfMRI | 0.470094173 | 9.64E-82 | 6.94E-81 | 0.481712708 | 393.9449209 | 13.55192265 | -0.581190542 |
| HCP-YA | HCP-D | rsfMRI | -0.034772772 | 0.457868791 | 0.499493226 | -0.048216493 | 150.2344692 | 11.62591558 | -9.585229738 |
| HCP-YA | HCP-A | rsfMRI | 0.076221426 | 0.195568127 | 0.234681753 | 0.072118064 | 960.7912681 | 28.03099932 | -4.457115884 |
| HCP-YA | HCP-LS | rsfMRI | 0.343424552 | 5.60E-42 | 2.24E-41 | 0.553562972 | 240.5941632 | 10.36392266 | 0.034318771 |
| HCP-A | HCP-D | rsfMRI | -0.191019849 | 3.88E-05 | 7.34E-05 | -0.168966736 | 1512.721349 | 38.66866493 | -105.5834165 |
| HCP-A | HCP-YA | rsfMRI | 0.017505426 | 0.638412889 | 0.656653257 | 0.01940277 | 752.840308 | 27.16579751 | -54.43652333 |
| HCP-A | HCP-LS | rsfMRI | 0.531034365 | 8.26E-108 | 9.91E-107 | 0.408777543 | 864.842643 | 27.16944634 | -2.47124924 |
| HCP-LS | HCP-D | rsfMRI | 0.578921171 | 2.45E-42 | 1.10E-41 | 0.589595455 | 101.4101075 | 9.562417269 | -6.14515977 |
| HCP-LS | HCP-YA | rsfMRI | 0.176315477 | 1.84E-06 | 4.14E-06 | 0.158674149 | 15.87893397 | 3.28558939 | -0.169269079 |
| HCP-LS | HCP-A | rsfMRI | 0.50317353 | 5.11E-20 | 1.49E-19 | 0.508685739 | 449.7162436 | 17.98808566 | -1.554304704 |

*Table S13. Cross-cohort rCPM model generalizability using final rCPM models.*

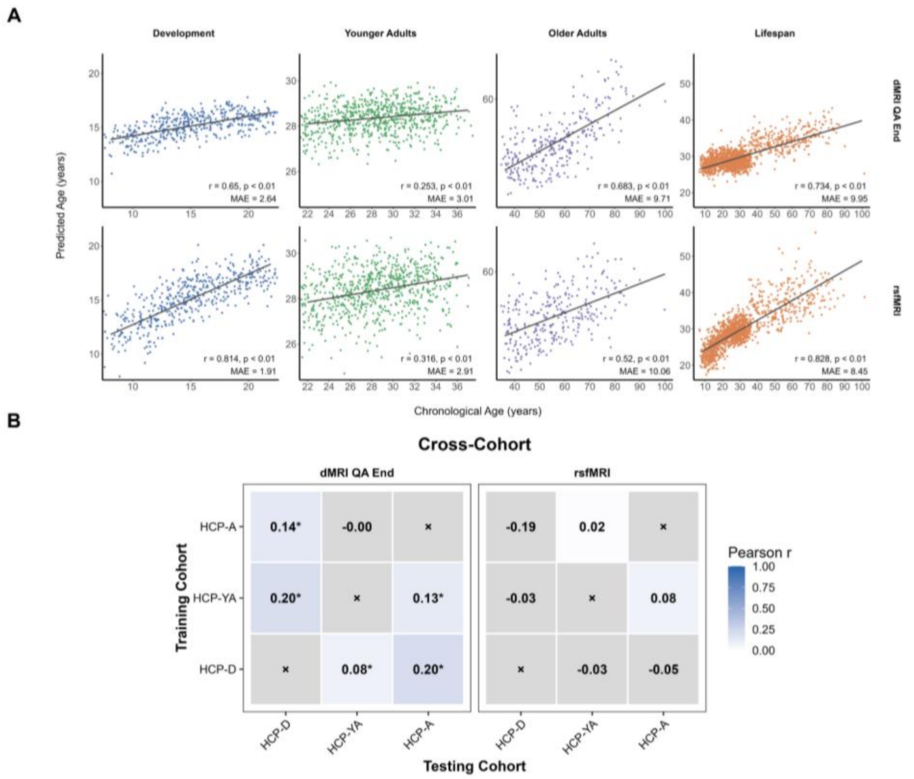

*Figure S3. Ridge regression connectome-based predictive modeling (rCPM) brain age prediction performance across age cohorts. Scatterplots show predicted age versus chronological age for the median-performing CPM model in the developmental (HCP-D), young adult (HCP-YA), older adult (HCP-A), and lifespan (HCP-LS) cohorts. The top row shows structural connectivity models derived from dMRI quantitative anisotropy (QA End), and the bottom row shows resting-state functional connectivity (rsfMRI) models. Each point represents an individual participant. Solid gray lines indicate the linear regression fit between chronological and predicted age. Insets display the Pearson correlation coefficient (r) and mean absolute error (MAE) for the median-performing model; all prediction models remained significant following permutation testing ($p < 0.0001$). Points are colored by cohort: blue = developmental, green = young adults, purple = older adults, and orange = lifespan. (B) Heatmaps summarize the generalizability of final CPM age prediction models across cohorts evaluated through external validation/cross-sex prediction. Heatmap values represent the Pearson correlation coefficient (r) between predicted and chronological age when models were applied to independent datasets. Darker shading indicates stronger age prediction performance (higher Pearson r). Asterisks indicate predictions that remained significant after false discovery rate (FDR) correction ($p < 0.05$). Gray cells marked with × indicate comparisons that were not performed (e.g., identical training and testing cohort).*

**Commented [MM3]:** Need to finish

**Exploration of age prediction model characteristics and biases**

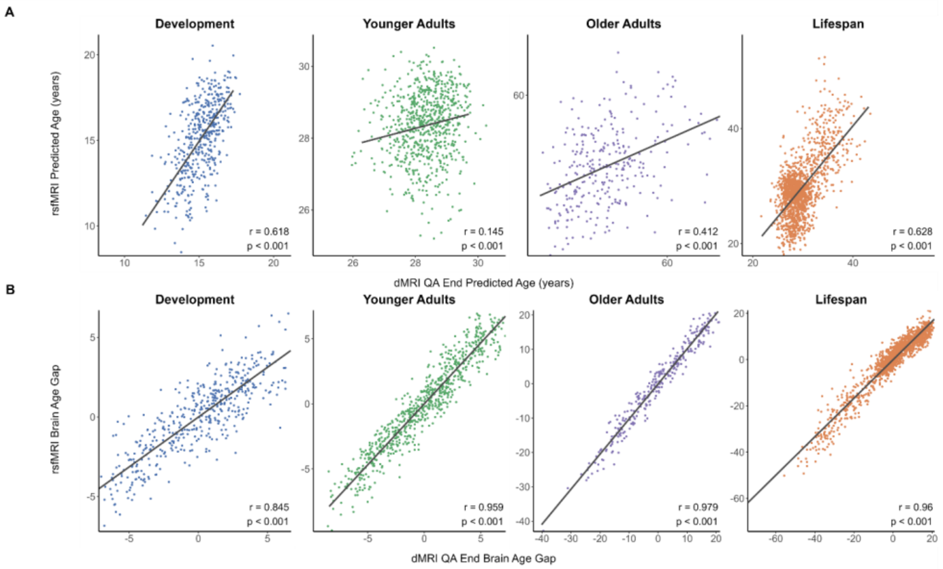

*Figure S4. (A) Correlation between structural and functional rCPM predicted ages. Scatterplots illustrate the relationship between predicted ages generated by the median-performing structural connectivity model (dMRI QA End; x-axis) and the median-performing rsfMRI model (y-axis) for the developmental (HCP-D), young adult (HCP-YA), older adult (HCP-A), and lifespan (HCP-LS) cohorts. (B) Correlation between structural and functional brain age gap (BAG) estimates. Scatterplots show the relationship between BAG (predicted age − chronological age) estimated by the median-performing structural dMRI QA End CPM model (x-axis) and the median-performing rsfMRI CPM model (y-axis) model for the developmental (HCP-D), young adult (HCP-YA), older adult (HCP-A), and lifespan (HCP-LS) cohorts. Each point represents an individual participant. Solid gray lines indicate the linear regression fit between structural and functional BAG estimates. Insets display the Pearson correlation coefficient (r) and false discovery rate (FDR)-corrected p-value for each cohort. Points are colored according to cohort (HCP-D, blue; HCP-YA, green; HCP-A, purple; HCP-LS, orange).*

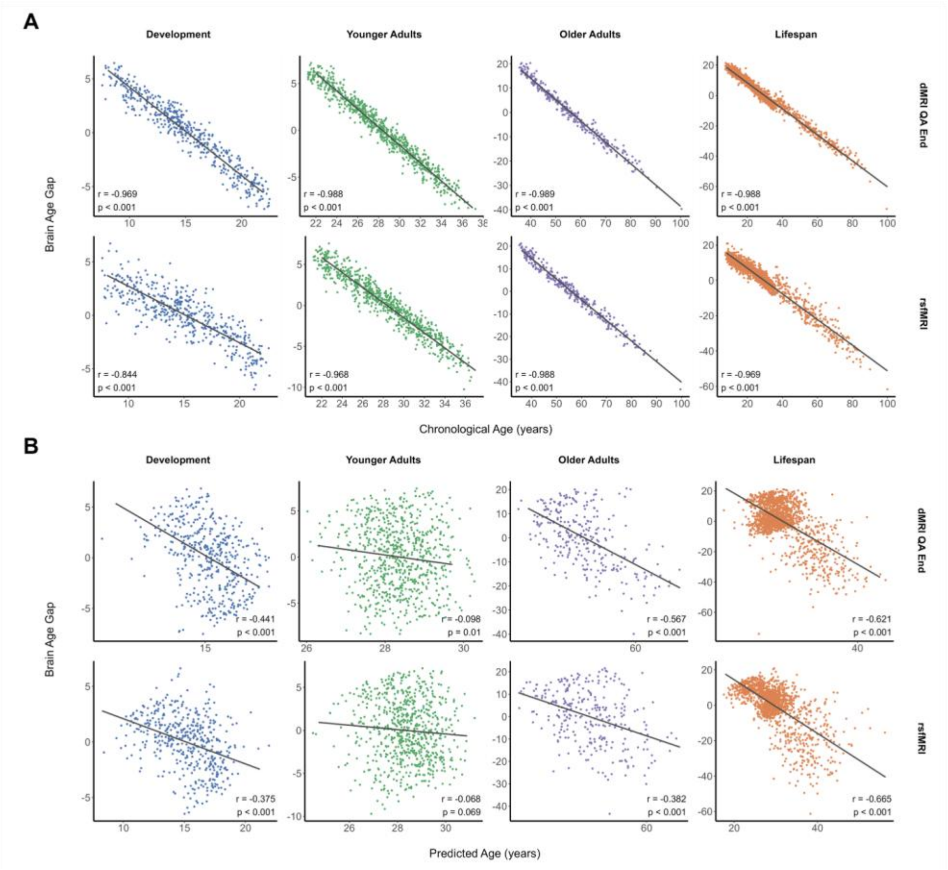

*Figure S5. (A) Relationship between brain age gap (BAG) and chronological age. Scatterplots show brain age gap (BAG; predicted age − chronological age) as a function of chronological age for the median-performing rCPM models in the developmental (HCP-D), young adult (HCP-YA), older adult (HCP-A), and lifespan (HCP-LS) cohorts. The top row shows structural connectivity models derived from dMRI QA End, and the bottom row shows rsfMRI models. (B) Relationship between brain age gap (BAG) and predicted age. Scatterplots show brain age gap (BAG; predicted age − chronological age; y-axis) as a function of CPM predicted age (x-axis) for the median-performing structural and functional CPM models in the developmental (HCP-D), young adult (HCP-YA), older adult (HCP-A), and lifespan (HCP-LS) cohorts. The top row shows dMRI QA End models, and the bottom row shows rsfMRI models. Each point represents an individual participant. Solid gray lines indicate the linear regression fit between predicted age and BAG. Insets display the Pearson correlation coefficient (r) and false discovery rate (FDR)-corrected p-value for each cohort and modality. Points are colored according to cohort (HCP-D, blue; HCP-YA, green; HCP-A, purple; HCP-LS, orange).*

**Density of predictive networks from the median-performing rCPM models**

| Age Cohort | DTI Total Edges | Rest Total Edges |
| --- | --- | --- |

| | | |
|---|---|---|
| **HCP-Development** | 8,397 | 20,398 |
| **HCP-Young Adults** | 5,729 | 10,311 |
| **HCP-Aging** | 8,613 | 10,872 |
| **HCP-Lifespan** | 15,398 | 26,701 |
| *HCP, Human Connectome Project* | | |

*Table S14. Total edge density for each imaging modality in age prediction from the median performing rCPM models*

**Structural-functional rCPM coefficient convergence**

At the edge level, structural and functional rCPM coefficient vectors exhibited minimal correspondence across all cohorts. Correlations between signed ridge coefficients were near zero in the developmental ($r = 0.024$), young adult ($r = -0.004$), older adult ($r = -0.016$), and lifespan ($r = -0.029$) cohorts, indicating little agreement in the direction or magnitude of corresponding edge weights between modalities. Although several comparisons reached statistical significance following permutation testing, effect sizes were negligible ($|r| < 0.03$), reflecting the large number of edges included in the analysis rather than meaningful coefficient similarity. Comparisons based on absolute coefficient magnitudes likewise demonstrated limited convergence. Absolute coefficient correlations were consistently negative across all cohorts ($r = -0.09$ to $-0.20$; all FDR-corrected $p < 0.001$), indicating that edges assigned relatively large predictive weights in one modality tended to receive smaller weights in the other. Together, these findings suggest that structural and functional rCPM models rely on distinct edge-level weighting patterns despite achieving similar age-prediction performance.

Node-level analyses demonstrated modest structural–functional convergence in regional predictive contributions. Regional contribution was quantified as the sum of the absolute ridge coefficients incident upon each node. Positive correlations between structural and functional node contribution profiles were observed in the developmental ($r = 0.167$, FDR-corrected $p = 0.014$), young adult ($r = 0.152$, FDR-corrected $p = 0.015$), and lifespan cohorts ($r = 0.189$, FDR-corrected $p = 0.006$), indicating that brain regions contributing strongly to age prediction in one modality also tended to contribute strongly in the other. In contrast, node-level convergence did not reach significance in the older adult cohort. Additionally, across all cohorts, the mean regional contribution was greater for structural than functional coefficient networks, reflecting larger overall ridge coefficient magnitudes within the structural models.

Network-level analyses compared the distribution of rCPM coefficients across the ten canonical Shen brain systems. Correlations between signed structural and functional network contribution matrices were generally weak, with no significant convergence observed in the developmental, young adult, or lifespan cohorts. In contrast, the older adult cohort demonstrated a significant negative correlation between signed network contribution matrices ($r = -0.522$, FDR-corrected $p = 0.006$), indicating that canonical network pairs receiving stronger signed coefficients in one modality tended to receive weaker or oppositely directed coefficients in the other.

Comparisons based on absolute coefficient magnitudes likewise showed no significant structural–functional convergence across canonical brain systems (all FDR-corrected $p > 0.17$). Together, these findings suggest that although structural and functional models may emphasize similar brain regions, they organize predictive information differently across large-scale network interactions.

| **Edge-level** | | | | |
|---|---|---|---|---|
| **Cohort** | **Type** | **Edge Correlation r** | **Permutation p** | **Permutation p-FDR** |
| **HCP-D** | Signed | 0.023648 | 0.0005 | 0.000667 |
| | Absolute | -0.12564 | 1.00E-04 | 0.00016 |
| **HCP-YA** | Signed | -0.00391 | 0.647935 | 0.647935 |

| | | | | |
|---|---|---|---|---|
| | Absolute | -0.14804 | 1.00E-04 | 0.00016 |
| **HCP-A** | Signed | -0.01586 | 0.041196 | 0.047081 |
| | Absolute | -0.20177 | 1.00E-04 | 0.00016 |
| **HCP-LS** | Signed | -0.02855 | 1.00E-04 | 0.00016 |
| | Absolute | -0.09048 | 1.00E-04 | 0.00016 |
| | | **Node-level** | | |
| **Cohort** | **Type** | **Node Correlation r** | **Permutation p** | **Permutation p-FDR** |
| **HCP-D** | Absolute | 0.16667 | 0.007099 | 0.014199 |
| **HCP-YA** | Absolute | 0.151511 | 0.011399 | 0.015198 |
| **HCP-A** | Absolute | 0.08198 | 0.175382 | 0.175382 |
| **HCP-LS** | Absolute | 0.188954 | 0.0014 | 0.005599 |
| | | **Network-level** | | |
| **Cohort** | **Type** | **Network Contribution r** | **Permutation p** | **Permutation p-FDR** |
| **HCP-D** | Signed | 0.014959 | 0.925207 | 0.925207 |
| | Absolute | 0.404167 | 0.059394 | 0.175182 |
| **HCP-YA** | Signed | 0.039289 | 0.791021 | 0.904024 |
| | Absolute | -0.26685 | 0.093291 | 0.186581 |
| **HCP-A** | Signed | -0.52172 | 0.0007 | 0.005599 |
| | Absolute | 0.155703 | 0.568043 | 0.757391 |
| **HCP-LS** | Signed | -0.27686 | 0.065693 | 0.175182 |
| | Absolute | 0.354797 | 0.206779 | 0.330847 |

*Table S15. rCPM structural-functional convergence analysis results at the edge, node, and network levels.*

| **Cohort** | **Train Modality** | **Test Modality** | **Pearson r** | **p-value** | **FDR-corrected p** | **Spearman rho** | **MSE** | **MAE** | **$q^2$** |
|---|---|---|---|---|---|---|---|---|---|
| HCP-D | dMRI QA End | rsfMRI | 0.351706 | 8.81E-15 | 2.64E-14 | 0.345637 | 12.88842 | 3.012596 | 0.091907 |
| HCP-D | rsfMRI | dMRI QA End | 0.107006 | 0.022002 | 0.027792 | 0.1114 | 14.11742 | 3.139406 | 0.005314 |
| HCP-YA | dMRI QA End | rsfMRI | -0.04908 | 0.187447 | 0.224937 | -0.04518 | 15.03144 | 3.216702 | -0.10686 |
| HCP-YA | rsfMRI | dMRI QA End | 0.044962 | 0.227249 | 0.259713 | 0.048641 | 20.23277 | 3.685273 | -0.48987 |
| HCP-A | dMRI QA End | rsfMRI | -0.1982 | 0.000688 | 0.001032 | -0.17208 | 213.8743 | 11.8113 | -0.21477 |
| HCP-A | rsfMRI | dMRI QA End | -0.38828 | 7.16E-12 | 1.91E-11 | -0.3565 | 181.5314 | 11.27967 | -0.03106 |
| HCP-LS | dMRI QA End | rsfMRI | -0.10467 | 5.76E-05 | 9.88E-05 | -0.24301 | 262.5958 | 11.94236 | -0.05399 |

| | | | | | | | | | |
|---|---|---|---|---|---|---|---|---|---|
| HCP-LS | rsfMRI | dMRI QA End | -0.14416 | 2.80E-08 | 5.61E-08 | -0.09543 | 250.6593 | 11.52805 | -0.00608 |

*Table S16. Cross-modality rCPM model generalizability using the final rCPM models.*

**Brain age prediction from individual canonical networks**

**Commented [MM4]:** Finishing and building figure and tables

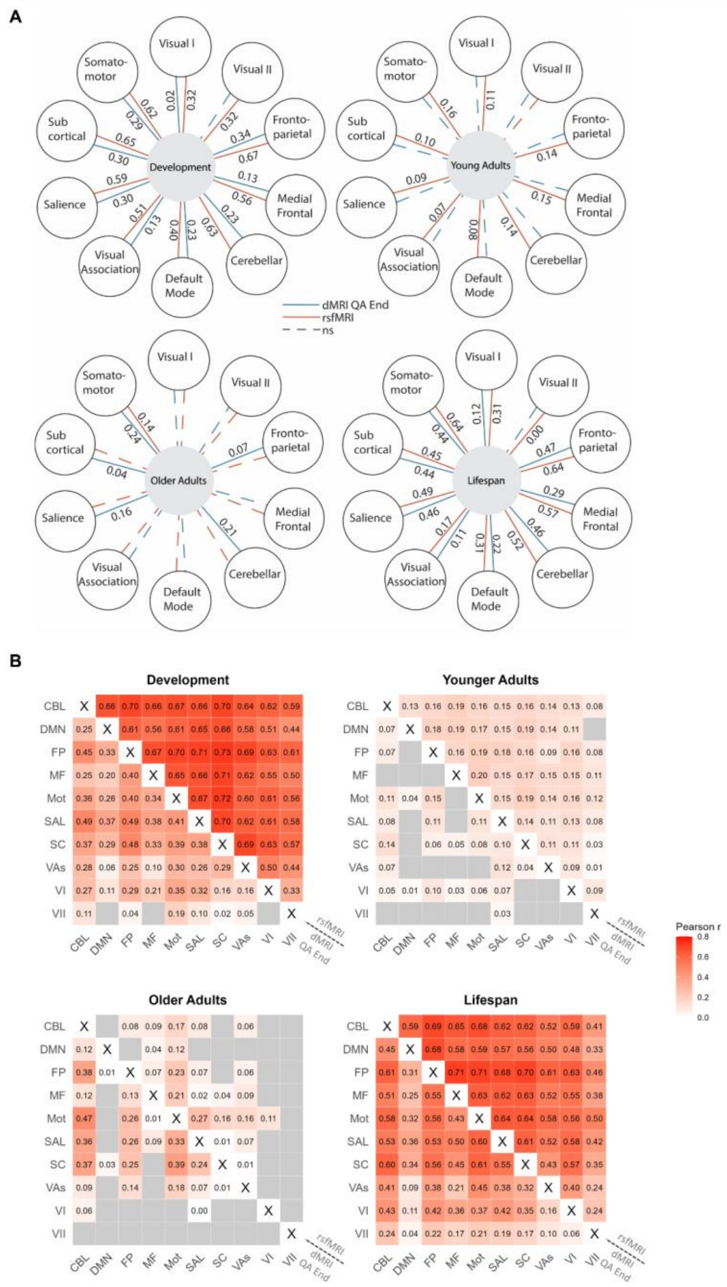
A
Development
Young Adults
Older Adults
Lifespan
Visual I
Visual II
Fronto-parietal
Medial Frontal
Cerebellar
Default Mode
Visual Association
Salience
Sub cortical
Somato-motor
dMRI QA End
rsfMRI
ns
B
Development
Younger Adults
Older Adults
Lifespan
CBL
DMN
FP
MF
Mot
SAL
SC
VAs
VI
VII
rsfMRI
dMRI QA End
Pearson r
0.8
0.6
0.4
0.2
0.0

*Figure S6. Individual network-level rCPM age prediction performance across the lifespan. (A) Individual within-network rCPM age prediction performance. Flower plots display the median-performing individual within-network CPM age prediction models for each cohort: developmental (top left), young adult (top right), older adult (bottom left), and lifespan (bottom right). Petals represent the 10 Shen 268-node atlas networks, the center denotes the age cohort, and stems indicate the Pearson correlation coefficient (r) between predicted and chronological age for each network-specific model. dMRI QA End models are shown with blue lines, and rsfMRI models are shown with red lines. Solid lines indicate network-level models that achieved significance following permutation testing against the null distribution, whereas dashed lines indicate non-significant models. The plots highlight modality- and network-specific variation in rCPM age prediction performance across age groups. (B) Individual between-network rCPM age prediction performance. Heatmaps display Pearson correlation coefficients (r) for significant between-network rCPM age prediction models after FDR correction across developmental, young adult, older adult, and lifespan cohorts. The upper triangle represents rsfMRI models, and the lower triangle represents dMRI QA End models. The diagonal is excluded. Non-significant connections are shown in gray, and significant connections are color-coded according to predictive performance (Pearson r). Values within each cell indicate the corresponding Pearson correlation coefficient. Network abbreviations: CBL = cerebellar, DMN = default mode network, FP = frontoparietal, MF = medial frontal, Mot = somatomotor, SAL = salience, SC = subcortical, VAs = visual association, VI = visual I, VII = visual II.*

| Between-Network | | | | | | | | |
|---|---|---|---|---|---|---|---|---|
| **Cohort** | **Modality** | **Network** | **Pearson r** | **pFDR** | **Spearman rho** | **MSE** | **MAE** | **q2** |
| **HCP-D** | dMRI QA End | CBL-DMN | 0.253943 | 0.001215 | 0.276827 | 13.9584 | 3.121065 | 0.016518 |
| **HCP-D** | dMRI QA End | CBL-FP | 0.448661 | 0.001215 | 0.449005 | 13.7184 | 3.098106 | 0.033428 |
| **HCP-D** | dMRI QA End | CBL-MF | 0.246062 | 0.001215 | 0.274958 | 14.01761 | 3.129574 | 0.012346 |
| **HCP-D** | dMRI QA End | CBL-Mot | 0.355786 | 0.001215 | 0.348533 | 13.62574 | 3.085613 | 0.039957 |
| **HCP-D** | dMRI QA End | CBL-SAL | 0.486188 | 0.001215 | 0.489579 | 13.65335 | 3.089389 | 0.038012 |
| **HCP-D** | dMRI QA End | CBL-SC | 0.365558 | 0.001215 | 0.397479 | 13.79273 | 3.101405 | 0.028191 |
| **HCP-D** | dMRI QA End | CBL-VAs | 0.27625 | 0.001215 | 0.286395 | 13.99979 | 3.128423 | 0.013602 |
| **HCP-D** | dMRI QA End | CBL-VII | 0.110532 | 0.002305 | 0.104677 | 14.13555 | 3.145716 | 0.004037 |
| **HCP-D** | dMRI QA End | CBL-VI | 0.265671 | 0.001215 | 0.249078 | 14.00324 | 3.12828 | 0.013359 |
| **HCP-D** | dMRI QA End | DMN-FP | 0.329761 | 0.001215 | 0.31303 | 13.96484 | 3.124213 | 0.016065 |
| **HCP-D** | dMRI QA End | DMN-MF | 0.204252 | 0.001215 | 0.182895 | 14.0656 | 3.134055 | 0.008965 |
| **HCP-D** | dMRI QA End | DMN-Mot | 0.26256 | 0.001215 | 0.274383 | 13.99401 | 3.126587 | 0.014009 |

| | | | | | | | | |
|---|---|---|---|---|---|---|---|---|
| **HCP-D** | dMRI QA End | DMN-SAL | 0.365799 | 0.001215 | 0.386667 | 13.91712 | 3.120959 | 0.019427 |
| **HCP-D** | dMRI QA End | DMN-SC | 0.289228 | 0.001215 | 0.293406 | 13.98434 | 3.121419 | 0.01469 |
| **HCP-D** | dMRI QA End | DMN-VAs | 0.056426 | 0.001215 | 0.083837 | 14.1676 | 3.14564 | 0.001779 |
| **HCP-D** | dMRI QA End | DMN-VII | -0.0376 | 0.021456 | -0.04008 | 14.2146 | 3.153738 | -0.00153 |
| **HCP-D** | dMRI QA End | DMN-VI | 0.110295 | 0.002305 | 0.094463 | 14.13409 | 3.141031 | 0.004139 |
| **HCP-D** | dMRI QA End | FP-MF | 0.404845 | 0.001215 | 0.402512 | 13.80949 | 3.106625 | 0.02701 |
| **HCP-D** | dMRI QA End | FP-Mot | 0.400176 | 0.001215 | 0.402332 | 13.67691 | 3.090274 | 0.036351 |
| **HCP-D** | dMRI QA End | FP-SAL | 0.489834 | 0.001215 | 0.491111 | 13.61919 | 3.084158 | 0.040418 |
| **HCP-D** | dMRI QA End | FP-SC | 0.484956 | 0.001215 | 0.48438 | 13.63071 | 3.083396 | 0.039606 |
| **HCP-D** | dMRI QA End | FP-VAs | 0.248404 | 0.001215 | 0.246375 | 14.00727 | 3.133924 | 0.013075 |
| **HCP-D** | dMRI QA End | FP-VII | 0.039874 | 0.003289 | 0.032887 | 14.17777 | 3.148254 | 0.001062 |
| **HCP-D** | dMRI QA End | FP-VI | 0.294246 | 0.001215 | 0.319061 | 13.98308 | 3.128013 | 0.014779 |
| **HCP-D** | dMRI QA End | MF-Mot | 0.339207 | 0.001215 | 0.342393 | 13.85511 | 3.110098 | 0.023796 |
| **HCP-D** | dMRI QA End | MF-SAL | 0.375299 | 0.001215 | 0.379733 | 13.87082 | 3.109791 | 0.022689 |
| **HCP-D** | dMRI QA End | MF-SC | 0.325655 | 0.001215 | 0.326678 | 13.97537 | 3.124172 | 0.015322 |
| **HCP-D** | dMRI QA End | MF-VAs | 0.098182 | 0.001215 | 0.075318 | 14.14288 | 3.1443 | 0.00352 |
| **HCP-D** | dMRI QA End | MF-VII | -0.00865 | 0.018818 | -0.02279 | 14.20235 | 3.147996 | -0.00067 |
| **HCP-D** | dMRI QA End | MF-VI | 0.206125 | 0.001215 | 0.195649 | 14.07288 | 3.138364 | 0.008452 |
| **HCP-D** | dMRI QA End | Mot-SAL | 0.410487 | 0.001215 | 0.409371 | 13.60618 | 3.0842 | 0.041335 |
| **HCP-D** | dMRI QA End | Mot-SC | 0.389542 | 0.001215 | 0.397455 | 13.69385 | 3.094296 | 0.035158 |
| **HCP-D** | dMRI QA End | Mot-VAs | 0.296594 | 0.001215 | 0.318232 | 13.75984 | 3.104871 | 0.030508 |
| **HCP-D** | dMRI QA End | Mot-VII | 0.189306 | 0.001215 | 0.204556 | 14.05712 | 3.138283 | 0.009563 |
| **HCP-D** | dMRI QA End | Mot-VI | 0.352807 | 0.001215 | 0.316887 | 13.91517 | 3.12221 | 0.019564 |

| | | | | | | | | |
|---|---|---|---|---|---|---|---|---|
| **HCP-D** | dMRI QA End | SAL-SC | 0.376641 | 0.001215 | 0.368346 | 13.92415 | 3.118341 | 0.018931 |
| **HCP-D** | dMRI QA End | SAL-VAs | 0.261707 | 0.001215 | 0.245802 | 14.00981 | 3.13049 | 0.012896 |
| **HCP-D** | dMRI QA End | SAL-VII | 0.096994 | 0.001215 | 0.090714 | 14.14221 | 3.143857 | 0.003567 |
| **HCP-D** | dMRI QA End | SAL-VI | 0.315243 | 0.001215 | 0.306373 | 13.9301 | 3.120826 | 0.018512 |
| **HCP-D** | dMRI QA End | SC-VAs | 0.288282 | 0.001215 | 0.269005 | 14.00943 | 3.131355 | 0.012923 |
| **HCP-D** | dMRI QA End | SC-VII | 0.015106 | 0.004281 | 0.021819 | 14.18974 | 3.150374 | 0.000218 |
| **HCP-D** | dMRI QA End | SC-VI | 0.157415 | 0.001215 | 0.145454 | 14.10693 | 3.141494 | 0.006053 |
| **HCP-D** | dMRI QA End | VAs-VII | 0.048795 | 0.001215 | 0.056365 | 14.17173 | 3.150253 | 0.001487 |
| **HCP-D** | dMRI QA End | VAs-VI | 0.161495 | 0.003289 | 0.153323 | 14.10307 | 3.139677 | 0.006325 |
| **HCP-D** | dMRI QA End | VI-VII | -0.11228 | 0.303696 | -0.0878 | 14.25782 | 3.154313 | -0.00458 |
| **HCP-YA** | dMRI QA End | CBL-DMN | 0.06907 | 0.013832 | 0.057931 | 13.54638 | 3.100888 | 0.002492 |
| **HCP-YA** | dMRI QA End | CBL-FP | 0.073956 | 0.02023 | 0.075496 | 13.53197 | 3.099287 | 0.003553 |
| **HCP-YA** | dMRI QA End | CBL-MF | 0.025163 | 0.081265 | -0.00102 | 13.57188 | 3.101643 | 0.000615 |
| **HCP-YA** | dMRI QA End | CBL-Mot | 0.113881 | 0.012844 | 0.118114 | 13.4894 | 3.091026 | 0.006688 |
| **HCP-YA** | dMRI QA End | CBL-SAL | 0.080549 | 0.013832 | 0.093703 | 13.53322 | 3.098662 | 0.003461 |
| **HCP-YA** | dMRI QA End | CBL-SC | 0.137214 | 0.011239 | 0.130562 | 13.48632 | 3.091457 | 0.006914 |
| **HCP-YA** | dMRI QA End | CBL-VAs | 0.070866 | 0.013832 | 0.054382 | 13.54755 | 3.099895 | 0.002406 |
| **HCP-YA** | dMRI QA End | CBL-VII | -0.06754 | 0.280969 | -0.08717 | 13.61202 | 3.106803 | -0.00234 |
| **HCP-YA** | dMRI QA End | CBL-VI | 0.045838 | 0.016562 | 0.031688 | 13.56073 | 3.099997 | 0.001435 |
| **HCP-YA** | dMRI QA End | DMN-FP | -0.03143 | 0.208213 | -0.03162 | 13.59914 | 3.105778 | -0.00139 |
| **HCP-YA** | dMRI QA End | DMN-MF | -0.07375 | 0.395186 | -0.06421 | 13.61664 | 3.108082 | -0.00268 |
| **HCP-YA** | dMRI QA End | DMN-Mot | 0.043514 | 0.040868 | 0.036233 | 13.56166 | 3.100484 | 0.001367 |
| **HCP-YA** | dMRI QA End | DMN-SAL | -0.01987 | 0.158118 | -0.02839 | 13.59258 | 3.104228 | -0.00091 |

| | | | | | | | | |
|---|---|---|---|---|---|---|---|---|
| **HCP-YA** | dMRI QA End | DMN-SC | -0.03543 | 0.208213 | -0.04047 | 13.60543 | 3.105319 | -0.00186 |
| **HCP-YA** | dMRI QA End | DMN-VAs | -0.04284 | 0.183907 | -0.04305 | 13.60236 | 3.105142 | -0.00163 |
| **HCP-YA** | dMRI QA End | DMN-VII | -0.10514 | 0.55683 | -0.08664 | 13.61799 | 3.107387 | -0.00278 |
| **HCP-YA** | dMRI QA End | DMN-VI | 0.012254 | 0.041046 | 0.024342 | 13.57849 | 3.099952 | 0.000128 |
| **HCP-YA** | dMRI QA End | FP-MF | 0.023107 | 0.076798 | 0.020668 | 13.57307 | 3.10057 | 0.000527 |
| **HCP-YA** | dMRI QA End | FP-Mot | 0.154668 | 0.011239 | 0.157474 | 13.44928 | 3.090558 | 0.009642 |
| **HCP-YA** | dMRI QA End | FP-SAL | 0.106621 | 0.012844 | 0.087335 | 13.52158 | 3.095799 | 0.004318 |
| **HCP-YA** | dMRI QA End | FP-SC | 0.060377 | 0.014048 | 0.05095 | 13.54954 | 3.097909 | 0.002259 |
| **HCP-YA** | dMRI QA End | FP-VAs | -0.00619 | 0.097938 | -0.03093 | 13.58785 | 3.102858 | -0.00056 |
| **HCP-YA** | dMRI QA End | FP-VII | -0.04492 | 0.183907 | -0.04902 | 13.6019 | 3.101979 | -0.0016 |
| **HCP-YA** | dMRI QA End | FP-VI | 0.098867 | 0.013832 | 0.111884 | 13.52744 | 3.097285 | 0.003886 |
| **HCP-YA** | dMRI QA End | MF-Mot | -0.0302 | 0.243218 | -0.03634 | 13.60576 | 3.105612 | -0.00188 |
| **HCP-YA** | dMRI QA End | MF-SAL | -0.02966 | 0.208213 | -0.01542 | 13.6006 | 3.105424 | -0.0015 |
| **HCP-YA** | dMRI QA End | MF-SC | 0.053121 | 0.014985 | 0.04593 | 13.55632 | 3.09907 | 0.00176 |
| **HCP-YA** | dMRI QA End | MF-VAs | -0.0338 | 0.178322 | -0.03116 | 13.5986 | 3.106684 | -0.00135 |
| **HCP-YA** | dMRI QA End | MF-VII | -0.11885 | 0.683317 | -0.11028 | 13.63045 | 3.109415 | -0.0037 |
| **HCP-YA** | dMRI QA End | MF-VI | 0.026238 | 0.036392 | 0.014611 | 13.57225 | 3.103179 | 0.000587 |
| **HCP-YA** | dMRI QA End | Mot-SAL | 0.112342 | 0.012844 | 0.095722 | 13.49368 | 3.096057 | 0.006373 |
| **HCP-YA** | dMRI QA End | Mot-SC | 0.082887 | 0.014985 | 0.066905 | 13.52875 | 3.095852 | 0.00379 |
| **HCP-YA** | dMRI QA End | Mot-VAs | -0.0098 | 0.183907 | -0.01302 | 13.59345 | 3.102083 | -0.00097 |
| **HCP-YA** | dMRI QA End | Mot-VII | -0.0402 | 0.208213 | -0.0421 | 13.60288 | 3.102408 | -0.00167 |
| **HCP-YA** | dMRI QA End | Mot-VI | 0.062182 | 0.014048 | 0.045532 | 13.54968 | 3.101487 | 0.002249 |
| **HCP-YA** | dMRI QA End | SAL-SC | 0.097737 | 0.011239 | 0.0936 | 13.521 | 3.096996 | 0.004361 |

| | | | | | | | | |
|---|---|---|---|---|---|---|---|---|
| **HCP-YA** | dMRI QA End | SAL-VAs | 0.117429 | 0.011239 | 0.111623 | 13.52803 | 3.096283 | 0.003843 |
| **HCP-YA** | dMRI QA End | SAL-VII | 0.030148 | 0.014048 | 0.006172 | 13.57053 | 3.10247 | 0.000713 |
| **HCP-YA** | dMRI QA End | SAL-VI | 0.069136 | 0.013832 | 0.080228 | 13.54709 | 3.099058 | 0.00244 |
| **HCP-YA** | dMRI QA End | SC-VAs | 0.044748 | 0.013832 | 0.041779 | 13.56385 | 3.098723 | 0.001206 |
| **HCP-YA** | dMRI QA End | SC-VII | -0.05124 | 0.195687 | -0.07325 | 13.60526 | 3.103903 | -0.00184 |
| **HCP-YA** | dMRI QA End | SC-VI | 0.004027 | 0.081265 | 0.002099 | 13.5835 | 3.104552 | -0.00024 |
| **HCP-YA** | dMRI QA End | VAs-VII | -0.02486 | 0.08325 | -0.04028 | 13.5933 | 3.103508 | -0.00096 |
| **HCP-YA** | dMRI QA End | VAs-VI | -0.06767 | 0.287274 | -0.06295 | 13.60786 | 3.106162 | -0.00204 |
| **HCP-YA** | dMRI QA End | VI-VII | -0.08777 | 0.395186 | -0.09051 | 13.62148 | 3.106261 | -0.00304 |
| **HCP-A** | dMRI QA End | CBL-DMN | 0.115548 | 0.001955 | 0.091505 | 175.2409 | 10.95744 | 0.004664 |
| **HCP-A** | dMRI QA End | CBL-FP | 0.376248 | 0.001955 | 0.353945 | 172.6212 | 10.86719 | 0.019544 |
| **HCP-A** | dMRI QA End | CBL-MF | 0.117597 | 0.001955 | 0.10094 | 175.2379 | 10.94704 | 0.004681 |
| **HCP-A** | dMRI QA End | CBL-Mot | 0.465676 | 0.001955 | 0.407929 | 171.0181 | 10.83167 | 0.028649 |
| **HCP-A** | dMRI QA End | CBL-SAL | 0.363014 | 0.001955 | 0.318653 | 172.3912 | 10.85854 | 0.02085 |
| **HCP-A** | dMRI QA End | CBL-SC | 0.372053 | 0.001955 | 0.341924 | 172.5897 | 10.87135 | 0.019723 |
| **HCP-A** | dMRI QA End | CBL-VAs | 0.087459 | 0.001955 | 0.149948 | 175.5161 | 10.94824 | 0.003101 |
| **HCP-A** | dMRI QA End | CBL-VII | -0.06048 | 0.005619 | -0.05705 | 176.5666 | 10.99282 | -0.00287 |
| **HCP-A** | dMRI QA End | CBL-VI | 0.060085 | 0.001955 | 0.041685 | 175.6972 | 10.95086 | 0.002072 |
| **HCP-A** | dMRI QA End | DMN-FP | 0.005731 | 0.001955 | 0.007053 | 176.0905 | 10.97775 | -0.00016 |
| **HCP-A** | dMRI QA End | DMN-MF | -0.08231 | 0.024975 | -0.06775 | 176.7258 | 11.00932 | -0.00377 |
| **HCP-A** | dMRI QA End | DMN-Mot | -0.07634 | 0.024975 | -0.00787 | 176.6477 | 10.98563 | -0.00333 |
| **HCP-A** | dMRI QA End | DMN-SAL | -0.01763 | 0.004496 | -0.02943 | 176.2571 | 10.99213 | -0.00111 |
| **HCP-A** | dMRI QA End | DMN-SC | 0.034733 | 0.001955 | 0.034135 | 175.8812 | 10.96601 | 0.001027 |

| | | | | | | | | |
|---|---|---|---|---|---|---|---|---|
| **HCP-A** | dMRI QA End | DMN-VAs | -0.11188 | 0.100284 | -0.06631 | 176.923 | 11.00101 | -0.00489 |
| **HCP-A** | dMRI QA End | DMN-VII | -0.14752 | 0.31162 | -0.10763 | 177.1611 | 11.00996 | -0.00624 |
| **HCP-A** | dMRI QA End | DMN-VI | -0.12149 | 0.129383 | -0.09463 | 176.9776 | 11.00794 | -0.0052 |
| **HCP-A** | dMRI QA End | FP-MF | 0.132477 | 0.001955 | 0.177215 | 175.1438 | 10.93466 | 0.005216 |
| **HCP-A** | dMRI QA End | FP-Mot | 0.262056 | 0.001955 | 0.220716 | 174.063 | 10.92317 | 0.011354 |
| **HCP-A** | dMRI QA End | FP-SAL | 0.256571 | 0.001955 | 0.251918 | 173.9454 | 10.91273 | 0.012022 |
| **HCP-A** | dMRI QA End | FP-SC | 0.252158 | 0.001955 | 0.21008 | 174.0647 | 10.89669 | 0.011345 |
| **HCP-A** | dMRI QA End | FP-VAs | 0.135297 | 0.001955 | 0.105333 | 175.1665 | 10.94612 | 0.005087 |
| **HCP-A** | dMRI QA End | FP-VII | -0.03211 | 0.003211 | -0.00123 | 176.3614 | 10.97423 | -0.0017 |
| **HCP-A** | dMRI QA End | FP-VI | -0.01302 | 0.004496 | -0.00751 | 176.2248 | 10.98697 | -0.00092 |
| **HCP-A** | dMRI QA End | MF-Mot | 0.006953 | 0.003211 | 0.077576 | 176.0817 | 10.96781 | -0.00011 |
| **HCP-A** | dMRI QA End | MF-SAL | 0.093617 | 0.001955 | 0.071539 | 175.4389 | 10.96069 | 0.00354 |
| **HCP-A** | dMRI QA End | MF-SC | -0.03256 | 0.003211 | -0.02558 | 176.3806 | 10.98593 | -0.00181 |
| **HCP-A** | dMRI QA End | MF-VAs | -0.05693 | 0.005619 | 0.001747 | 176.506 | 10.98404 | -0.00252 |
| **HCP-A** | dMRI QA End | MF-VII | -0.1306 | 0.175638 | -0.09818 | 177.0147 | 11.00775 | -0.00541 |
| **HCP-A** | dMRI QA End | MF-VI | -0.08212 | 0.013623 | -0.05932 | 176.7133 | 11.00951 | -0.0037 |
| **HCP-A** | dMRI QA End | Mot-SAL | 0.329549 | 0.001955 | 0.328935 | 173.2984 | 10.87791 | 0.015697 |
| **HCP-A** | dMRI QA End | Mot-SC | 0.394253 | 0.001955 | 0.349359 | 172.1913 | 10.85779 | 0.021985 |
| **HCP-A** | dMRI QA End | Mot-VAs | 0.183916 | 0.001955 | 0.17921 | 174.5038 | 10.93637 | 0.008851 |
| **HCP-A** | dMRI QA End | Mot-VII | -0.07806 | 0.024975 | -0.04782 | 176.6929 | 10.987 | -0.00358 |
| **HCP-A** | dMRI QA End | Mot-VI | -0.02849 | 0.003211 | -0.00445 | 176.3503 | 10.98414 | -0.00164 |
| **HCP-A** | dMRI QA End | SAL-SC | 0.236 | 0.001955 | 0.196781 | 174.2663 | 10.93797 | 0.0102 |
| **HCP-A** | dMRI QA End | SAL-VAs | 0.071971 | 0.001955 | 0.071252 | 175.6125 | 10.95036 | 0.002553 |

| | | | | | | | | |
|---|---|---|---|---|---|---|---|---|
| **HCP-A** | dMRI QA End | SAL-VII | -0.10073 | 0.06804 | -0.07256 | 176.8361 | 10.97856 | -0.0044 |
| **HCP-A** | dMRI QA End | SAL-VI | 0.003628 | 0.001955 | 0.015463 | 176.1098 | 10.9884 | -0.00027 |
| **HCP-A** | dMRI QA End | SC-VAs | 0.010149 | 0.001955 | 0.057834 | 176.0594 | 10.97847 | 1.51E-05 |
| **HCP-A** | dMRI QA End | SC-VII | -0.11912 | 0.128122 | -0.1143 | 176.9753 | 11.01731 | -0.00519 |
| **HCP-A** | dMRI QA End | SC-VI | -0.01591 | 0.003211 | 0.004073 | 176.2396 | 10.97642 | -0.00101 |
| **HCP-A** | dMRI QA End | VAs-VII | -0.11136 | 0.07808 | -0.067 | 176.9148 | 10.99125 | -0.00484 |
| **HCP-A** | dMRI QA End | VAs-VI | -0.12575 | 0.14985 | -0.08986 | 177.003 | 10.99331 | -0.00534 |
| **HCP-A** | dMRI QA End | VI-VII | -0.16417 | 0.473526 | -0.17749 | 177.1921 | 11.02922 | -0.00642 |
| **HCP-LS** | dMRI QA End | CBL-DMN | 0.45436 | 0.000999 | 0.307487 | 245.9409 | 11.25211 | 0.012858 |
| **HCP-LS** | dMRI QA End | CBL-FP | 0.608367 | 0.000999 | 0.393209 | 240.9175 | 11.15201 | 0.033021 |
| **HCP-LS** | dMRI QA End | CBL-MF | 0.507501 | 0.000999 | 0.298777 | 245.1047 | 11.23368 | 0.016215 |
| **HCP-LS** | dMRI QA End | CBL-Mot | 0.583894 | 0.000999 | 0.386146 | 240.3951 | 11.12833 | 0.035118 |
| **HCP-LS** | dMRI QA End | CBL-SAL | 0.528534 | 0.000999 | 0.286752 | 243.2999 | 11.20471 | 0.023459 |
| **HCP-LS** | dMRI QA End | CBL-SC | 0.602238 | 0.000999 | 0.363304 | 241.5802 | 11.16527 | 0.030361 |
| **HCP-LS** | dMRI QA End | CBL-VAs | 0.409981 | 0.000999 | 0.300801 | 246.8263 | 11.26839 | 0.009305 |
| **HCP-LS** | dMRI QA End | CBL-VII | 0.239433 | 0.000999 | 0.138561 | 247.9597 | 11.29515 | 0.004755 |
| **HCP-LS** | dMRI QA End | CBL-VI | 0.433142 | 0.000999 | 0.262736 | 246.3383 | 11.26533 | 0.011263 |
| **HCP-LS** | dMRI QA End | DMN-FP | 0.31321 | 0.000999 | 0.152279 | 247.1393 | 11.27835 | 0.008048 |
| **HCP-LS** | dMRI QA End | DMN-MF | 0.2542 | 0.000999 | 0.169571 | 247.8569 | 11.28976 | 0.005168 |
| **HCP-LS** | dMRI QA End | DMN-Mot | 0.321284 | 0.000999 | 0.2441 | 247.3924 | 11.28389 | 0.007032 |
| **HCP-LS** | dMRI QA End | DMN-SAL | 0.362956 | 0.000999 | 0.258098 | 246.9318 | 11.27354 | 0.008881 |
| **HCP-LS** | dMRI QA End | DMN-SC | 0.335127 | 0.000999 | 0.264634 | 247.3338 | 11.28197 | 0.007268 |
| **HCP-LS** | dMRI QA End | DMN-VAs | 0.092291 | 0.000999 | 0.066366 | 248.7532 | 11.3139 | 0.001571 |

| | | | | | | | | |
|---|---|---|---|---|---|---|---|---|
| **HCP-LS** | dMRI QA End | DMN-VII | 0.043412 | 0.000999 | 0.023475 | 248.9717 | 11.31595 | 0.000693 |
| **HCP-LS** | dMRI QA End | DMN-VI | 0.111734 | 0.000999 | 0.065104 | 248.6535 | 11.30919 | 0.001971 |
| **HCP-LS** | dMRI QA End | FP-MF | 0.550029 | 0.000999 | 0.346801 | 244.4119 | 11.219 | 0.018995 |
| **HCP-LS** | dMRI QA End | FP-Mot | 0.561008 | 0.000999 | 0.40242 | 243.0644 | 11.18583 | 0.024404 |
| **HCP-LS** | dMRI QA End | FP-SAL | 0.528943 | 0.000999 | 0.36324 | 242.8949 | 11.17866 | 0.025084 |
| **HCP-LS** | dMRI QA End | FP-SC | 0.560356 | 0.000999 | 0.364497 | 243.8631 | 11.20842 | 0.021198 |
| **HCP-LS** | dMRI QA End | FP-VAs | 0.381595 | 0.000999 | 0.27807 | 247.087 | 11.27859 | 0.008258 |
| **HCP-LS** | dMRI QA End | FP-VII | 0.221193 | 0.000999 | 0.150551 | 248.1061 | 11.30094 | 0.004168 |
| **HCP-LS** | dMRI QA End | FP-VI | 0.422635 | 0.000999 | 0.269366 | 246.0067 | 11.25191 | 0.012594 |
| **HCP-LS** | dMRI QA End | MF-Mot | 0.433867 | 0.000999 | 0.271372 | 246.1151 | 11.25586 | 0.012159 |
| **HCP-LS** | dMRI QA End | MF-SAL | 0.502164 | 0.000999 | 0.322297 | 245.4972 | 11.2452 | 0.014639 |
| **HCP-LS** | dMRI QA End | MF-SC | 0.44727 | 0.000999 | 0.315832 | 246.5064 | 11.26275 | 0.010588 |
| **HCP-LS** | dMRI QA End | MF-VAs | 0.213787 | 0.000999 | 0.144561 | 248.1486 | 11.30124 | 0.003997 |
| **HCP-LS** | dMRI QA End | MF-VII | 0.167312 | 0.000999 | 0.057915 | 248.3335 | 11.30492 | 0.003255 |
| **HCP-LS** | dMRI QA End | MF-VI | 0.360808 | 0.000999 | 0.210437 | 246.7757 | 11.26521 | 0.009508 |
| **HCP-LS** | dMRI QA End | Mot-SAL | 0.598753 | 0.000999 | 0.46496 | 243.1309 | 11.18995 | 0.024137 |
| **HCP-LS** | dMRI QA End | Mot-SC | 0.606188 | 0.000999 | 0.416245 | 241.7047 | 11.16454 | 0.029861 |
| **HCP-LS** | dMRI QA End | Mot-VAs | 0.45355 | 0.000999 | 0.357294 | 245.3438 | 11.23479 | 0.015255 |
| **HCP-LS** | dMRI QA End | Mot-VII | 0.209305 | 0.000999 | 0.128426 | 248.0898 | 11.29625 | 0.004233 |
| **HCP-LS** | dMRI QA End | Mot-VI | 0.369449 | 0.000999 | 0.240449 | 246.7575 | 11.26845 | 0.009581 |
| **HCP-LS** | dMRI QA End | SAL-SC | 0.550574 | 0.000999 | 0.390118 | 244.1159 | 11.21078 | 0.020183 |
| **HCP-LS** | dMRI QA End | SAL-VAs | 0.380054 | 0.000999 | 0.284879 | 247.1638 | 11.27412 | 0.00795 |
| **HCP-LS** | dMRI QA End | SAL-VII | 0.193833 | 0.000999 | 0.164861 | 248.2572 | 11.30165 | 0.003561 |

| | | | | | | | | |
|---|---|---|---|---|---|---|---|---|
| **HCP-LS** | dMRI QA End | SAL-VI | 0.415145 | 0.000999 | 0.272918 | 246.3657 | 11.25901 | 0.011153 |
| **HCP-LS** | dMRI QA End | SC-VAs | 0.319794 | 0.000999 | 0.213892 | 247.557 | 11.28412 | 0.006372 |
| **HCP-LS** | dMRI QA End | SC-VII | 0.167915 | 0.000999 | 0.123043 | 248.3767 | 11.30492 | 0.003082 |
| **HCP-LS** | dMRI QA End | SC-VI | 0.345248 | 0.000999 | 0.244981 | 247.1717 | 11.2739 | 0.007918 |
| **HCP-LS** | dMRI QA End | VAs-VII | 0.09779 | 0.000999 | 0.077566 | 248.7206 | 11.30834 | 0.001702 |
| **HCP-LS** | dMRI QA End | VAs-VI | 0.164203 | 0.000999 | 0.130243 | 248.4195 | 11.30673 | 0.00291 |
| **HCP-LS** | dMRI QA End | VI-VII | 0.061246 | 0.000999 | -0.00125 | 248.8853 | 11.31675 | 0.00104 |
| **HCP-D** | rsfMRI | CBL-DMN | 0.660898 | 0.000999 | 0.68906 | 12.62019 | 2.964904 | 0.110806 |
| **HCP-D** | rsfMRI | CBL-FP | 0.700008 | 0.000999 | 0.717807 | 11.80405 | 2.873244 | 0.16831 |
| **HCP-D** | rsfMRI | CBL-MF | 0.659054 | 0.000999 | 0.673862 | 12.12651 | 2.902308 | 0.145589 |
| **HCP-D** | rsfMRI | CBL-Mot | 0.674803 | 0.000999 | 0.688187 | 10.99579 | 2.761061 | 0.225258 |
| **HCP-D** | rsfMRI | CBL-SAL | 0.655567 | 0.000999 | 0.683629 | 12.04944 | 2.895352 | 0.15102 |
| **HCP-D** | rsfMRI | CBL-SC | 0.702511 | 0.000999 | 0.72305 | 12.30701 | 2.929346 | 0.132872 |
| **HCP-D** | rsfMRI | CBL-VAs | 0.638415 | 0.000999 | 0.648053 | 12.43867 | 2.943249 | 0.123595 |
| **HCP-D** | rsfMRI | CBL-VII | 0.591118 | 0.000999 | 0.600622 | 13.52565 | 3.075134 | 0.047009 |
| **HCP-D** | rsfMRI | CBL-VI | 0.619497 | 0.000999 | 0.632231 | 12.9866 | 3.00661 | 0.084989 |
| **HCP-D** | rsfMRI | DMN-FP | 0.608227 | 0.000999 | 0.617189 | 13.04687 | 3.017929 | 0.080743 |
| **HCP-D** | rsfMRI | DMN-MF | 0.558315 | 0.000999 | 0.571473 | 13.28088 | 3.040002 | 0.064255 |
| **HCP-D** | rsfMRI | DMN-Mot | 0.607957 | 0.000999 | 0.620654 | 12.45222 | 2.946526 | 0.12264 |
| **HCP-D** | rsfMRI | DMN-SAL | 0.649337 | 0.000999 | 0.676404 | 12.71289 | 2.97533 | 0.104275 |
| **HCP-D** | rsfMRI | DMN-SC | 0.663845 | 0.000999 | 0.681027 | 13.19767 | 3.03554 | 0.070118 |
| **HCP-D** | rsfMRI | DMN-VAs | 0.575269 | 0.000999 | 0.589151 | 13.21543 | 3.042763 | 0.068866 |
| **HCP-D** | rsfMRI | DMN-VII | 0.440576 | 0.000999 | 0.446553 | 13.82977 | 3.110035 | 0.025581 |

| | | | | | | | | |
|---|---|---|---|---|---|---|---|---|
| **HCP-D** | rsfMRI | DMN-VI | 0.513192 | 0.000999 | 0.524141 | 13.74997 | 3.097863 | 0.031204 |
| **HCP-D** | rsfMRI | FP-MF | 0.670357 | 0.000999 | 0.673801 | 12.67401 | 2.976097 | 0.107014 |
| **HCP-D** | rsfMRI | FP-Mot | 0.704194 | 0.000999 | 0.708418 | 11.51242 | 2.836528 | 0.188857 |
| **HCP-D** | rsfMRI | FP-SAL | 0.70848 | 0.000999 | 0.730092 | 11.9278 | 2.887204 | 0.15959 |
| **HCP-D** | rsfMRI | FP-SC | 0.730487 | 0.000999 | 0.745911 | 12.26429 | 2.925939 | 0.135882 |
| **HCP-D** | rsfMRI | FP-VAs | 0.694088 | 0.000999 | 0.702335 | 12.81275 | 2.992777 | 0.097238 |
| **HCP-D** | rsfMRI | FP-VII | 0.605578 | 0.000999 | 0.611631 | 13.56018 | 3.082439 | 0.044576 |
| **HCP-D** | rsfMRI | FP-VI | 0.629772 | 0.000999 | 0.641016 | 13.25428 | 3.042728 | 0.066129 |
| **HCP-D** | rsfMRI | MF-Mot | 0.645326 | 0.000999 | 0.652396 | 12.09286 | 2.903991 | 0.14796 |
| **HCP-D** | rsfMRI | MF-SAL | 0.661656 | 0.000999 | 0.68418 | 12.28899 | 2.921722 | 0.134142 |
| **HCP-D** | rsfMRI | MF-SC | 0.705631 | 0.000999 | 0.71844 | 12.585 | 2.965903 | 0.113285 |
| **HCP-D** | rsfMRI | MF-VAs | 0.618856 | 0.000999 | 0.628846 | 13.06405 | 3.015286 | 0.079533 |
| **HCP-D** | rsfMRI | MF-VII | 0.504226 | 0.000999 | 0.508554 | 13.68011 | 3.090471 | 0.036126 |
| **HCP-D** | rsfMRI | MF-VI | 0.552667 | 0.000999 | 0.561228 | 13.43994 | 3.061305 | 0.053048 |
| **HCP-D** | rsfMRI | Mot-SAL | 0.673453 | 0.000999 | 0.694198 | 11.25577 | 2.797854 | 0.20694 |
| **HCP-D** | rsfMRI | Mot-SC | 0.721965 | 0.000999 | 0.728979 | 11.66677 | 2.859407 | 0.177982 |
| **HCP-D** | rsfMRI | Mot-VAs | 0.601488 | 0.000999 | 0.603671 | 12.29991 | 2.935068 | 0.133372 |
| **HCP-D** | rsfMRI | Mot-VII | 0.560843 | 0.000999 | 0.562366 | 13.3005 | 3.051404 | 0.062873 |
| **HCP-D** | rsfMRI | Mot-VI | 0.611463 | 0.000999 | 0.627892 | 12.682 | 2.975179 | 0.106451 |
| **HCP-D** | rsfMRI | SAL-SC | 0.696721 | 0.000999 | 0.716252 | 12.3457 | 2.929884 | 0.130146 |
| **HCP-D** | rsfMRI | SAL-VAs | 0.622776 | 0.000999 | 0.641616 | 12.60105 | 2.964401 | 0.112155 |
| **HCP-D** | rsfMRI | SAL-VII | 0.5765 | 0.000999 | 0.579182 | 13.55563 | 3.077003 | 0.044896 |
| **HCP-D** | rsfMRI | SAL-VI | 0.605784 | 0.000999 | 0.61211 | 13.18882 | 3.033193 | 0.070741 |

| | | | | | | | | |
|---|---|---|---|---|---|---|---|---|
| **HCP-D** | rsfMRI | SC-VAs | 0.69385 | 0.000999 | 0.70463 | 12.81239 | 2.991566 | 0.097264 |
| **HCP-D** | rsfMRI | SC-VII | 0.573645 | 0.000999 | 0.565228 | 13.65491 | 3.09355 | 0.037901 |
| **HCP-D** | rsfMRI | SC-VI | 0.633224 | 0.000999 | 0.638402 | 13.31933 | 3.049672 | 0.061546 |
| **HCP-D** | rsfMRI | VAs-VII | 0.440506 | 0.000999 | 0.43453 | 13.87359 | 3.117592 | 0.022494 |
| **HCP-D** | rsfMRI | VAs-VI | 0.499816 | 0.000999 | 0.504311 | 13.67413 | 3.092851 | 0.036547 |
| **HCP-D** | rsfMRI | VI-VII | 0.327931 | 0.000999 | 0.341753 | 13.99773 | 3.128682 | 0.013747 |
| **HCP-YA** | rsfMRI | CBL-DMN | 0.132134 | 0.001215 | 0.142168 | 13.48041 | 3.085586 | 0.00735 |
| **HCP-YA** | rsfMRI | CBL-FP | 0.156648 | 0.001215 | 0.157888 | 13.4176 | 3.07769 | 0.011975 |
| **HCP-YA** | rsfMRI | CBL-MF | 0.189562 | 0.001215 | 0.191679 | 13.33382 | 3.061242 | 0.018144 |
| **HCP-YA** | rsfMRI | CBL-Mot | 0.164844 | 0.001215 | 0.178481 | 13.29246 | 3.04757 | 0.02119 |
| **HCP-YA** | rsfMRI | CBL-SAL | 0.148161 | 0.001215 | 0.159234 | 13.39615 | 3.072272 | 0.013554 |
| **HCP-YA** | rsfMRI | CBL-SC | 0.162057 | 0.001215 | 0.16545 | 13.4323 | 3.077674 | 0.010893 |
| **HCP-YA** | rsfMRI | CBL-VAs | 0.138279 | 0.001215 | 0.152799 | 13.41091 | 3.071674 | 0.012467 |
| **HCP-YA** | rsfMRI | CBL-VII | 0.079569 | 0.001215 | 0.095489 | 13.53505 | 3.092029 | 0.003327 |
| **HCP-YA** | rsfMRI | CBL-VI | 0.130664 | 0.001215 | 0.143107 | 13.46442 | 3.082554 | 0.008527 |
| **HCP-YA** | rsfMRI | DMN-FP | 0.181708 | 0.001215 | 0.186494 | 13.42532 | 3.082119 | 0.011407 |
| **HCP-YA** | rsfMRI | DMN-MF | 0.18823 | 0.001215 | 0.196173 | 13.43735 | 3.079238 | 0.010521 |
| **HCP-YA** | rsfMRI | DMN-Mot | 0.166098 | 0.001215 | 0.179637 | 13.38479 | 3.070746 | 0.014391 |
| **HCP-YA** | rsfMRI | DMN-SAL | 0.14814 | 0.001215 | 0.153873 | 13.46262 | 3.085845 | 0.00866 |
| **HCP-YA** | rsfMRI | DMN-SC | 0.185253 | 0.001215 | 0.184629 | 13.43993 | 3.084623 | 0.01033 |
| **HCP-YA** | rsfMRI | DMN-VAs | 0.136956 | 0.001215 | 0.140716 | 13.50036 | 3.090984 | 0.005881 |
| **HCP-YA** | rsfMRI | DMN-VII | -0.03309 | 0.112887 | -0.03358 | 13.59978 | 3.105332 | -0.00144 |
| **HCP-YA** | rsfMRI | DMN-VI | 0.109698 | 0.001215 | 0.111319 | 13.51914 | 3.092376 | 0.004498 |

| | | | | | | | | |
|---|---|---|---|---|---|---|---|---|
| **HCP-YA** | rsfMRI | FP-MF | 0.161528 | 0.001215 | 0.183761 | 13.41756 | 3.078931 | 0.011978 |
| **HCP-YA** | rsfMRI | FP-Mot | 0.192063 | 0.001215 | 0.19635 | 13.31208 | 3.060315 | 0.019745 |
| **HCP-YA** | rsfMRI | FP-SAL | 0.184812 | 0.001215 | 0.189732 | 13.405 | 3.07708 | 0.012902 |
| **HCP-YA** | rsfMRI | FP-SC | 0.158049 | 0.001215 | 0.162391 | 13.4522 | 3.087379 | 0.009427 |
| **HCP-YA** | rsfMRI | FP-VAs | 0.093662 | 0.003289 | 0.106459 | 13.51573 | 3.090805 | 0.004749 |
| **HCP-YA** | rsfMRI | FP-VII | 0.08474 | 0.005352 | 0.073176 | 13.54209 | 3.096135 | 0.002808 |
| **HCP-YA** | rsfMRI | FP-VI | 0.163039 | 0.001215 | 0.175203 | 13.44372 | 3.082945 | 0.010051 |
| **HCP-YA** | rsfMRI | MF-Mot | 0.199257 | 0.001215 | 0.204564 | 13.28513 | 3.051549 | 0.02173 |
| **HCP-YA** | rsfMRI | MF-SAL | 0.149679 | 0.001215 | 0.157987 | 13.42546 | 3.077765 | 0.011396 |
| **HCP-YA** | rsfMRI | MF-SC | 0.169702 | 0.001215 | 0.191048 | 13.41978 | 3.076639 | 0.011814 |
| **HCP-YA** | rsfMRI | MF-VAs | 0.14847 | 0.001215 | 0.152739 | 13.43922 | 3.078863 | 0.010383 |
| **HCP-YA** | rsfMRI | MF-VII | 0.10828 | 0.001215 | 0.10417 | 13.52593 | 3.095924 | 0.003998 |
| **HCP-YA** | rsfMRI | MF-VI | 0.153885 | 0.001215 | 0.162405 | 13.43649 | 3.079422 | 0.010584 |
| **HCP-YA** | rsfMRI | Mot-SAL | 0.146573 | 0.001215 | 0.176993 | 13.36689 | 3.065723 | 0.015709 |
| **HCP-YA** | rsfMRI | Mot-SC | 0.192881 | 0.001215 | 0.203201 | 13.28166 | 3.055393 | 0.021985 |
| **HCP-YA** | rsfMRI | Mot-VAs | 0.142326 | 0.002248 | 0.155962 | 13.37084 | 3.059693 | 0.015418 |
| **HCP-YA** | rsfMRI | Mot-VII | 0.116281 | 0.001215 | 0.127272 | 13.48937 | 3.089622 | 0.00669 |
| **HCP-YA** | rsfMRI | Mot-VI | 0.155429 | 0.001215 | 0.17527 | 13.39754 | 3.070867 | 0.013452 |
| **HCP-YA** | rsfMRI | SAL-SC | 0.135021 | 0.001215 | 0.141555 | 13.46504 | 3.085894 | 0.008481 |
| **HCP-YA** | rsfMRI | SAL-VAs | 0.105559 | 0.001215 | 0.114858 | 13.49894 | 3.088321 | 0.005985 |
| **HCP-YA** | rsfMRI | SAL-VII | 0.079176 | 0.002248 | 0.072839 | 13.53694 | 3.095568 | 0.003187 |
| **HCP-YA** | rsfMRI | SAL-VI | 0.13382 | 0.001215 | 0.134554 | 13.47225 | 3.089039 | 0.00795 |
| **HCP-YA** | rsfMRI | SC-VAs | 0.112157 | 0.001215 | 0.105879 | 13.49969 | 3.087795 | 0.00593 |

| | | | | | | | | |
|---|---|---|---|---|---|---|---|---|
| **HCP-YA** | rsfMRI | SC-VII | 0.025895 | 0.014637 | 0.021911 | 13.57168 | 3.100572 | 0.000629 |
| **HCP-YA** | rsfMRI | SC-VI | 0.109359 | 0.002248 | 0.131785 | 13.49975 | 3.090117 | 0.005926 |
| **HCP-YA** | rsfMRI | VAs-VII | 0.014116 | 0.029629 | 0.026166 | 13.57804 | 3.100221 | 0.000161 |
| **HCP-YA** | rsfMRI | VAs-VI | 0.09093 | 0.001215 | 0.108748 | 13.51687 | 3.091446 | 0.004665 |
| **HCP-YA** | rsfMRI | VI-VII | 0.088555 | 0.001215 | 0.094167 | 13.53926 | 3.095606 | 0.003017 |
| **HCP-A** | rsfMRI | CBL-DMN | -0.01539 | 0.003458 | -0.01605 | 176.2532 | 10.99115 | -0.00109 |
| **HCP-A** | rsfMRI | CBL-FP | 0.081191 | 0.002141 | 0.087202 | 175.525 | 10.95901 | 0.003051 |
| **HCP-A** | rsfMRI | CBL-MF | 0.08547 | 0.002141 | 0.060969 | 175.4736 | 10.95992 | 0.003342 |
| **HCP-A** | rsfMRI | CBL-Mot | 0.168699 | 0.002141 | 0.150989 | 174.6891 | 10.92412 | 0.007798 |
| **HCP-A** | rsfMRI | CBL-SAL | 0.082285 | 0.002141 | 0.051682 | 175.5212 | 10.95391 | 0.003072 |
| **HCP-A** | rsfMRI | CBL-SC | -0.00257 | 0.002141 | 0.011071 | 176.1564 | 10.96335 | -0.00054 |
| **HCP-A** | rsfMRI | CBL-VAs | 0.06323 | 0.002141 | 0.075721 | 175.6853 | 10.9709 | 0.00214 |
| **HCP-A** | rsfMRI | CBL-VII | -0.10894 | 0.075924 | -0.09226 | 176.8978 | 10.99798 | -0.00475 |
| **HCP-A** | rsfMRI | CBL-VI | -0.04536 | 0.018225 | -0.06786 | 176.4606 | 11.00321 | -0.00226 |
| **HCP-A** | rsfMRI | DMN-FP | -0.01624 | 0.008701 | -0.0359 | 176.2627 | 10.98946 | -0.00114 |
| **HCP-A** | rsfMRI | DMN-MF | 0.042117 | 0.002141 | 0.044066 | 175.8254 | 10.95011 | 0.001345 |
| **HCP-A** | rsfMRI | DMN-Mot | 0.124902 | 0.002141 | 0.116686 | 175.0819 | 10.92646 | 0.005567 |
| **HCP-A** | rsfMRI | DMN-SAL | -0.04644 | 0.014129 | -0.029 | 176.4813 | 10.99599 | -0.00238 |
| **HCP-A** | rsfMRI | DMN-SC | -0.03101 | 0.011239 | -0.02365 | 176.3746 | 10.97957 | -0.00177 |
| **HCP-A** | rsfMRI | DMN-VAs | -0.04156 | 0.002141 | -0.03742 | 176.4426 | 10.98338 | -0.00216 |
| **HCP-A** | rsfMRI | DMN-VII | -0.0943 | 0.047148 | -0.09008 | 176.7982 | 10.99402 | -0.00418 |
| **HCP-A** | rsfMRI | DMN-VI | -0.06419 | 0.020111 | -0.02568 | 176.5984 | 11.0018 | -0.00305 |
| **HCP-A** | rsfMRI | FP-MF | 0.067966 | 0.003458 | 0.059247 | 175.6028 | 10.95836 | 0.002609 |

| | | | | | | | | |
|---|---|---|---|---|---|---|---|---|
| **HCP-A** | rsfMRI | FP-Mot | 0.227119 | 0.002141 | 0.196515 | 173.7625 | 10.87408 | 0.013061 |
| **HCP-A** | rsfMRI | FP-SAL | 0.068211 | 0.002141 | 0.074112 | 175.602 | 10.95108 | 0.002613 |
| **HCP-A** | rsfMRI | FP-SC | -0.00366 | 0.007751 | -0.01764 | 176.1691 | 10.98952 | -0.00061 |
| **HCP-A** | rsfMRI | FP-VAs | 0.057446 | 0.002141 | 0.045502 | 175.7025 | 10.95701 | 0.002042 |
| **HCP-A** | rsfMRI | FP-VII | -0.05822 | 0.008701 | -0.02442 | 176.53 | 10.99186 | -0.00266 |
| **HCP-A** | rsfMRI | FP-VI | -0.04709 | 0.014129 | -0.04291 | 176.4748 | 10.99734 | -0.00234 |
| **HCP-A** | rsfMRI | MF-Mot | 0.209685 | 0.002141 | 0.195791 | 173.9201 | 10.90187 | 0.012166 |
| **HCP-A** | rsfMRI | MF-SAL | 0.015762 | 0.004995 | 0.017436 | 176.0256 | 10.96521 | 0.000207 |
| **HCP-A** | rsfMRI | MF-SC | 0.036168 | 0.002141 | 0.024016 | 175.8608 | 10.96398 | 0.001143 |
| **HCP-A** | rsfMRI | MF-VAs | 0.093237 | 0.002141 | 0.092922 | 175.4297 | 10.93536 | 0.003592 |
| **HCP-A** | rsfMRI | MF-VII | -0.09787 | 0.047148 | -0.08426 | 176.8031 | 11.00746 | -0.00421 |
| **HCP-A** | rsfMRI | MF-VI | -0.02432 | 0.006422 | -0.01306 | 176.315 | 10.96236 | -0.00144 |
| **HCP-A** | rsfMRI | Mot-SAL | 0.265899 | 0.002141 | 0.255698 | 173.698 | 10.89593 | 0.013428 |
| **HCP-A** | rsfMRI | Mot-SC | 0.159068 | 0.002141 | 0.157785 | 174.6861 | 10.92432 | 0.007815 |
| **HCP-A** | rsfMRI | Mot-VAs | 0.158664 | 0.002141 | 0.131256 | 174.759 | 10.92969 | 0.007401 |
| **HCP-A** | rsfMRI | Mot-VII | -0.05082 | 0.014985 | -0.04178 | 176.4881 | 10.9938 | -0.00242 |
| **HCP-A** | rsfMRI | Mot-VI | 0.105216 | 0.002141 | 0.078642 | 175.2859 | 10.94309 | 0.004409 |
| **HCP-A** | rsfMRI | SAL-SC | 0.00652 | 0.003458 | -0.0174 | 176.0883 | 10.98828 | -0.00015 |
| **HCP-A** | rsfMRI | SAL-VAs | 0.068413 | 0.002141 | 0.053609 | 175.6371 | 10.96411 | 0.002414 |
| **HCP-A** | rsfMRI | SAL-VII | -0.10736 | 0.055172 | -0.10466 | 176.8826 | 10.99711 | -0.00466 |
| **HCP-A** | rsfMRI | SAL-VI | -0.03227 | 0.013623 | -0.01326 | 176.3731 | 10.98326 | -0.00177 |
| **HCP-A** | rsfMRI | SC-VAs | 0.007467 | 0.003458 | 0.028427 | 176.0858 | 10.97133 | -0.00013 |
| **HCP-A** | rsfMRI | SC-VII | -0.08786 | 0.031123 | -0.04872 | 176.7203 | 11.00419 | -0.00374 |

| | | | | | | | | |
|---|---|---|---|---|---|---|---|---|
| **HCP-A** | rsfMRI | SC-VI | -0.0033 | 0.002141 | -0.00718 | 176.159 | 10.98514 | -0.00055 |
| **HCP-A** | rsfMRI | VAs-VII | -0.09901 | 0.053319 | -0.06476 | 176.8084 | 10.99936 | -0.00424 |
| **HCP-A** | rsfMRI | VAs-VI | -0.01398 | 0.003458 | -0.0261 | 176.2376 | 10.97412 | -0.001 |
| **HCP-A** | rsfMRI | VI-VII | -0.10358 | 0.053319 | -0.06259 | 176.8356 | 11.01357 | -0.00439 |
| **HCP-LS** | rsfMRI | CBL-DMN | 0.591543 | 0.000999 | 0.563819 | 243.0062 | 11.18319 | 0.024638 |
| **HCP-LS** | rsfMRI | CBL-FP | 0.691496 | 0.000999 | 0.717659 | 237.5028 | 11.0465 | 0.046727 |
| **HCP-LS** | rsfMRI | CBL-MF | 0.649848 | 0.000999 | 0.651456 | 240.0783 | 11.11407 | 0.036389 |
| **HCP-LS** | rsfMRI | CBL-Mot | 0.676776 | 0.000999 | 0.664892 | 238.1181 | 11.07336 | 0.044257 |
| **HCP-LS** | rsfMRI | CBL-SAL | 0.6207 | 0.000999 | 0.624234 | 241.2436 | 11.14173 | 0.031712 |
| **HCP-LS** | rsfMRI | CBL-SC | 0.618455 | 0.000999 | 0.664566 | 241.2096 | 11.13524 | 0.031848 |
| **HCP-LS** | rsfMRI | CBL-VAs | 0.524699 | 0.000999 | 0.4973 | 245.4314 | 11.24236 | 0.014903 |
| **HCP-LS** | rsfMRI | CBL-VII | 0.412265 | 0.000999 | 0.390428 | 246.9124 | 11.26712 | 0.008959 |
| **HCP-LS** | rsfMRI | CBL-VI | 0.589038 | 0.000999 | 0.550517 | 243.7634 | 11.19753 | 0.021598 |
| **HCP-LS** | rsfMRI | DMN-FP | 0.682277 | 0.000999 | 0.669415 | 242.134 | 11.15297 | 0.028138 |
| **HCP-LS** | rsfMRI | DMN-MF | 0.583322 | 0.000999 | 0.553809 | 243.5377 | 11.19816 | 0.022504 |
| **HCP-LS** | rsfMRI | DMN-Mot | 0.591342 | 0.000999 | 0.568433 | 241.9662 | 11.16563 | 0.028812 |
| **HCP-LS** | rsfMRI | DMN-SAL | 0.574384 | 0.000999 | 0.51626 | 243.503 | 11.19722 | 0.022643 |
| **HCP-LS** | rsfMRI | DMN-SC | 0.555497 | 0.000999 | 0.539866 | 244.9703 | 11.22482 | 0.016754 |
| **HCP-LS** | rsfMRI | DMN-VAs | 0.500062 | 0.000999 | 0.474339 | 245.5888 | 11.24219 | 0.014272 |
| **HCP-LS** | rsfMRI | DMN-VII | 0.325535 | 0.000999 | 0.337415 | 247.4125 | 11.27909 | 0.006952 |
| **HCP-LS** | rsfMRI | DMN-VI | 0.477114 | 0.000999 | 0.419463 | 246.0479 | 11.25125 | 0.012429 |
| **HCP-LS** | rsfMRI | FP-MF | 0.714809 | 0.000999 | 0.683068 | 237.9516 | 11.05927 | 0.044925 |
| **HCP-LS** | rsfMRI | FP-Mot | 0.708248 | 0.000999 | 0.679428 | 236.017 | 11.01512 | 0.05269 |

| | | | | | | | | |
|---|---|---|---|---|---|---|---|---|
| **HCP-LS** | rsfMRI | FP-SAL | 0.681811 | 0.000999 | 0.657819 | 238.2631 | 11.07607 | 0.043675 |
| **HCP-LS** | rsfMRI | FP-SC | 0.703681 | 0.000999 | 0.712395 | 239.8427 | 11.10528 | 0.037335 |
| **HCP-LS** | rsfMRI | FP-VAs | 0.612248 | 0.000999 | 0.575503 | 243.3525 | 11.19158 | 0.023247 |
| **HCP-LS** | rsfMRI | FP-VII | 0.457303 | 0.000999 | 0.431724 | 246.2023 | 11.25555 | 0.011809 |
| **HCP-LS** | rsfMRI | FP-VI | 0.626139 | 0.000999 | 0.587626 | 243.4454 | 11.18429 | 0.022875 |
| **HCP-LS** | rsfMRI | MF-Mot | 0.626501 | 0.000999 | 0.601371 | 238.017 | 11.07758 | 0.044663 |
| **HCP-LS** | rsfMRI | MF-SAL | 0.62345 | 0.000999 | 0.579661 | 240.5058 | 11.13039 | 0.034673 |
| **HCP-LS** | rsfMRI | MF-SC | 0.627068 | 0.000999 | 0.620632 | 242.4038 | 11.17221 | 0.027055 |
| **HCP-LS** | rsfMRI | MF-VAs | 0.523251 | 0.000999 | 0.52172 | 244.6728 | 11.22144 | 0.017948 |
| **HCP-LS** | rsfMRI | MF-VII | 0.375179 | 0.000999 | 0.351627 | 246.9415 | 11.27113 | 0.008842 |
| **HCP-LS** | rsfMRI | MF-VI | 0.549266 | 0.000999 | 0.442075 | 244.6431 | 11.2219 | 0.018068 |
| **HCP-LS** | rsfMRI | Mot-SAL | 0.644995 | 0.000999 | 0.607295 | 235.5608 | 11.02684 | 0.054521 |
| **HCP-LS** | rsfMRI | Mot-SC | 0.641594 | 0.000999 | 0.556532 | 240.0445 | 11.12752 | 0.036525 |
| **HCP-LS** | rsfMRI | Mot-VAs | 0.578843 | 0.000999 | 0.455053 | 243.314 | 11.19881 | 0.023402 |
| **HCP-LS** | rsfMRI | Mot-VII | 0.499539 | 0.000999 | 0.34162 | 244.9967 | 11.23283 | 0.016648 |
| **HCP-LS** | rsfMRI | Mot-VI | 0.56363 | 0.000999 | 0.432725 | 242.5373 | 11.1779 | 0.026519 |
| **HCP-LS** | rsfMRI | SAL-SC | 0.611286 | 0.000999 | 0.612191 | 241.5371 | 11.1495 | 0.030534 |
| **HCP-LS** | rsfMRI | SAL-VAs | 0.519939 | 0.000999 | 0.517944 | 244.1216 | 11.20602 | 0.02016 |
| **HCP-LS** | rsfMRI | SAL-VII | 0.419761 | 0.000999 | 0.387924 | 246.7044 | 11.26809 | 0.009794 |
| **HCP-LS** | rsfMRI | SAL-VI | 0.580071 | 0.000999 | 0.466627 | 243.5619 | 11.20363 | 0.022407 |
| **HCP-LS** | rsfMRI | SC-VAs | 0.427661 | 0.000999 | 0.376166 | 246.5874 | 11.26611 | 0.010263 |
| **HCP-LS** | rsfMRI | SC-VII | 0.346481 | 0.000999 | 0.342858 | 247.3526 | 11.28024 | 0.007192 |
| **HCP-LS** | rsfMRI | SC-VI | 0.566397 | 0.000999 | 0.477865 | 244.5153 | 11.21703 | 0.01858 |

| | | | | | | | | |
|---|---|---|---|---|---|---|---|---|
| **HCP-LS** | rsfMRI | VAs-VII | 0.23546 | 0.000999 | 0.241998 | 248.0554 | 11.29579 | 0.004371 |
| **HCP-LS** | rsfMRI | VAs-VI | 0.400318 | 0.000999 | 0.372052 | 246.7394 | 11.26649 | 0.009654 |
| **HCP-LS** | rsfMRI | VI-VII | 0.240903 | 0.000999 | 0.255372 | 248.0003 | 11.29379 | 0.004593 |
| **Within-Network** | | | | | | | | |
| **Cohort** | **Modality** | **Network** | **Pearson r** | **pFDR** | **Spearman rho** | **MSE** | **MAE** | **q2** |
| **HCP-D** | dMRI QA End | CBL | 0.232719 | 0.00111 | 0.253628 | 13.99821 | 3.123869 | 0.013713 |
| **HCP-D** | dMRI QA End | DMN | 0.233067 | 0.00111 | 0.206389 | 14.05392 | 3.133684 | 0.009788 |
| **HCP-D** | dMRI QA End | FP | 0.340549 | 0.00111 | 0.347884 | 13.90747 | 3.120441 | 0.020107 |
| **HCP-D** | dMRI QA End | MF | 0.127096 | 0.00111 | 0.110475 | 14.13322 | 3.144121 | 0.004201 |
| **HCP-D** | dMRI QA End | Mot | 0.290283 | 0.00111 | 0.292491 | 13.62114 | 3.085957 | 0.040281 |
| **HCP-D** | dMRI QA End | SAL | 0.300447 | 0.00111 | 0.313817 | 13.9885 | 3.127139 | 0.014397 |
| **HCP-D** | dMRI QA End | SC | 0.296918 | 0.00111 | 0.295174 | 13.9995 | 3.129361 | 0.013622 |
| **HCP-D** | dMRI QA End | VAs | 0.125937 | 0.00111 | 0.127984 | 14.12553 | 3.145049 | 0.004743 |
| **HCP-D** | dMRI QA End | VII | -0.1271 | 0.444328 | -0.12759 | 14.25412 | 3.159411 | -0.00432 |
| **HCP-D** | dMRI QA End | VI | 0.017643 | 0.00111 | 0.026203 | 14.18843 | 3.150094 | 0.00031 |
| **HCP-YA** | dMRI QA End | CBL | 0.008732 | 0.127373 | 0.011732 | 13.58179 | 3.102398 | -0.00012 |
| **HCP-YA** | dMRI QA End | DMN | -0.08007 | 0.385864 | -0.09336 | 13.61577 | 3.107438 | -0.00262 |
| **HCP-YA** | dMRI QA End | FP | 0.008736 | 0.127373 | -0.01 | 13.58028 | 3.102274 | -4.24E-06 |
| **HCP-YA** | dMRI QA End | MF | -0.03077 | 0.14652 | -0.03258 | 13.59743 | 3.104327 | -0.00127 |
| **HCP-YA** | dMRI QA End | Mot | 0.050149 | 0.10989 | 0.046611 | 13.55577 | 3.10035 | 0.0018 |
| **HCP-YA** | dMRI QA End | SAL | -0.04481 | 0.316826 | -0.04813 | 13.60639 | 3.1062 | -0.00193 |
| **HCP-YA** | dMRI QA End | SC | 0.02251 | 0.10989 | 0.019143 | 13.57361 | 3.101052 | 0.000487 |
| **HCP-YA** | dMRI QA End | VAs | -0.04552 | 0.14652 | -0.05233 | 13.60221 | 3.106964 | -0.00162 |
| **HCP-YA** | dMRI QA End | VII | -0.11465 | 0.547988 | -0.11037 | 13.45551 | 3.091926 | -0.00256 |

| | | | | | | | | |
|---|---|---|---|---|---|---|---|---|
| **HCP-YA** | dMRI QA End | VI | -0.10204 | 0.547988 | -0.10046 | 13.62138 | 3.107036 | -0.00303 |
| **HCP-A** | dMRI QA End | CBL | 0.208699 | 0.001998 | 0.177175 | 174.489 | 10.9375 | 0.008935 |
| **HCP-A** | dMRI QA End | DMN | -0.12832 | 0.183566 | -0.09731 | 176.9915 | 11.00656 | -0.00528 |
| **HCP-A** | dMRI QA End | FP | 0.068805 | 0.001998 | 0.086004 | 175.6497 | 10.96508 | 0.002342 |
| **HCP-A** | dMRI QA End | MF | -0.1024 | 0.072784 | -0.05683 | 176.8674 | 10.99635 | -0.00457 |
| **HCP-A** | dMRI QA End | Mot | 0.235582 | 0.001998 | 0.201049 | 173.3433 | 10.89235 | 0.015442 |
| **HCP-A** | dMRI QA End | SAL | 0.158219 | 0.001998 | 0.197228 | 174.9332 | 10.92991 | 0.006412 |
| **HCP-A** | dMRI QA End | SC | 0.03909 | 0.001998 | 0.078513 | 175.8567 | 10.97472 | 0.001166 |
| **HCP-A** | dMRI QA End | VAs | -0.05882 | 0.00333 | -0.0166 | 176.5411 | 10.99539 | -0.00272 |
| **HCP-A** | dMRI QA End | VII | -0.15568 | 0.418605 | -0.09543 | 177.1814 | 11.00101 | -0.00636 |
| **HCP-A** | dMRI QA End | VI | -0.15405 | 0.3774 | -0.13227 | 177.1808 | 11.01466 | -0.00635 |
| **HCP-LS** | dMRI QA End | CBL | 0.463544 | 0.00111 | 0.309033 | 245.159 | 11.23218 | 0.015997 |
| **HCP-LS** | dMRI QA End | DMN | 0.222334 | 0.00111 | 0.110188 | 247.9463 | 11.29678 | 0.004809 |
| **HCP-LS** | dMRI QA End | FP | 0.472722 | 0.00111 | 0.358774 | 245.5356 | 11.23961 | 0.014485 |
| **HCP-LS** | dMRI QA End | MF | 0.288714 | 0.00111 | 0.155728 | 247.622 | 11.28726 | 0.006111 |
| **HCP-LS** | dMRI QA End | Mot | 0.440046 | 0.00111 | 0.306955 | 243.4702 | 11.18332 | 0.022775 |
| **HCP-LS** | dMRI QA End | SAL | 0.45661 | 0.00111 | 0.327101 | 246.0635 | 11.25406 | 0.012366 |
| **HCP-LS** | dMRI QA End | SC | 0.442587 | 0.00111 | 0.309502 | 245.9905 | 11.25143 | 0.012659 |
| **HCP-LS** | dMRI QA End | VAs | 0.111012 | 0.00111 | 0.102804 | 248.6657 | 11.30767 | 0.001922 |
| **HCP-LS** | dMRI QA End | VII | -0.04424 | 0.078332 | -0.00048 | 249.3501 | 11.32776 | -0.00083 |
| **HCP-LS** | dMRI QA End | VI | 0.124632 | 0.00111 | 0.049869 | 248.5765 | 11.30934 | 0.00228 |
| **HCP-D** | rsfMRI | CBL | 0.631644 | 0.00101 | 0.641235 | 12.65751 | 2.974151 | 0.108177 |
| **HCP-D** | rsfMRI | DMN | 0.397086 | 0.00101 | 0.406444 | 13.79509 | 3.102982 | 0.028025 |

| | | | | | | | | |
|---|---|---|---|---|---|---|---|---|
| **HCP-D** | rsfMRI | FP | 0.671287 | 0.00101 | 0.674587 | 12.92079 | 3.009813 | 0.089626 |
| **HCP-D** | rsfMRI | MF | 0.560368 | 0.00101 | 0.56372 | 13.39422 | 3.05916 | 0.056269 |
| **HCP-D** | rsfMRI | Mot | 0.6198 | 0.00101 | 0.625793 | 11.95627 | 2.892453 | 0.157584 |
| **HCP-D** | rsfMRI | SAL | 0.593221 | 0.00101 | 0.63172 | 13.00852 | 3.007397 | 0.083445 |
| **HCP-D** | rsfMRI | SC | 0.649913 | 0.00101 | 0.655887 | 13.30878 | 3.05372 | 0.062289 |
| **HCP-D** | rsfMRI | VAs | 0.510198 | 0.00101 | 0.510386 | 13.6832 | 3.092346 | 0.035909 |
| **HCP-D** | rsfMRI | VII | -0.00697 | 0.00101 | -0.01359 | 14.19938 | 3.149851 | -0.00046 |
| **HCP-D** | rsfMRI | VI | 0.317007 | 0.00101 | 0.291792 | 14.01144 | 3.130864 | 0.012781 |
| **HCP-YA** | rsfMRI | CBL | 0.137727 | 0.001998 | 0.144947 | 13.44113 | 3.078751 | 0.010243 |
| **HCP-YA** | rsfMRI | DMN | 0.076339 | 0.002498 | 0.070545 | 13.53606 | 3.095075 | 0.003252 |
| **HCP-YA** | rsfMRI | FP | 0.137458 | 0.001998 | 0.132316 | 13.47496 | 3.091163 | 0.007751 |
| **HCP-YA** | rsfMRI | MF | 0.153549 | 0.001998 | 0.16041 | 13.46757 | 3.084167 | 0.008295 |
| **HCP-YA** | rsfMRI | Mot | 0.162647 | 0.001998 | 0.175725 | 13.3238 | 3.055389 | 0.018882 |
| **HCP-YA** | rsfMRI | SAL | 0.093017 | 0.002498 | 0.110685 | 13.51763 | 3.09266 | 0.004609 |
| **HCP-YA** | rsfMRI | SC | 0.102526 | 0.001998 | 0.106728 | 13.5164 | 3.091808 | 0.0047 |
| **HCP-YA** | rsfMRI | VAs | 0.070229 | 0.00444 | 0.097874 | 13.54053 | 3.094735 | 0.002923 |
| **HCP-YA** | rsfMRI | VII | -0.05925 | 0.073367 | -0.05402 | 13.60327 | 3.102696 | -0.0017 |
| **HCP-YA** | rsfMRI | VI | 0.106555 | 0.002498 | 0.127254 | 13.52213 | 3.091592 | 0.004278 |
| **HCP-A** | rsfMRI | CBL | -0.01229 | 0.007493 | -0.02425 | 176.2153 | 10.97947 | -0.00087 |
| **HCP-A** | rsfMRI | DMN | -0.04579 | 0.007493 | -0.02584 | 176.5047 | 10.99358 | -0.00251 |
| **HCP-A** | rsfMRI | FP | -0.07988 | 0.061605 | -0.05681 | 176.6841 | 10.99883 | -0.00353 |
| **HCP-A** | rsfMRI | MF | -0.00492 | 0.007493 | 0.00114 | 176.1699 | 10.96612 | -0.00061 |
| **HCP-A** | rsfMRI | Mot | 0.137267 | 0.007493 | 0.133694 | 174.7919 | 10.91558 | 0.007214 |

| | | | | | | | | |
|---|---|---|---|---|---|---|---|---|
| **HCP-A** | rsfMRI | SAL | -0.10119 | 0.108463 | -0.07502 | 176.8435 | 10.99513 | -0.00444 |
| **HCP-A** | rsfMRI | SC | -0.13005 | 0.23976 | -0.12563 | 177.0888 | 11.01941 | -0.00583 |
| **HCP-A** | rsfMRI | VAs | -0.07501 | 0.021978 | -0.08202 | 176.6884 | 10.99099 | -0.00356 |
| **HCP-A** | rsfMRI | VII | -0.15755 | 0.418933 | -0.15356 | 177.229 | 11.0046 | -0.00663 |
| **HCP-A** | rsfMRI | VI | -0.14535 | 0.29082 | -0.11268 | 177.1633 | 11.01342 | -0.00625 |
| **HCP-LS** | rsfMRI | CBL | 0.522453 | 0.001003 | 0.566678 | 245.1585 | 11.22656 | 0.015999 |
| **HCP-LS** | rsfMRI | DMN | 0.312829 | 0.001003 | 0.266831 | 247.5197 | 11.28794 | 0.006522 |
| **HCP-LS** | rsfMRI | FP | 0.638595 | 0.001003 | 0.643449 | 243.1222 | 11.17584 | 0.024172 |
| **HCP-LS** | rsfMRI | MF | 0.574674 | 0.001003 | 0.539779 | 244.4178 | 11.2132 | 0.018972 |
| **HCP-LS** | rsfMRI | Mot | 0.640717 | 0.001003 | 0.43895 | 239.4546 | 11.11953 | 0.038893 |
| **HCP-LS** | rsfMRI | SAL | 0.486297 | 0.001003 | 0.479985 | 244.5474 | 11.21813 | 0.018451 |
| **HCP-LS** | rsfMRI | SC | 0.445823 | 0.001003 | 0.435448 | 246.2915 | 11.26008 | 0.011451 |
| **HCP-LS** | rsfMRI | VAs | 0.170686 | 0.001003 | 0.18041 | 248.38 | 11.30453 | 0.003068 |
| **HCP-LS** | rsfMRI | VII | 0.002126 | 0.001003 | 0.022946 | 249.1537 | 11.31886 | -3.71E-05 |
| **HCP-LS** | rsfMRI | VI | 0.31011 | 0.001003 | 0.253986 | 247.5697 | 11.28492 | 0.006321 |

*Table S17. rCPM performance metrics from the median performing diffusion MRI (dMRI) QA End and resting-state fMRI (rsfMRI) between-network and within-network models for each age cohort (HCP-D, HCP-YA, HCP-A, and HCP-LS). Networks based on the 10 Shen networks. Model performance was evaluated with Pearson's correlation (r), Spearman's rank correlation (ρ), mean absolute error (MAE), mean squared error (MSE), and the predictive coefficient of determination ($q^2$). Network abbreviations: CBL = cerebellar, DMN = default mode network, FP = frontoparietal, MF = medial frontal, Mot = somatomotor, SAL = salience, SC = subcortical, VAs = visual association, VI = visual I, VII = visual II.*

**Sex-stratified rCPM age prediction**

**Commented [MM5]:** Need to finish nulls

| **Modality** | **Cohort** | **Sex** | **Pearson r** | **Null CPM FDR-corrected p** | **Spearman rho** | **MAE** | **MSE** | **$q^2$** |
|---|---|---|---|---|---|---|---|---|
| **dMRI QA End** | HCP-D | Males | 0.567756 | <0.05 | 0.571547 | 2.544144 | 9.78452 | 0.247821 |
| | | Females | 0.650162 | <0.05 | 0.664042 | 2.962843 | 12.20511 | 0.196099 |

| | | | | | | | | |
|---|---|---|---|---|---|---|---|---|
| | HCP-YA | Males | 0.131913 | <0.05 | 0.144707 | 2.976401 | 12.80464 | 0.016801 |
| | | Females | 0.143793 | <0.05 | 0.134855 | 2.957415 | 12.6504 | 0.020545 |
| | HCP-A | Males | 0.691508 | <0.05 | 0.678035 | 10.42856 | 155.7693 | 0.171963 |
| | | Females | 0.610931 | <0.05 | 0.556478 | 9.715866 | 142.3562 | 0.142151 |
| | HCP-LS | Males | 0.74871 | <0.05 | 0.445126 | 9.520398 | 178.9854 | 0.258148 |
| | | Females | 0.697981 | <0.05 | 0.550147 | 10.4703 | 201.857 | 0.208582 |
| **rsfMRI** | HCP-D | Males | 0.802447 | <0.05 | 0.816917 | 1.915133 | 5.724575 | 0.559927 |
| | | Females | 0.787644 | <0.05 | 0.806126 | 2.135613 | 6.671953 | 0.560546 |
| | HCP-YA | Males | 0.283683 | <0.05 | 0.301184 | 2.886439 | 12.01293 | 0.077592 |
| | | Females | 0.274338 | <0.05 | 0.290834 | 2.838311 | 11.95088 | 0.074706 |
| | HCP-A | Males | 0.493775 | <0.05 | 0.45447 | 10.82699 | 166.3684 | 0.11562 |
| | | Females | 0.433404 | <0.05 | 0.400697 | 9.974116 | 151.6503 | 0.086144 |
| | HCP-LS | Males | 0.824949 | <0.05 | 0.765169 | 8.217749 | 139.9886 | 0.41978 |
| | | Females | 0.822798 | <0.05 | 0.818272 | 8.872306 | 151.1019 | 0.407577 |

*Table S18. rCPM performance metrics from the median performing dMRI QA End and rsfMRI male-only and female-only models for each age cohort (HCP-D, HCP-YA, HCP-A, and HCP-LS). Model performance was evaluated with Pearson's correlation (r), Spearman's rank correlation (ρ), mean absolute error (MAE), mean squared error (MSE), and the predictive coefficient of determination ($q^2$).*

**Comparison of male and female rCPM performance metrics**

| Cohort | Modality | Male r | Female r | Male N | Female N | Fisher Z | P | FDR |
|---|---|---|---|---|---|---|---|---|
| **HCP_A** | dMRI QA End | 0.691508 | 0.610931 | 119 | 171 | 1.16333 | 0.244696 | 0.587269 |
| **HCP_A** | rsfMRI | 0.493775 | 0.433404 | 119 | 171 | 0.637507 | 0.523794 | 0.898262 |
| **HCP_D** | dMRI QA End | 0.567756 | 0.650162 | 210 | 248 | -1.39158 | 0.164051 | 0.587269 |
| **HCP_D** | rsfMRI | 0.802447 | 0.787644 | 210 | 248 | 0.426372 | 0.669837 | 0.898262 |
| **HCP_LS** | dMRI QA End | 0.74871 | 0.697981 | 684 | 787 | 2.03619 | 0.041731 | 0.500775 |
| **HCP_LS** | rsfMRI | 0.824949 | 0.822798 | 684 | 787 | 0.127857 | 0.898262 | 0.898262 |
| **HCP_YA** | dMRI QA End | 0.131913 | 0.143793 | 355 | 368 | -0.1621 | 0.871226 | 0.898262 |
| **HCP_YA** | rsfMRI | 0.283683 | 0.274338 | 355 | 368 | 0.135642 | 0.892104 | 0.898262 |

*Table S19. Comparison of Pearson r correlations from the median performing Male-only and Female-only rCPM models using Fisher r-to-z transformation.*

| Cohort | Modality | Train Sex | Test Sex | Pearson r | p-value | FDR-corrected p | Spearman rho | MSE | MAE | q |
|---|---|---|---|---|---|---|---|---|---|---|
| HCP-D | dMRI QA End | F | M | 0.752493 | 1.37E-39 | 3.30E-39 | 0.759509 | 9.829126 | 2.571786 | 0.244392 |
| HCP-D | dMRI QA End | M | F | 0.472759 | 3.26E-15 | 4.89E-15 | 0.476444 | 12.54474 | 2.935223 | 0.173729 |
| HCP-D | rsfMRI | F | M | 0.78412 | 5.66E-45 | 1.70E-44 | 0.796946 | 5.927785 | 1.904936 | 0.544305 |
| HCP-D | rsfMRI | M | F | 0.789076 | 5.61E-54 | 1.92E-53 | 0.804417 | 7.406772 | 2.283369 | 0.512146 |
| HCP-YA | dMRI QA End | F | M | 0.212191 | 5.58E-05 | 6.38E-05 | 0.194269 | 15.17368 | 3.313125 | -0.1651 |
| HCP-YA | dMRI QA End | M | F | 0.160765 | 0.001977 | 0.002063 | 0.145441 | 15.46567 | 3.278256 | -0.19743 |
| HCP-YA | rsfMRI | F | M | 0.156058 | 0.003198 | 0.003198 | 0.178775 | 15.173 | 3.267411 | -0.16505 |
| HCP-YA | rsfMRI | M | F | 0.1955 | 0.000161 | 0.000175 | 0.213046 | 15.5792 | 3.300215 | -0.20622 |
| HCP-A | dMRI QA End | F | M | 0.759663 | 1.32E-23 | 2.63E-23 | 0.739933 | 156.531 | 10.45602 | 0.167914 |
| HCP-A | dMRI QA End | M | F | 0.623896 | 7.83E-20 | 1.25E-19 | 0.557701 | 140.2499 | 9.727861 | 0.154844 |
| HCP-A | rsfMRI | F | M | 0.566766 | 1.83E-11 | 2.58E-11 | 0.576645 | 172.975 | 10.97508 | 0.080501 |
| HCP-A | rsfMRI | M | F | 0.4309 | 4.03E-09 | 5.37E-09 | 0.414763 | 153.249 | 10.14453 | 0.076511 |
| HCP-LS | dMRI QA End | F | M | 0.742398 | ######## | ######## | 0.439109 | 186.3995 | 9.840841 | 0.227418 |
| HCP-LS | dMRI QA End | M | F | 0.704116 | ######## | ######## | 0.557728 | 196.1267 | 10.25649 | 0.231048 |
| HCP-LS | rsfMRI | F | M | 0.820609 | ######## | ######## | 0.789838 | 144.9105 | 8.975071 | 0.39938 |
| HCP-LS | rsfMRI | M | F | 0.78742 | ######## | ######## | 0.773374 | 172.0511 | 9.184007 | 0.325442 |

*Table S20. Cross-sex rCPM model generalizability using final rCPM models.*

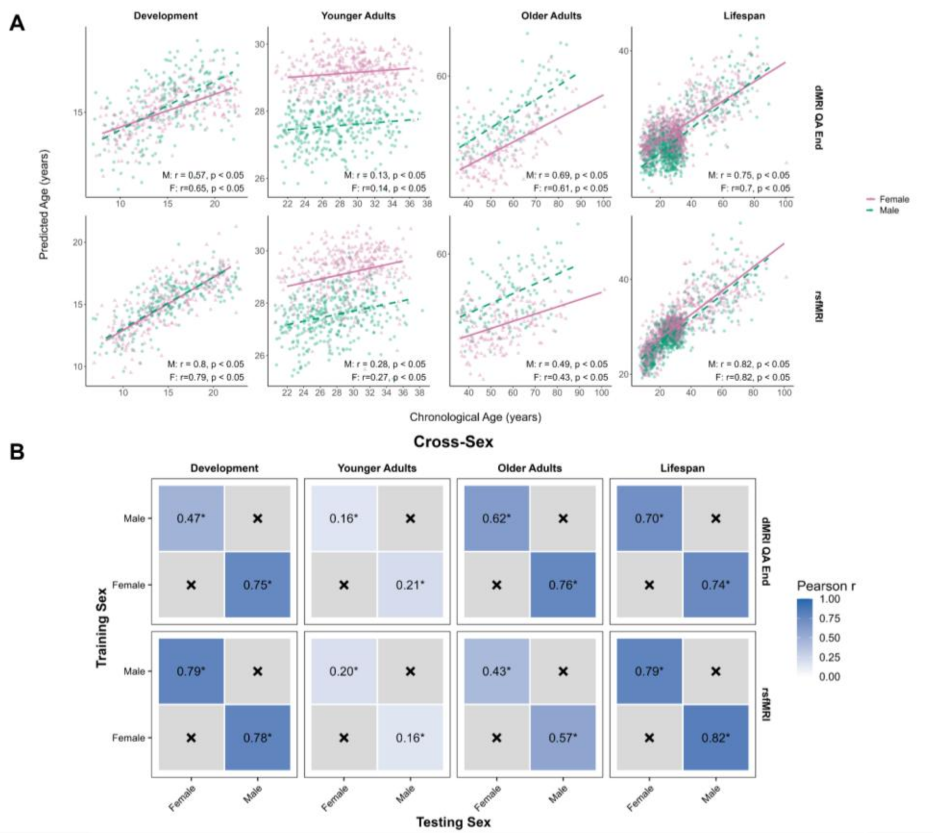

*Figure S4. (A) Sex-stratified rCPM age prediction performance. Predicted age plotted against chronological age for male-only and female-only rCPM models across age cohorts and imaging modalities. Male-only and female-only models were trained separately within each cohort, and sex-specific regression lines were fitted to visualize prediction accuracy, age-related prediction bias, and the relationship between chronological and predicted age. Male models are shown in green and female models are shown in pink. (B) Heatmaps summarize the generalizability of final rCPM age prediction models evaluated through three forms of external validation/cross-prediction. Heatmap values represent the Pearson correlation coefficient (r) between predicted and chronological age when models were applied to independent datasets. Darker shading indicates stronger age prediction performance (higher Pearson r). Asterisks indicate predictions that remained significant after false discovery rate (FDR) correction ($p < 0.05$). Gray cells marked with × indicate comparisons that were not performed (e.g., identical training and testing sex).*

**Multimodal rCPM**

| Multimodal rCPM |
| --- |

| Age Cohort | dMRI QA End Total Edges | rsfMRI Total Edges |
|---|---|---|
| **HCP-Development** | 8,433 | 22,157 |
| **HCP-Young Adults** | 5,751 | 10,017 |
| **HCP-Aging** | 8,593 | 11,577 |
| **HCP-Lifespan** | 15,437 | 26,977 |

*Table S21. Total edge density from dMRI QA End and rsfMRI components from the median performing multimodal rCPM coefficient predictive networks.*

#### Multimodal vs unimodal rCPM performance

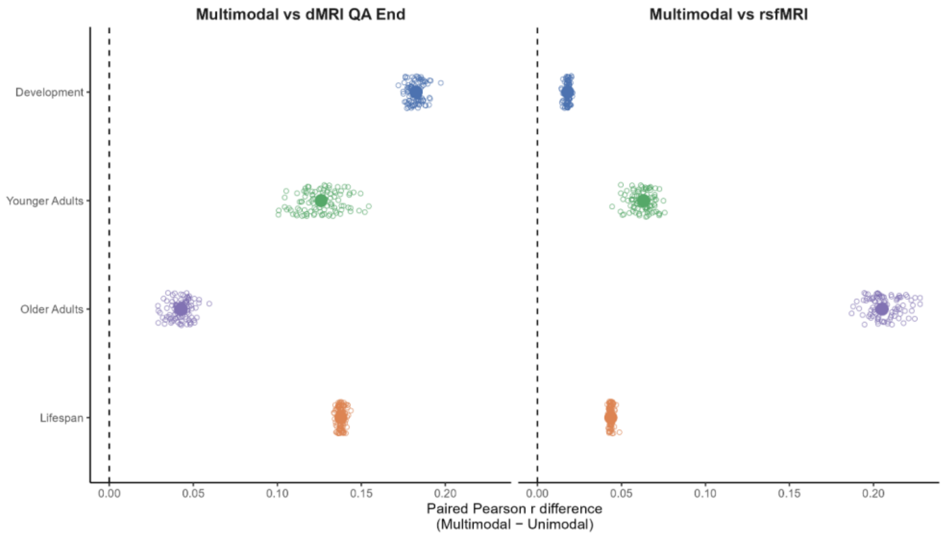

*Figure S7. Paired comparison of multimodal and unimodal rCPM age prediction performance using dMRI QA End and rsfMRI connectomes. Distribution of paired iteration-wise differences in Pearson correlation coefficients between multimodal rCPM combining dMRI QA End and rsfMRI connectomes and corresponding unimodal rCPM models across HCP cohorts. Each point represents one of 100 paired cross-validation iterations, and larger points indicate the median difference across iterations. Positive values indicate higher prediction performance for the multimodal model, whereas negative values indicate higher performance for the corresponding unimodal model. Comparisons are shown for multimodal rCPM versus dMRI QA End alone (left) and multimodal rCPM versus rsfMRI alone (right) across Development (HCP-D), Young Adult (HCP-YA), Aging (HCP-A), and Lifespan (HCP-LS) cohorts.*

| Cohort | Comparison | Metric | Iterations Favoring Multimodal | Iterations Favoring Unimodal | % Iterations Multimodal Better | p-value | FDR-corrected p |
|---|---|---|---|---|---|---|---|
| **HCP_A** | multimodal vs dMRI QA End | mae | 100 | 0 | 100 | 7.89E-31 | 7.89E-31 |
| **HCP_A** | multimodal vs dMRI QA End | mse | 100 | 0 | 100 | 7.89E-31 | 7.89E-31 |

| | | | | | | | |
|---|---|---|---|---|---|---|---|
| **HCP_A** | multimodal vs dMRI QA End | pearson_r | 100 | 0 | 100 | 7.89E-31 | 7.89E-31 |
| **HCP_D** | multimodal vs dMRI QA End | mae | 100 | 0 | 100 | 7.89E-31 | 7.89E-31 |
| **HCP_D** | multimodal vs dMRI QA End | mse | 100 | 0 | 100 | 7.89E-31 | 7.89E-31 |
| **HCP_D** | multimodal vs dMRI QA End | pearson_r | 100 | 0 | 100 | 7.89E-31 | 7.89E-31 |
| **HCP_LS** | multimodal vs dMRI QA End | mae | 100 | 0 | 100 | 7.89E-31 | 7.89E-31 |
| **HCP_LS** | multimodal vs dMRI QA End | mse | 100 | 0 | 100 | 7.89E-31 | 7.89E-31 |
| **HCP_LS** | multimodal vs dMRI QA End | pearson_r | 100 | 0 | 100 | 7.89E-31 | 7.89E-31 |
| **HCP_YA** | multimodal vs dMRI QA End | mae | 100 | 0 | 100 | 7.89E-31 | 7.89E-31 |
| **HCP_YA** | multimodal vs dMRI QA End | mse | 100 | 0 | 100 | 7.89E-31 | 7.89E-31 |
| **HCP_YA** | multimodal vs dMRI QA End | pearson_r | 100 | 0 | 100 | 7.89E-31 | 7.89E-31 |
| **HCP_A** | multimodal vs rsfMRI | mae | 100 | 0 | 100 | 7.89E-31 | 7.89E-31 |
| **HCP_A** | multimodal vs rsfMRI | mse | 100 | 0 | 100 | 7.89E-31 | 7.89E-31 |
| **HCP_A** | multimodal vs rsfMRI | pearson_r | 100 | 0 | 100 | 7.89E-31 | 7.89E-31 |
| **HCP_D** | multimodal vs rsfMRI | mae | 100 | 0 | 100 | 7.89E-31 | 7.89E-31 |
| **HCP_D** | multimodal vs rsfMRI | mse | 100 | 0 | 100 | 7.89E-31 | 7.89E-31 |
| **HCP_D** | multimodal vs rsfMRI | pearson_r | 100 | 0 | 100 | 7.89E-31 | 7.89E-31 |
| **HCP_LS** | multimodal vs rsfMRI | mae | 100 | 0 | 100 | 7.89E-31 | 7.89E-31 |

| | | | | | | | |
|---|---|---|---|---|---|---|---|
| **HCP_LS** | multimodal vs rsfMRI | mse | 100 | 0 | 100 | 7.89E-31 | 7.89E-31 |
| **HCP_LS** | multimodal vs rsfMRI | pearson_r | 100 | 0 | 100 | 7.89E-31 | 7.89E-31 |
| **HCP_YA** | multimodal vs rsfMRI | mae | 100 | 0 | 100 | 7.89E-31 | 7.89E-31 |
| **HCP_YA** | multimodal vs rsfMRI | mse | 100 | 0 | 100 | 7.89E-31 | 7.89E-31 |
| **HCP_YA** | multimodal vs rsfMRI | pearson_r | 100 | 0 | 100 | 7.89E-31 | 7.89E-31 |

*Table S22. Paired iteration-wise comparisons of multimodal and unimodal rCPM performance across cohorts.*

**Multimodal-unimodal rCPM coefficient similarity**

| **Edge-level** | | | | | |
|---|---|---|---|---|---|
| **Comparison** | Cohort | Type | Edge Correlation r | Permutation p | Permutation p-FDR |
| **DTI_MultiQAEND** | HCP-D | Signed | 0.944675 | 1.00E-04 | 1.00E-04 |
| **DTI_MultiQAEND** | HCP-D | Absolute | 0.91608 | 1.00E-04 | 1.00E-04 |
| **DTI_MultiQAEND** | HCP-YA | Signed | 0.981898 | 1.00E-04 | 1.00E-04 |
| **DTI_MultiQAEND** | HCP-YA | Absolute | 0.976126 | 1.00E-04 | 1.00E-04 |
| **DTI_MultiQAEND** | HCP-A | Signed | 0.995224 | 1.00E-04 | 1.00E-04 |
| **DTI_MultiQAEND** | HCP-A | Absolute | 0.992746 | 1.00E-04 | 1.00E-04 |
| **DTI_MultiQAEND** | HCP-LS | Signed | 0.994462 | 1.00E-04 | 1.00E-04 |
| **DTI_MultiQAEND** | HCP-LS | Absolute | 0.99015 | 1.00E-04 | 1.00E-04 |
| **REST_MultiQAEND** | HCP-D | Signed | 0.971031 | 1.00E-04 | 1.00E-04 |
| **REST_MultiQAEND** | HCP-D | Absolute | 0.941518 | 1.00E-04 | 1.00E-04 |
| **REST_MultiQAEND** | HCP-YA | Signed | 0.978667 | 1.00E-04 | 1.00E-04 |
| **REST_MultiQAEND** | HCP-YA | Absolute | 0.963809 | 1.00E-04 | 1.00E-04 |
| **REST_MultiQAEND** | HCP-A | Signed | 0.979181 | 1.00E-04 | 1.00E-04 |
| **REST_MultiQAEND** | HCP-A | Absolute | 0.958498 | 1.00E-04 | 1.00E-04 |
| **REST_MultiQAEND** | HCP-LS | Signed | 0.9941 | 1.00E-04 | 1.00E-04 |
| **REST_MultiQAEND** | HCP-LS | Absolute | 0.983699 | 1.00E-04 | 1.00E-04 |
| **Node-level** | | | | | |
| **Comparison** | Cohort | Type | Node Pearson r | Permutation p | Permutation p-FDR |
| **DTI_MultiQAEND** | HCP-D | Absolute | 0.98436 | 1.00E-04 | 1.00E-04 |
| **DTI_MultiQAEND** | HCP-YA | Absolute | 0.988542 | 1.00E-04 | 1.00E-04 |
| **DTI_MultiQAEND** | HCP-A | Absolute | 0.999092 | 1.00E-04 | 1.00E-04 |
| **DTI_MultiQAEND** | HCP-LS | Absolute | 0.998205 | 1.00E-04 | 1.00E-04 |
| **REST_MultiQAEND** | HCP-D | Absolute | 0.989472 | 1.00E-04 | 1.00E-04 |

| | | | | | |
|---|---|---|---|---|---|
| **REST_MultiQAEND** | HCP-YA | Absolute | 0.9905 | 1.00E-04 | 1.00E-04 |
| **REST_MultiQAEND** | HCP-A | Absolute | 0.991686 | 1.00E-04 | 1.00E-04 |
| **REST_MultiQAEND** | HCP-LS | Absolute | 0.992442 | 1.00E-04 | 1.00E-04 |
| | | | **Network-level** | | |
| **Comparison** | Cohort | Type | Network Contribution r | Permutation p | Permutation p-FDR |
| **DTI_MultiQAEND** | HCP-D | Signed | 0.953303 | 1.00E-04 | 1.00E-04 |
| **DTI_MultiQAEND** | HCP-D | Absolute | 0.958921 | 1.00E-04 | 1.00E-04 |
| **DTI_MultiQAEND** | HCP-YA | Signed | 0.991067 | 1.00E-04 | 1.00E-04 |
| **DTI_MultiQAEND** | HCP-YA | Absolute | 0.990178 | 1.00E-04 | 1.00E-04 |
| **DTI_MultiQAEND** | HCP-A | Signed | 0.998956 | 1.00E-04 | 1.00E-04 |
| **DTI_MultiQAEND** | HCP-A | Absolute | 0.999108 | 1.00E-04 | 1.00E-04 |
| **DTI_MultiQAEND** | HCP-LS | Signed | 0.998638 | 1.00E-04 | 1.00E-04 |
| **DTI_MultiQAEND** | HCP-LS | Absolute | 0.998072 | 1.00E-04 | 1.00E-04 |
| **REST_MultiQAEND** | HCP-D | Signed | 0.974605 | 1.00E-04 | 1.00E-04 |
| **REST_MultiQAEND** | HCP-D | Absolute | 0.995258 | 1.00E-04 | 1.00E-04 |
| **REST_MultiQAEND** | HCP-YA | Signed | 0.987211 | 1.00E-04 | 1.00E-04 |
| **REST_MultiQAEND** | HCP-YA | Absolute | 0.972808 | 1.00E-04 | 1.00E-04 |
| **REST_MultiQAEND** | HCP-A | Signed | 0.988146 | 1.00E-04 | 1.00E-04 |
| **REST_MultiQAEND** | HCP-A | Absolute | 0.985079 | 1.00E-04 | 1.00E-04 |
| **REST_MultiQAEND** | HCP-LS | Signed | 0.994338 | 1.00E-04 | 1.00E-04 |
| **REST_MultiQAEND** | HCP-LS | Absolute | 0.994955 | 1.00E-04 | 1.00E-04 |

*Table S23. rCPM multimodal-unimodal similarity analysis at the edge, node, and network levels.*
